# Functional Characterization of Arabidopsis α-Amylase3 (AMY3): Amylose Specificity and Structural Insights into Its Duplex Carbohydrate-Binding Module

**DOI:** 10.1101/2025.09.19.677355

**Authors:** Ruhi Rahman, Sara E. Scanlan, Fady Sidarous, Jonathan D. Monroe, Christopher E. Berndsen

## Abstract

Flowering plants contain a plastid-localized ɑ-amylase (AMY3) with several recognized domains: two N-terminal carbohydrate-binding modules, an alpha-alpha hairpin that binds the catalytically inactive activator β-amylase9, and a C-terminal catalytic domain. Despite considerable research on leaf starch metabolism in Arabidopsis, little is known about the function of AMY3. Starch is composed of two glucose polymers: highly branched amylopectin (70-90%) and largely unbranched amylose (10-30%). Because ɑ-amylases are endo-amylases, we predicted that AMY3 might prefer to hydrolyze amylose, in contrast to the exo-amylase β-amylase1 (BAM1), which we predicted would prefer amylopectin. Using iodine spectroscopy, we observed that AMY3 caused the percent amylose of corn starch to decrease, whereas BAM1 caused the percent amylose to increase before a gradual decrease. Enzyme assays revealed that AMY3 has a higher affinity for amylose compared to amylopectin, whereas BAM1 has a higher affinity for amylopectin. Furthermore, we found that the starch in *amy3* leaves had twice as much amylose as WT leaves at the end of the day which suggests potential health and industrial applications. We purified the AMY3 CBM region and studied its structure with small-angle X-ray scattering, seeking to understand its role in substrate specificity. The AMY3 CBMs form a unique duplex where two interlocking β-strands connect CBM1 and CBM2. The duplex CBM forms a dimer in solution and can bind amylose and amylopectin. However, AMY3 without the CBMs still prefers amylose, indicating the duplex CBM is not essential for amylose specificity *in vitro*. However, the duplex CBM is important for AMY3 dimerization.

## INTRODUCTION

Plants carry out photosynthesis to generate reduced carbon in the form of sugars for use in biosynthesis and as a source of energy. Because light is only available for a portion of each day, the rate of photosynthesis is higher than that needed for daytime use, and much of the photosynthate generated is stored in plastids, usually as starch (1). At night or during periods of stress when reduced carbon demands exceed supply, starch is broken down to maintain metabolism (2, 3). Storing glucose as starch is advantageous because starch isn’t soluble in water, so the large starch granules don’t disrupt the cell’s osmotic balance. Starch granules are composed of two types of glucose polymers: amylopectin and amylose. Both polymers contain long chains of ɑ-1,4-linked glucose, but only amylopectin, comprising about 70-90% of most starch granules, contains ɑ-1,6 branches that enable the polymer to form massive semi-crystalline granules (2). The role of the amylose fraction in starch is not understood; mutants lacking the enzyme responsible for amylose synthesis, GRANULE-BOUND STARCH SYNTHASE (GBSS), and even some naturally occurring plants lacking amylose have no obvious phenotype (4).

The pathway used for leaf starch degradation at night is relatively well understood and has been recently reviewed (5). To initiate enzymatic attack, the outer chains of amylopectin are first phosphorylated by GLUCAN WATER DIKINASE1 (GWD1) and PHOSPHOGLUCAN WATER DIKINASE (PWD) (6, 7). This modification causes hydration of the outer chains (8), facilitating their hydrolysis primarily by β-AMYLASE3 (BAM3), and partially by BAM1 (9, 10). For hydrolysis to proceed, the phosphates are removed by STARCH EXCESS4 (SEX4) and LIKE SEX FOUR2 (LSF2) (11, 12). The ɑ-1,6 bonds are removed by ISOAMYLASE3 (ISA3) and LIMIT DEXTRINASE (LD) (7, 13). Despite advances in understanding starch degradation, details about the specific roles of some enzymes are lacking, especially when there are multiple forms of an enzyme (e.g., β-amylase) (14, 15). How the overall process of starch degradation is regulated is also largely unknown.

One enzyme that, based on its catalytic activity and plastid localization, is predicted to play a role in leaf starch degradation is α-AMYLASE3 (AMY3), however, no specific role for this enzyme has been described. Unlike mutants of Arabidopsis that accumulate starch in leaves if they lack essential proteins involved in starch degradation, AMY3 mutants show no diurnal leaf starch accumulation phenotype (16). AMY3, along with BAM1, is thought to contribute to starch degradation in guard cells at dawn (17) or in response to osmotic stress (18). Importantly, AMY3 is active only in the presence of reducing agents and is thus predicted to be active only during the day when reducing levels in the chloroplast are high, which seems contradictory to its role in starch degradation (19). Seung *et al.* (19) used mutagenesis to suggest that a disulfide bridge between Cys499 and Cys587, both in the catalytic domain, regulates its activity. In addition to the catalytic amylase domain, AMY3 contains two N-terminal carbohydrate-binding modules (CBM), and an alpha-alpha hairpin (AAH), both of which are unique amongst amylases to AMY3 and its orthologs (20–22). We recently showed that AMY3 activity is elevated up to 3-fold by binding to catalytically inactive BAM9, which is predicted to bind to the conserved AAH structure of AMY3 (20). Moreover, in the absence of BAM9, the reduced form of AMY3 forms a homodimer, and in the presence of BAM9, the homodimer is replaced by a heterodimer of AMY3 and BAM9 subunits (20). What role this dimeric interaction plays in the function of the enzyme is difficult to predict without a known substrate for AMY3. Glaring et al. demonstrated that the CBMs of AMY3 and their related CAZy family 45 counterparts in GWD exhibit weak starch binding, yet the structure and function of AMY3’s CBMs remain uncharacterized (21).

The primary catalytic difference between an α-amylase and a β-amylase is that the former is an endo-amylase, cleaving internal α-1,4 linkages in starch, whereas the latter is an exo-amylase, cleaving the penultimate α-1,4 linkages at the non-reducing ends of starch, releasing maltose (23). Exploring the older literature revealed an awareness of these differences as early as the late 1800s, even before there was an understanding of the nature of proteins or starch (24–26). Experiments on germinating barley revealed two types of enzymes that had different effects on starch; one was a liquifying enzyme that decreased the viscosity of a starch solution without generating much reducing power (27). Later, this enzyme was understood to be α-amylase (28). The other enzyme was a saccharifying enzyme that generated reducing sugars without affecting the substrate’s viscosity, and this enzyme was later understood to be β-amylase (29).

Differences between the amylases and our current understanding of the composition of starch led us to speculate that the two types of amylases might prefer the different polymers of starch: the exo-amylase (BAMs) preferring amylopectin because of its many reducing ends, and the endo-amylase (AMYs) preferring amylose with its few ends and long unbranched regions. To this end, we describe *in vitro* and *in vivo* experiments revealing that AMY3 has a preference for amylose as its substrate and provide evidence for how each domain plays a role in defining this specificity. We suggest that one function of AMY3 is to counteract the activity of GBSS in amylose synthesis during daytime starch synthesis (4, 30–33). Surprisingly, amylose preference is encoded largely by the amylase domain. Furthermore, we find that AMY3 dimerization, and therefore its activity, is linked to the dimerization of the CBMs, a previously undescribed function.

## METHODS

### Bioinformatics

Reference amino acid sequences of AMY3 and GWD1 were obtained from NCBI using the Arabidopsis sequences as the query in BLASTp searches (https://blast.ncbi.nlm.nih.gov/Blast.cgi). The 20 taxa used represented 20 different orders of flowering plants to minimize high identity due to closely related species. The sequences were aligned using Clustal Omega (34, 35) and visualized using Boxshade (https://junli.netlify.app/apps/boxshade/). The list of species and accession numbers is in Supplemental Table 1, and the sequence alignments are in Supplemental Figures 1 and 2. For CBM analysis, sequences of each CBM domain were isolated manually and aligned to generate pairwise percent identities.

### Protein Purifications

Residues 56-391 of Arabidopsis AMY3 (At1g69830, NP_564977.1, Uniprot ID: Q94A41) which included both CBMs of Arabidopsis AMY3 was cloned into pET21a by Genscript. Cells expressing CBM were grown in 2xYT and 0.06 mg/mL ampicillin to an OD600 of approximately 0.6 at 37°C and 225 rpm, then induced with 1 mM Isopropyl-β-D-thiogalactopyranoside (IPTG) at 20°C overnight. Cells were lysed in 50 mM Tris, pH 8.0, 100 mM sodium chloride, 0.1 mM EDTA, and 1 mM imidazole using sonication at 65% power for 5 s on and 5 s off for 5 min. Cell debris was removed by centrifugation at 17,000 × g at 4°C. The supernatant was then passed over 2 mL of Ni-NTA resin, and the column was allowed to drain by gravity. The column was washed with 20 mL of Wash Buffer (50 mM Tris, pH 8.0, 300 mM sodium chloride, 0.1 mM EDTA, 10 mM Imidazole). Protein was eluted with 10 mL steps from 9:1 wash buffer: elution buffer (50 mM Tris, pH 8.0, 50 mM sodium chloride, 0.1 mM EDTA, 300 mM Imidazole) to 0:10 wash buffer: elution buffer. The presence of protein was confirmed via SDS-PAGE, combined elutions were concentrated to 2 mL, and then injected onto a Sephacryl-200 16/60 column in 50 mM sodium phosphate, pH 7, 100 mM sodium chloride, and 5 mM DTT. The presence of protein in the fractions was confirmed via SDS-PAGE. Protein-containing fractions were pooled, concentrated, and then stored at −80°C. Purification of the C-terminal catalytic domain (residues 484-837 referred to as AMY3_cat_) was as previously described (20). BAM1 was purified as previously described (36).

### CBM Crosslinking

For crosslinking assays, CBM was diluted to 20 μM in 20 mM HEPES, pH 7.5 prior in 25 μL to the addition of an equal volume of 0.05% glutaraldehyde. The crosslinking reaction was kept at room temperature (∼22°C) for 30 min before being quenched with 30 μL of 1 M Tris base. Glutaraldehyde and Tris were allowed to react for 5 min before the reaction was mixed with 2x SDS-PAGE loading dye. Proteins were then separated via SDS-PAGE.

### Small-Angle X-Ray Scattering (SAXS)

Purified CBM was dialyzed into 100 mM potassium phosphate at pH 7.4. Purified protein samples of CBM alone and cross-linked CBM were shipped overnight on dry ice and analyzed using SEC-SAXS at the SIBYLS beamline at the Advanced Light Source, Lawrence Berkeley National Laboratory, Berkeley, CA (37). Samples were separated on a Shodex K-W-803 column at a flow rate of 0.5 mL/min at 10 °C, and the eluate was measured in line with UV/vis absorbance at 280 nm, multi-angle X-ray scattering (MALS), and SAXS. The incident light wavelength was 1.127 Å at a sample-to-detector distance of 2.1 m. This setup results in scattering vectors, q, ranging from 0.0114 to 0.4 Å^−1^, where the scattering vector is defined as q = 4πsin(θ)/λ, with θ being the measured scattering angle.

AMY3_cat_ samples were similarly dialyzed into 100 mM potassium phosphate, pH 7.4, but were collected via HT-SAXS and SEC-SAXS (37, 38). For HT-SAXS, AMY3_cat_ was diluted in dialysis buffer over a concentration range of 0.3 to 1 mg/mL. For HT-SAXS, data frames were collected every 0.3 seconds for 50 frames of data. SEC-SAXS data were collected as above for CBM with 60 μL of 1 mg/mL AMY3_cat_.

Radially averaged SAXS data files were processed and analyzed in RAW (39). The radius of gyration (Rg) was calculated for each of the subtracted frames using the Guinier approximation: I(q) = I(0) exp(−q2Rg2/3) with the limits qRg < 1.3. The elution peak was compared to the integral of the ratios to background and Rg relative to the recorded frame using the RAW program. Uniform Rg values across an elution peak represent a homogeneous sample. The final merged SAXS profiles, derived by integrating multiple frames at the elution peak, were used for further analysis. We calculated the Guinier plot to provide information on the aggregation state and the pair distribution function [P(r)] to calculate the maximal inter-particle dimension. Models were fitted to the SAXS data using FOXS, and for CBM models, refined using BilboMD (40, 41). All SAXS data are available through the SASBDB under codes SASDX79 and SASDX99 for the AMY3 CBM and SASDX89 for AMY3_cat_ (42).

### Starch Binding Assay: CBM in different pHs

Starches [corn amylopectin (Sigma #A7780), soluble starch (Acros #424495000), wheat starch (Sigma S5127), corn starch (Sigma #S4126), and pea starch (Big Green Inc., #678452140751)] were blocked in 1% BSA with shaking for 30 min at 24°C, then the BSA was removed by three washes with dH_2_O followed by centrifugation at 4000g for 10 min at 24°C and resuspension in dH_2_O. 200 mg of starch was then transferred to a microcentrifuge tube and pelleted with centrifugation. To the starch pellets, 50 μL of buffer (50 mM MES/acetate/TRIS, pH 7 or as described in the experiment) containing 20 mM DTT and 5 mg protein were added, and pellets were vortexed and then shaken at 4°C for 60 min. Tubes were centrifuged for 2 min at 15,000g then 40 μL of the supernatant was removed and combined with 20 μL of 4X SDS loading buffer containing β-mercaptoethanol. The starch pellet was then washed three times with 0.5 mL of the same buffer, followed by vortexing and centrifugation for 2 min each time. After removing the supernatant of the last wash, 50 μL of 2X SDS loading buffer was added to elute proteins bound to the starch pellet. After one last centrifugation, 40 μL of the supernatant was removed for analysis. Samples were then boiled for 5 min, and proteins were separated by SDS-PAGE. For amylose competition assays, the same procedure was followed using insoluble amylopectin, but various amounts of soluble amylose were included with the protein in the binding step. Potato amylose (Sigma #A0512) was dissolved in 2 N NaOH, neutralized with 2 N HCl, precipitated in 70% EtOH, centrifuged at 4000g for 10 min at 24 °C, and then the pellet was dissolved in dH_2_O.

### Enzyme Assays and Starch Analysis

Assays with purified enzymes were conducted in 0.5 mL containing 50 mM MOPS, pH 7.0, 100 mM KCl, 1 mM MgCl_2_, 5 mM DTT, and various concentrations of specified substrate. Corn amylopectin and corn starch were heated to 100°C in the assay buffer and then cooled to 25°C and used immediately. Potato amylose (Sigma A0512) was dissolved in 2 N NaOH, neutralized with an equal volume of 2 N HCl, then diluted with the assay buffer. Enzymes were diluted with 50 mM MOPS and 1 mg mL^-1^ BSA prior to initiating the assays, which were conducted at 25°C for 20-45 min and then stopped in a boiling water bath. Reducing sugars were measured using the Somogyi-Nelson assay (43). For percent amylose measurements, 4 μL of assay solution was removed at designated times and combined with 200 μL of 10% Lugol’s iodine [0.34% (w/v) I_2_ and 0.68% (w/v) KI], mixed by pipetting and then the A_525_ and A_700_ were measured immediately. Percent amylose was then calculated by the method in (44).

Percent amylose in extracted leaf starch was measured using the protocol of Hovenkamp-Hermelink et al. (45). Arabidopsis leaves from plants grown as described in Monroe et al. were harvested at the end of the light period and processed immediately (46). The *amy3* T-DNA line (SALK_048475) was obtained from the Arabidopsis Biological Resource Center (Ohio State University) and confirmed using the PCR primers: 5’`-`TACTGTCCACTGGGGAGTTTG-3’ and 5’-TCCCAGATTTATTGGACTCCC-3’. Approximately 200 mg of leaves from at least 3 age-matched plants per replicate were sliced into 1-2 mm strips, submerged in 2 mL of 45% perchloric acid, shaken gently for 15 min, then 10 mL of water was added and mixed. To 180 μL of this extract, 20 μL of Lugol’s iodine was added, and the percent amylose was measured as described above.

### Circular Dynamics (CD) Spectroscopy

For CD measurements, the cuvette contained 7 μM CBM in 20 mM HEPES, 50 mM potassium chloride, and at pH 7.0 in a 0.2 cm quartz cuvette. Data were collected from 300 nm to 200 nm at 1 nm/sec with three accumulations on a Jasco J-1500 CD spectrometer. The predicted CD spectrum of a refined SAXS-based model of CBM was generated using PDBMD2CD to calculate the percent helix, sheet, and coil, and the data were overlaid with the predicted spectrum in R (47).

## RESULTS

Inspired by the notion that α-amylase, an endo-amylase, might have a preference for hydrolyzing amylose due to its long regions of unbranched α-1,4-linked glucose and few non-reducing ends, we tested this idea by observing the effect of purified AMY3 and a β-amylase, BAM1, on the percent amylose of corn starch containing both amylose and amylopectin. We predicted that AMY3 would cause the percent amylose to decrease, and BAM1, an exo-amylase that acts on non-reducing ends that are abundant in highly branched amylopectin, would cause the percent amylose in a corn starch to increase. Before digestion, the corn starch contained about 30% amylose. As predicted, BAM1 caused the percentage of amylose to increase to approximately 40% before declining, suggesting that it hydrolyzed the amylopectin fraction preferentially but could hydrolyze both polymers (Figure 1A). In contrast, AMY3 caused the percent amylose to decrease to zero, suggesting that it preferred amylose (Figure 1A). We tested the AMY3 catalytic domain (AMY3_cat_) and found it also reduced the percent amylose to zero, showing the catalytic domain partly determines AMY3’s preference for amylose (Figure 1A) (20).

**Figure 1.**
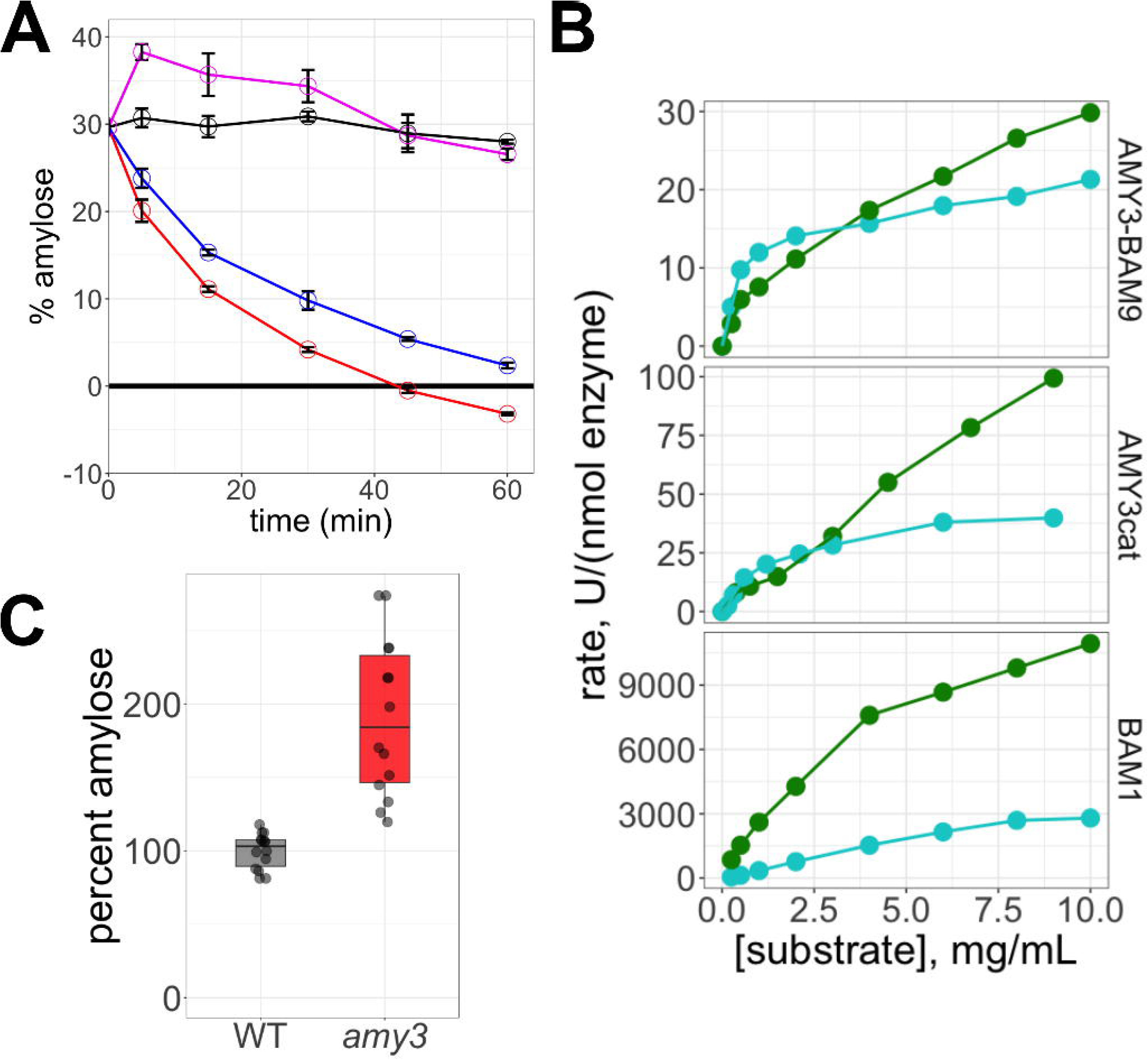
Characterization of Arabidopsis AMY3 substrate specificity. **A**. Effect of purified AMY3 (blue), AMY3 _cat_ (red), BAM1 (pink) or no enzyme (black) on the percent amylose content of solubilized corn starch over time measured using iodine spectroscopy. Points are means +/- standard deviation (n=3). **B**. Kinetics of AMY3 (+BAM9) (upper panel), AMY3_cat_, and BAM1 (lower panel) assayed with various concentrations of solubilized amylose (light blue) or amylopectin (green). Rates are μmoles of reducing equivalents produced per min per nmol enzyme and are representative of 2 to 4 independent experiments for each enzyme. **C**. Percent amylose in starch extracted from leaves of WT Arabidopsis and a mutant lacking AMY3 (*amy3*). Data were from four independent experiments; each was normalized to the average percent amylose in the WT leaves. All data points are shown, and the boxes indicate one standard deviation from the mean.

We then attempted to measure the affinity of AMY3 for amylose and amylopectin, along with BAM1, using enzyme assays. Amylose and amylopectin are challenging substrates with which to measure kinetic constants because of their structural complexity, lack of solubility, and because the relative number of cleavable chains may be affected by the actions of the amylase. However, the specificity differences between AMY3 and BAM1 were still apparent in the differences in estimated substrate concentration required to produce half the maximum rate (K_0.5_). We included BAM9 to enhance AMY3 activity in the assays, but its presence did not alter the shape of the substrate concentration curve, suggesting that BAM9 does not affect AMY3 specificity (Supplemental Figure 3). At low concentrations of substrate, AMY3 had more activity per mole of enzyme with amylose than with amylopectin, and the estimated K_0.5_ value was about 10-fold lower with amylose than with amylopectin (Figure 1B). At higher substrate concentrations, AMY3 was more active with amylopectin. In contrast, BAM1 was far more active with amylopectin than with amylose at all concentrations used, and the estimated K_0.5_ value was about 10-fold lower with amylopectin than with amylose (Figure 1B). Lastly, AMY3_cat_ also had a high affinity for amylose, like the full-length enzyme (Figure 1B). While both enzymes could act on both substrates, AMY3 had a higher apparent affinity for amylose, and BAM1 had a higher apparent affinity for amylopectin. If AMY3 prefers amylose as its substrate, then plants lacking AMY3 might contain starch with a higher percentage of amylose. Moreover, because of AMY3’s predicted daytime activity resulting from its redox regulation (19), we reasoned that any difference might appear at the end of the day. Indeed, leaves from the *amy3* mutant had a percent amylose content, on average, almost 2-fold higher than those of WT leaves at the end of the day (Figure 1C). The *amy3* leaves contained no observable starch at the end of the night (Supplemental Figure 4), as previously observed by Yu et al. (2005) (16). Therefore, the difference in percent amylose we measured between WT and *amy3* leaves developed during the 12-hour light period before harvesting. Together, these results suggest that AMY3 has a preference for amylose and that specificity is determined at least in part by the catalytic domain.

### Bioinformatic Analysis of the AMY3 CBMs

Given that the CBMs of AMY3 were not required for catalytic activity or even amylose preference, what role does the CBM play? Given the limited structural and mechanistic information available on AMY3, it was difficult to predict based on the literature. To provide some information about the structure, we compared the sequence conservation and organization of the CBMs of AMY3 with those in GWD1. AMY3 and GWD1 are the only two classes of proteins that contain CBMs classified in CAZy family 45 (48). Both enzymes are plastid-localized and can act on starch. Additionally, both contain tandem, N-terminal CBMs with a conserved “HWG/A” motif containing the tryptophan residues involved in carbohydrate binding (22). We compared the amino acid sequences of the two CBM regions from AMY3 and GWD1 sequences from 20 species of flowering plants (Supplemental Figures 1 and 2). The selected species represent 20 different orders of flowering plants to avoid inclusion of closely related species. The CBM sequences were aligned, and average pairwise percent identities were obtained for each of the four CBM sets (Figure 2A). When aligned with their orthologs (e.g. AMY3-CBM1 vs 19 other AMY3-CBM1 sequences), each of the four CBMs ranged from 55-65% identical to their respective orthologs. Each CBM1 had a higher percent identity than the CBM2s from the same gene, but we do not believe these differences are consequential. In contrast, when comparing the sequence of each CBM to the second CBM within the same gene, or to each CBM of the other gene, all the percent identities were approximately 26% and did not differ from one another. Despite sharing the common “HWG/A” motif, the overall sequences of these four CBMs are different from each other between the two genes, suggesting a lack of functional similarity.

**Figure 2.**
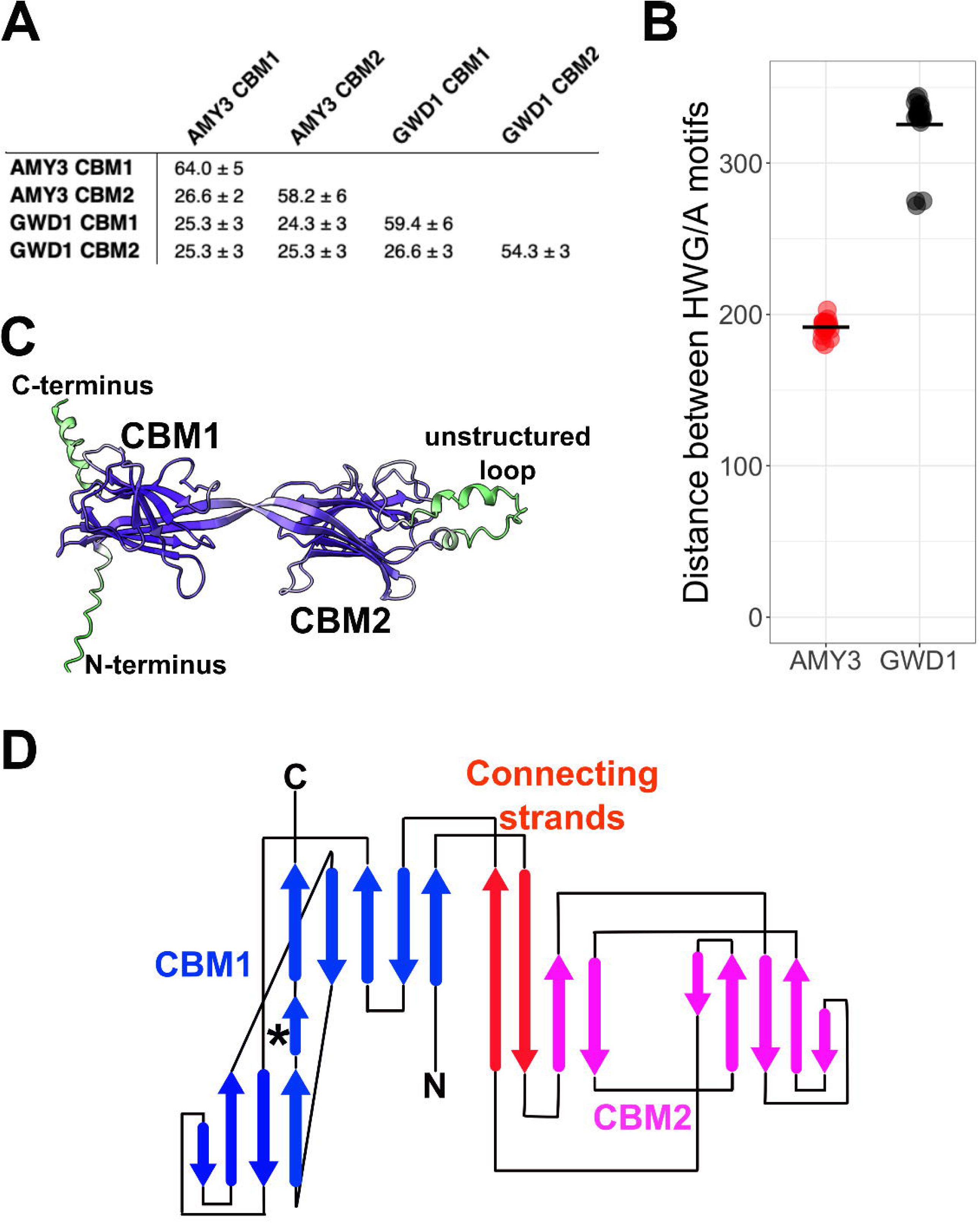
Bioinformatic analysis of the carbohydrate binding modules of AMY3 and GWD1, and the predicted structure of the AMY3 CBM duplex. **A**. Average pairwise percent identities of CBM1 and CBM2 regions of AMY3 and GWD1 from twenty species of angiosperms (see Supplemental Table 1 for species and accession numbers). Only one species per order of angiosperms was used. **B**. Number of amino acids between the conserved tryptophan residues in the “HWG/A” motif of each CBM region from AMY3 and GWD1 from the same 20 species used in A. **C**. AlphaFold3-generated structure of the CBM1-CBM2 duplex from AMY3. Amino acids are colored by pLDDT score. Blue colors indicate high confidence, while green colors indicate lower confidence. **D**. 2-D topology diagram showing the integrated duplex structure of CBM1 and CBM2 from AMY3. The strand marked with an * indicates the predicted dimer interface of the CBM duplex.

We then analyzed the relative spacing between the CBMs of each gene. In the primary sequences, the conserved tryptophan residues of the two CBM domains of GWD1 are separated by an average of 326 +/- 22 residues, whereas the two CBMs of AMY3 are separated by only 192 +/- 5 residues (Figure 2B). Glaring et al. (2011) previously noted that the two CBMs of AMY3 in Arabidopsis are separated by a shorter linker than the two CBMs of GWD1 (21); however, they acknowledged that no structures were available for the domains from either enzyme at that time. We then used AlphaFold3 to generate putative structures for each gene, and in doing so, the predicted structural differences between AMY3 and GWD1 became apparent (49). Overall, the structure of the CBMs was the expected Ig-like fold (50). The differences are in the organization and connection between the CBMs. The CBMs of GWD1 are likely to fold independently and are separated by a long, disordered loop that is predicted to contain one or two helical bundles between the two CBMs. The two CBMs are sufficiently far apart to be positioned on opposite sides of the catalytic domain in the AlphaFold3 model; however, this position should not be interpreted as representing the actual orientation in 3D space (Supplement Figure 5). In contrast, the predicted structure of the two CBMs of AMY3 suggests that an antiparallel set of β-strands connects them. In the model, the distance between the two AMY3 CBMs was a span of only 10 residues (Figure 2B). Each of the five models of the CBM duplex produced by AlphaFold3 had this same basic structure (Figures 2C and 2D). We termed this structure a duplex CBM, as opposed to a tandem CBM, because, despite being derived from an ancient duplication event, the two CBMs fold as a single unit (Figure 2C). In addition, each CBM duplex contains a poorly conserved and unstructured loop that is within CBM2 and is not a spacer between the CBMs, as might be assumed from the sequence alignments (Figures 2C and 2D). Our analysis reveals that despite being homologous, the CBMs of AMY3 and GWD1 have important differences in structure and function that may have been masked by their common CAZy family number.

### Biochemical Analysis of the AMY3 CBM

Given the differences in the structural models of the CBMs in GWD1 and AMY3, we next turned to support our proposed model of the AMY3 duplex CBM. Previous characterization of the AMY3 CBMs suggested that they could not purify it out of the context of the full-length AMY3 (21). We therefore tried a distinct construct of residues 56-391 containing just the duplex CBM of AMY3 with an N-terminal His tag, finding that we could purify the AMY3 CBM to apparent homogeneity on SDS-PAGE using a Ni-NTA affinity column (Figure 3A). This construct is longer than the construct used by Glaring, *et al.* (21), which may explain our success. Because we observed the protein eluted as a single peak after the void volume on size-exclusion chromatography, we believed the protein to be folded, which we then confirmed using CD (Figure 3B). The secondary structure content of the AlphaFold3 model shows overall agreement with the primarily β-sheet and coil structure, supporting the further use of this model (Figure 3B). Together, these data suggested we had obtained pure, folded AMY3 CBM for structural and biochemical study.

**Figure 3.**
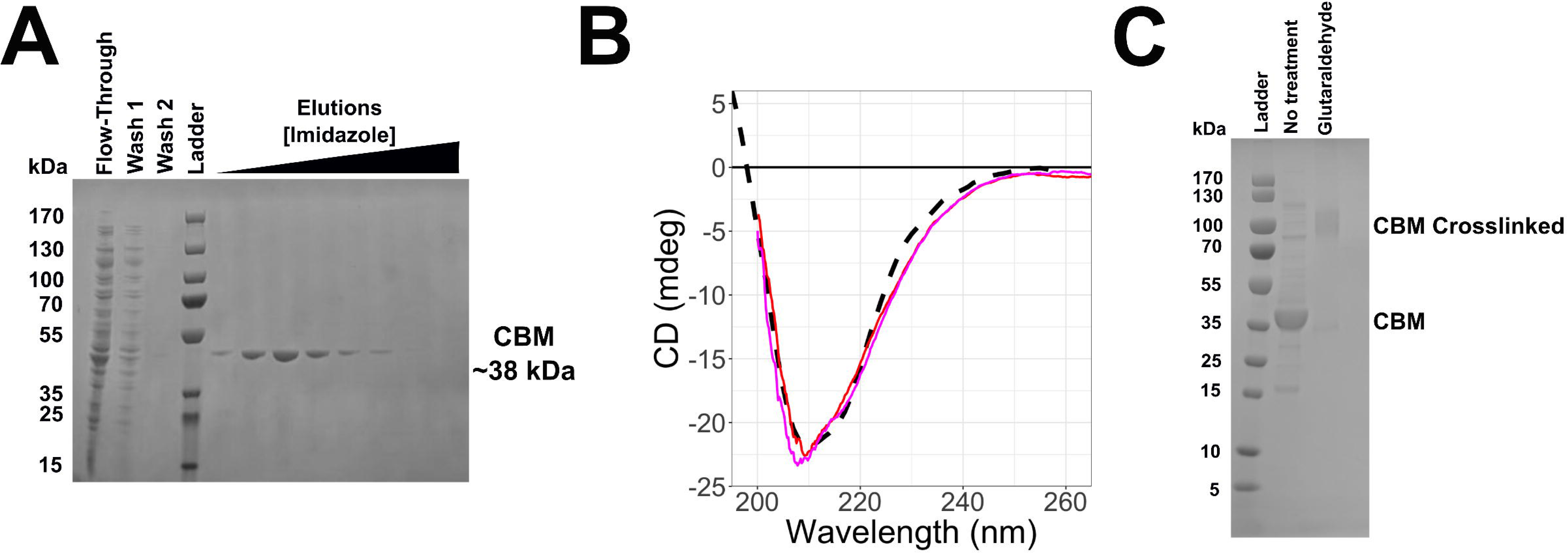
Purification of the duplex CBMs of AMY3. **A.** Nickel affinity purification of Arabidopsis CBM (residues 56-391 + C-terminal His tag). The predicted molecular weight of the expressed protein sequence is 38.8 kDa. **B.** Circular dichroism of the CBM. Data were collected at 10 °C in 20 mM HEPES, 50 mM potassium chloride, and at a pH of 7.0. The predicted CD from the SAXS model is shown as a dashed line, while the red and magenta lines are two representative experimental spectra of the CBM duplex. **C.** Glutaraldehyde crosslinking of the CBM.

Given that AMY3 forms dimers in solution (20), we investigated whether the duplex CBM itself also forms dimers. We observed on reducing SDS-PAGE that the duplex CBM domain alone migrated between the 35 and 55 kDa markers, as predicted by the sequence weight of 38.8 kDa. When crosslinked with glutaraldehyde, the primary band migrated similarly to the 100 kDa marker, suggestive of a dimer (Figure 3C). We also observed a second band that migrated more quickly than the CBM without any crosslinking, indicative of an intramolecular crosslink (Figure 3C). We further investigated whether the presence of sugars or soluble starch altered the crosslinking of CBM, which might suggest a regulatory role of the domain. However, we observed minimal effects of sugars on the crosslinking of the CBM, suggesting that sugar binding does not block any of the crosslinking sites of CBM (Supplemental Figure 6). Together, these data indicate that the CBM is one of the dimerization domains of AMY3, which may be important for the regulation of activity.

### Small Angle X-Ray Scattering (SAXS)

Because there is limited structural information on CBMs and no information on CBMs from the CBM45 family, we then used a hybrid modeling/SAXS approach to describe the structure of the AMY3 CBM duplex. Using SAXS and Kratky replotting, we found that the CBM formed a mostly globular structure, supporting our conclusion that we had purified a folded protein. Moreover, the molecular weight values we calculated from the two data sets were 74.3 and 83.1 kDa, which are approximately 2 times the expressed sequence molecular weight of 38.8 kDa (Figure 4A and Table 1). Collectively, these data suggest that the duplex CBM forms a dimer of duplexes. When we fitted the AlphaFold3 model to the SAXS data, a monomeric structure did not fit well, with an X^2^ value of approximately 103. When we used AlphaFold3 to predict a dimeric structure, the X^2^ value improved to 12. We further refined the CBM duplex dimer model to adjust the relative orientation of the CBM monomers and then used BilboMD to adjust the conformation of flexible regions (40). We eventually produced a model with an X^2^ value of 1.01 (Figure 4B). Additionally, we fitted and attempted to refine an AMY3 CBM dimer model generated by ESMfold, which is structurally similar to the models predicted for GWD1 (Supplemental Figure 7). The best-fitting dimeric model had an X^2^ of 42, while the best multi-state model with flexibility accounted for had an X^2^ of 13, indicating that a GWD1-like model of the AMY3 CBM does not fit well to the SAXS data (Supplemental Figure 7) (51).

**Figure 4.**
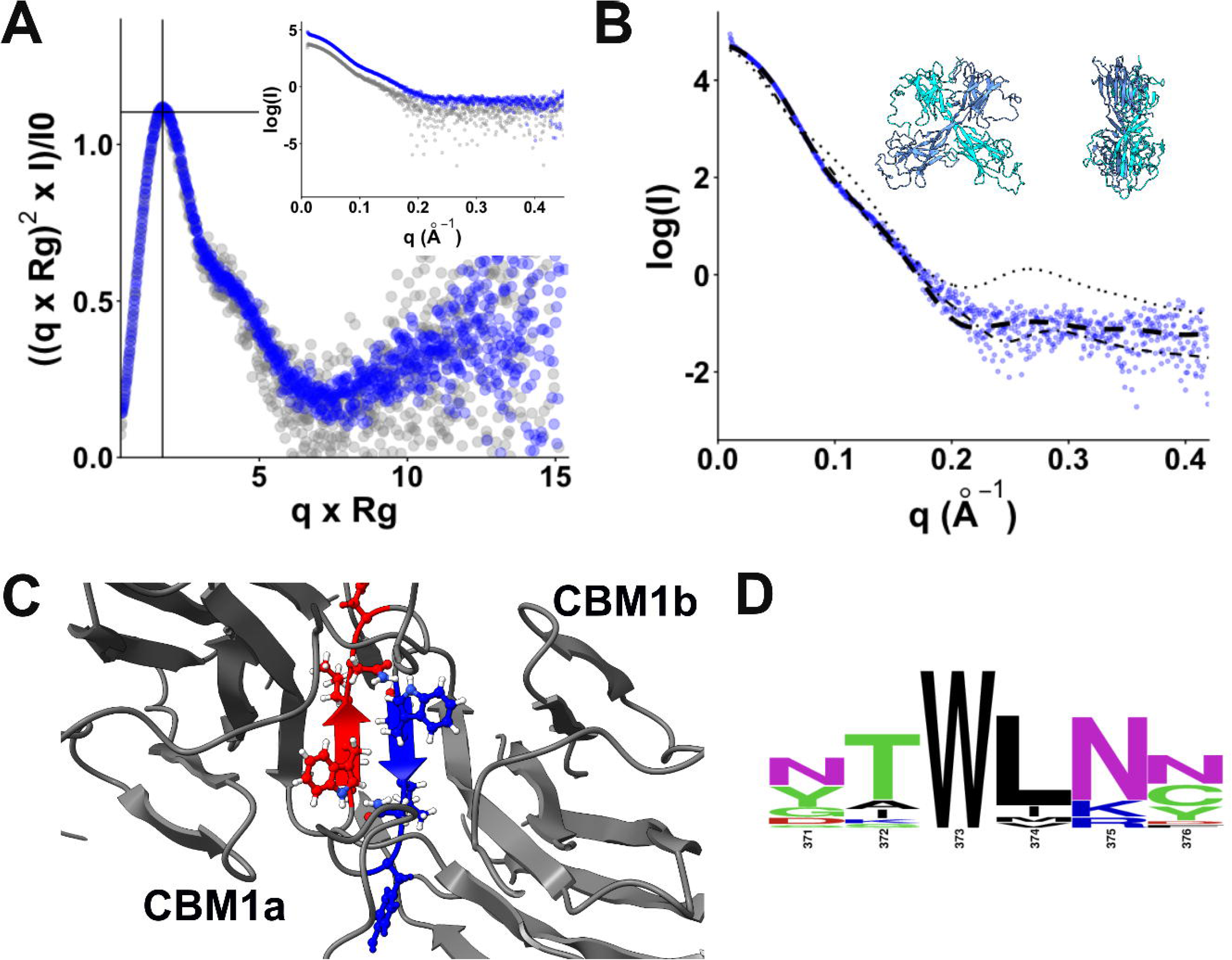
Structural description of the CBM dimer. **A.** Dimensionless Kratky replot of the size-exclusion chromatography coupled to SAXS data for the CBM. The two colors represent separate runs of two concentrations. The inset shows the same data in a log(intensity) vs. q plot. **B.** Fitting of the SAXS data to models of the CBM. A fit to the monomeric models is shown as a dotted line, the fit to the initial AlphaFold3 model is shown as a dot-dashed line, and the fit to the final dimeric model is shown as a heavy dashed line (FOXS fit X^2^ = 1.01). **C.** Dimer interaction strand from the best fitting model of the CBM. The predicted interacting strand is marked with an * in Figure 2D. **D.** Weblogo showing the conservation of the interaction strand shown in C using the same 20 sequences described Figure 2.

**TABLE 1.**
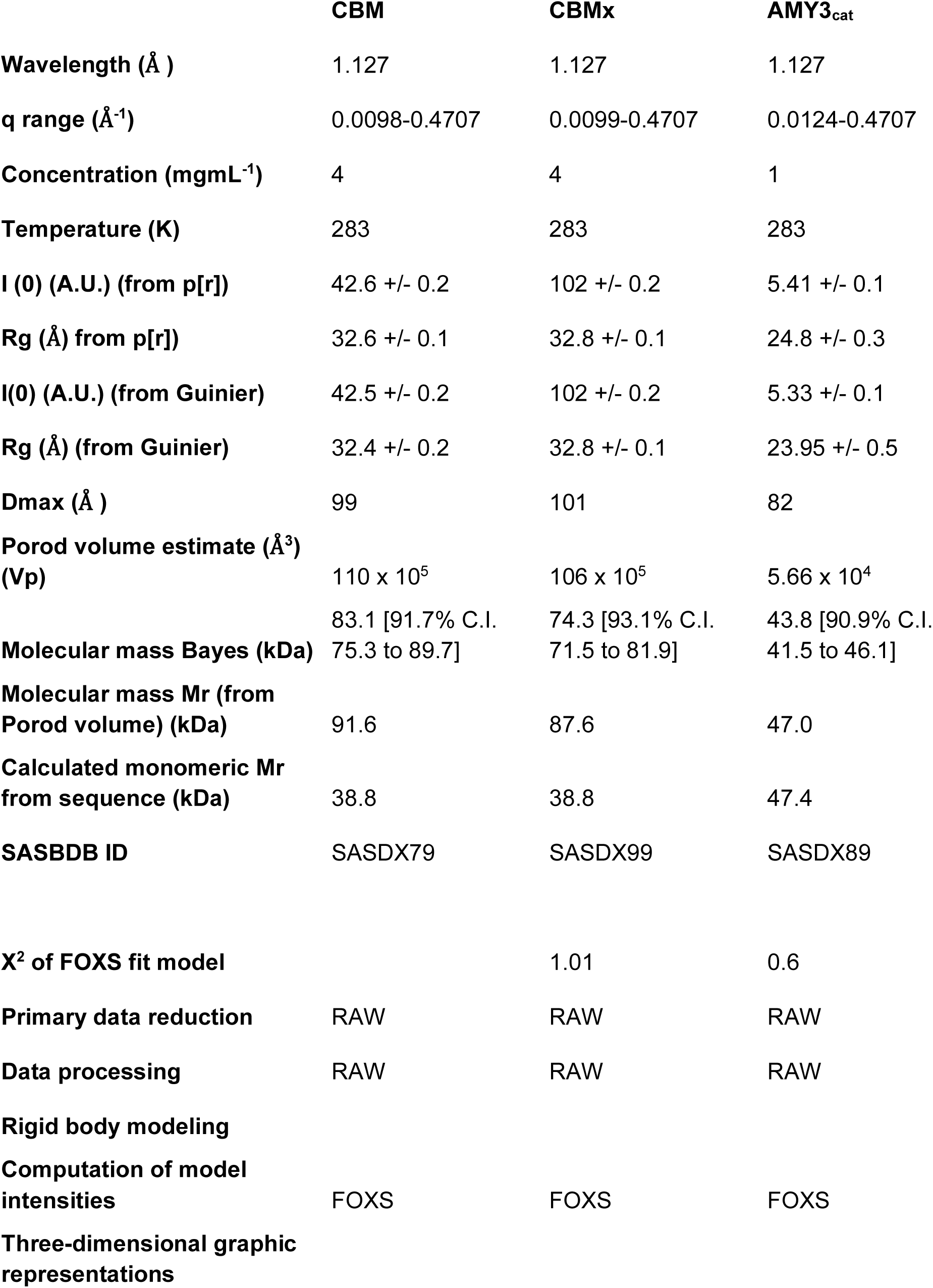
SAXS statistics.

|  | **CBM** | **CBMx** | **AMY3<sub>cat</sub>** |
| --- | --- | --- | --- |
| **Wavelength (Å )** | 1.127 | 1.127 | 1.127 |
| **q range (Å<sup>-1</sup>)** | 0.0098-0.4707 | 0.0099-0.4707 | 0.0124-0.4707 |
| **Concentration (mgmL<sup>-1</sup>)** | 4 | 4 | 1 |
| **Temperature (K)** | 283 | 283 | 283 |
| **I (0) (A.U.) (from p[r])** | 42.6 +/- 0.2 | 102 +/- 0.2 | 5.41 +/- 0.1 |
| **Rg (Å) from p[r])** | 32.6 +/- 0.1 | 32.8 +/- 0.1 | 24.8 +/- 0.3 |
| **I(0) (A.U.) (from Guinier)** | 42.5 +/- 0.2 | 102 +/- 0.2 | 5.33 +/- 0.1 |
| **Rg (Å) (from Guinier)** | 32.4 +/- 0.2 | 32.8 +/- 0.1 | 23.95 +/- 0.5 |
| **Dmax (Å )** | 99 | 101 | 82 |
| **Porod volume estimate (Å<sup>3</sup>) (Vp)** | 110 x 10<sup>5</sup> | 106 x 10<sup>5</sup> | 5.66 x 10<sup>4</sup> |
| **Molecular mass Bayes (kDa)** | 83.1 [91.7% C.I.<br>75.3 to 89.7] | 74.3 [93.1% C.I.<br>71.5 to 81.9] | 43.8 [90.9% C.I.<br>41.5 to 46.1] |
| **Molecular mass Mr (from Porod volume) (kDa)** | 91.6 | 87.6 | 47.0 |
| **Calculated monomeric Mr from sequence (kDa)** | 38.8 | 38.8 | 47.4 |
| **SASBDB ID** | SASDX79 | SASDX99 | SASDX89 |
| **X<sup>2</sup> of FOXS fit model** |  | 1.01 | 0.6 |
| **Primary data reduction** | RAW | RAW | RAW |
| **Data processing** | RAW | RAW | RAW |
| **Rigid body modeling** |  |  |  |
| **Computation of model intensities** | FOXS | FOXS | FOXS |
| **Three-dimensional graphic representations** |  |  |  |

We next focused on the interaction site in the CBM model from Figure 4B. We predicted a parallel strand interaction between the second strand in each monomer and an anti-parallel strand interaction between the final strands in each monomer to form the dimer interface in the best-fitting model (Figures 2D and 4C). The interface sequences are located within conserved regions of the alignment (Figure 4D). The final model suggests that AMY3 forms a dimer between the CBMs of each AMY3 monomer.

#### AMY_cat_ structure

We and others have proposed that AMY3 is a multidomain protein, and therefore, it could strengthen dimerization through domains other than just the CBMs (19–21). We collected SAXS data on a construct corresponding solely to the AMY3 catalytic domain, AMY3_cat_, and examined whether this catalytic domain alone could form a dimer. Overall, the SAXS data indicated a globular protein, which is consistent with the structures of other alpha-amylase domains (52–57). The molecular weight calculated from the SAXS data was 44 kDa [90.9% C.I. 41.5 to 46.1 kDa] (Table 1). The expected molecular weight of the expressed sequence was 47 kDa; thus, these data are more consistent with a monomeric AMY3_cat_. We generated a model of the AMY3_cat_ domain using AlphaFold3 and then fitted the monomeric model of AMY3_cat_ to the SAXS data using FoXS, finding an X^2^ value of 0.6 (Figure 5B). Given our finding that AMY3_cat_ does not form a dimer, but the CBM does, we then refined our previous model of dimeric, full-length AMY3 with the restraint that dimerization occurs through the CBM. The three models generated by CORAL were compared to the AMY3 dimer SAXS data published by Berndsen *et. al.* (20, 58). The χ² values ranged from 1.43 to 1.54, with the best-fitting model shown in Figures 5C and 5D. These data indicated that these new models with the CBM as the dimerization interface are plausible.

**Figure 5.**
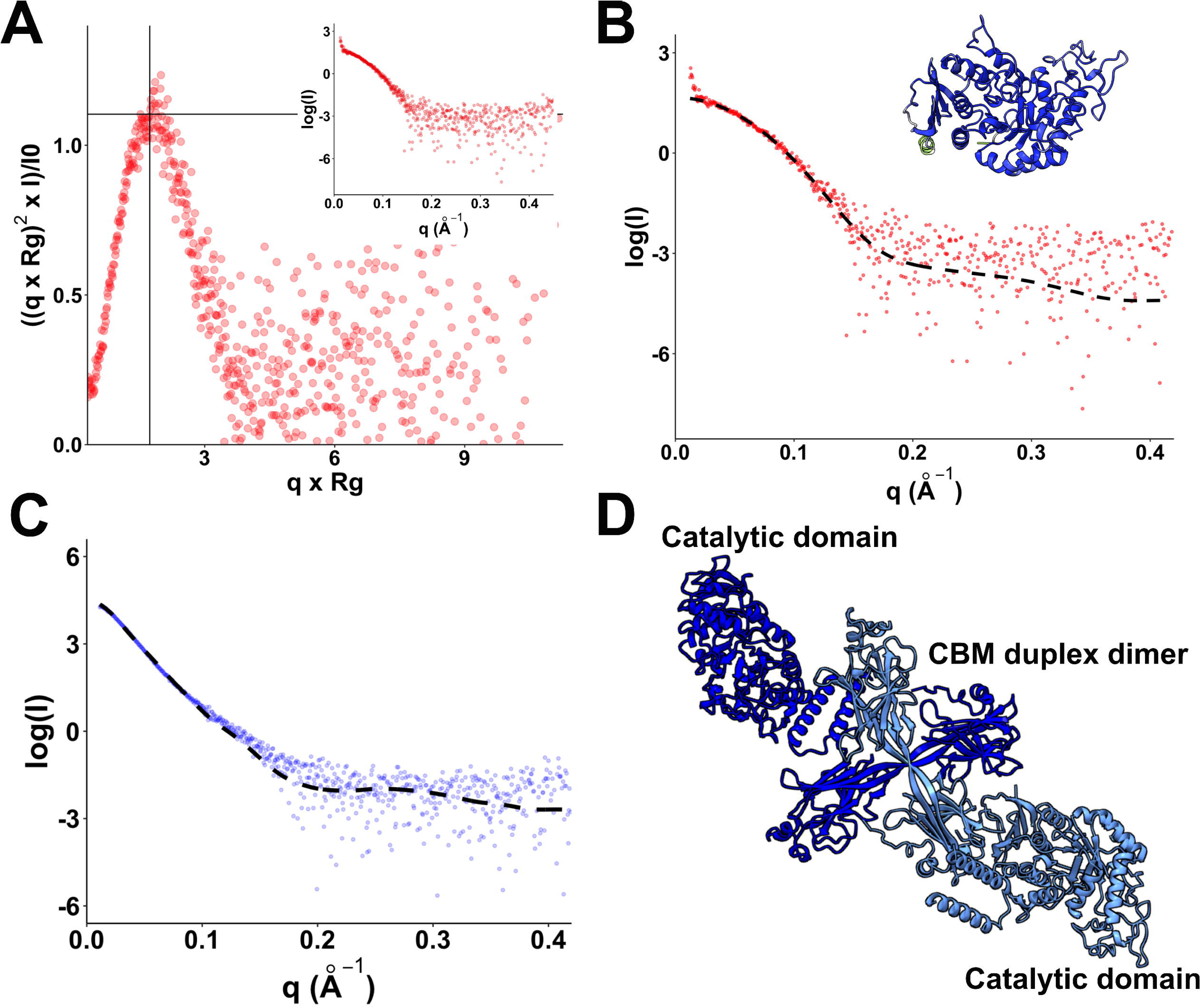
Structural description of the AMY3 catalytic domain. **A.** Dimensionless Kratky replot of the size-exclusion chromatography coupled to SAXS data for AMY_cat_. The inset shows the same data in a log(intensity) vs. q plot. **B.** Fitting of the SAXS data to the AlphaFold3 model of AMY3_cat_. (FOXS fit X^2^ = 0.6). **C.** Fitting of the full-length AMY3 dimer with the CBM as the interaction surface shown in **D.** to the SAXS data previously published in Berndsen et al. (20). The X^2^ range from FOXS fitting was 1.43 to 1.54.

#### Starch binding assay

Having developed confidence that we had purified a folded CBM, we next determined whether the CBM could bind to amylose or was only involved in regulating AMY3 activity through dimerization. We performed a coprecipitation binding assay to test the interaction between CBM and starches with different amounts of amylose. The CBM protein appeared in both the pellet and the supernatant when binding to amylopectin (Figure 6A). In contrast, we observed that CBM was primarily found in the pellet of soluble starch, wheat starch, corn starch, and pea starch, all of which contain amylose in addition to amylopectin. We further investigated the influence of pH on CBM binding to corn starch by performing binding assays across a range of pH conditions and at near constant ionic strength (59). We found that CBM binds most effectively at pH 7, as indicated by a darker band in the pellet and a very faint band in the supernatant. This suggests that CBM has stronger starch-binding activity at pH 7, which falls within the pH range of the chloroplast (7 to 8).

**Figure 6.**
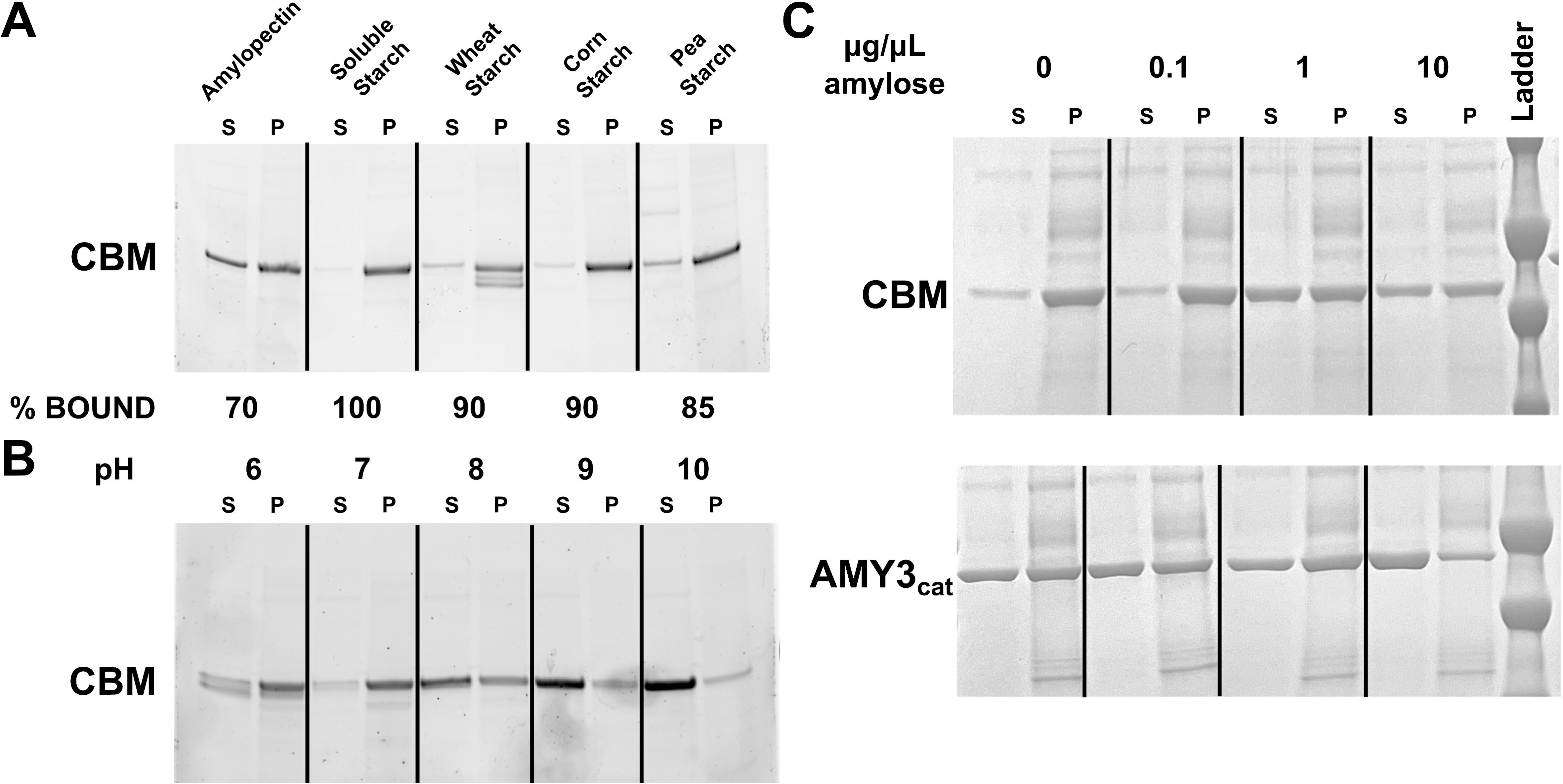
Starch-binding properties of the CBM of AMY3. **A.** Carbohydrate pulldown of the CBM with amylopectin or starch from the indicated sources. Bands in “S” lanes show protein that did not bind, while those in the “P” lanes were in the carbohydrate pellet. **B.** Pulldowns of CBM using soluble starch at several pH values. **C.** Amylopectin pulldown assays using amylose at the indicated concentrations as a soluble competitor.

To test whether the CBM prefers amylose binding, we conducted amylopectin binding assays with varying amounts of soluble amylose to act as a competitor. In the absence of amylose in the supernatant, CBM was mainly in the pellet, and AMY3_cat_ was equally split between the pellet and supernatant (Figure 6). There was no effect of 0.1 mg/mL amylose on the binding of either protein, but with 1 mg/mL amylose as a competitor, more CBM appeared in the supernatant. Only at 10 mg/mL amylose did the amount of AMY3_cat_ in the supernatant increase. This suggests that both the CBM and the catalytic domain will bind to amylose, but the CBM appears to have a higher affinity for amylose than the catalytic domain. Collectively, these data show that the CBM forms dimers in solution and can bind preferentially to amylose, supporting our finding that AMY3 prefers amylose as its substrate.

## DISCUSSION

This investigation was driven by two different goals: to understand the function of AMY3 in leaf starch metabolism and to understand the role of the N-terminal carbohydrate-binding module in the function of AMY3. Regarding the former, despite knowing of AMY3’s presence in leaf chloroplasts and decades of investigations on leaf starch metabolism, primarily using Arabidopsis, little has been revealed about the catalytic function of AMY3. Mutants lacking AMY3 appear to contain normal levels of starch at the end of a 12-hour day period and are capable of degrading leaf starch completely over a 12-hour night period (16) (Supplemental Figure 4). Knowing that α-amylases are endo-amylases, that amylose has few non-reducing ends, and that AMY3 is active under reducing conditions typical of chloroplast conditions during the day, we wondered if the starch in *amy3* plants might contain a higher percentage of amylose than WT leaves. Indeed, we observed variable but consistently higher apparent percent amylose in starch extracted from *amy3* leaves compared to WT leaves at the end of a 12 hr day (Figure 1C). Whether this elevated apparent amylose is bona fide amylose or merely extra-long chains of amylopectin is unclear from the current data.

Using *in vitro* assays, we found that Arabidopsis AMY3 preferentially degrades the amylose component of corn starch (Figure 1A) and has a higher affinity for amylose than for amylopectin (Figure 1B). In contrast, the exoamylase BAM1 preferentially degrades the amylopectin component of corn starch and has a higher affinity for amylopectin compared to amylose. Previous activity assays on full-length AMY3 using tobacco leaf starch as a substrate produced a K_0.5_ value of 36 mg/mL (21), which is within an order of magnitude of our K_0.5_ value for amylopectin (21). We did not fully saturate the enzymes with amylopectin; thus, our K_0.5_ value represents a lower limit on the affinity. The difference in values is likely because we were using substrates that are mostly amylopectin or amylose, while Glaring and coworkers used purified leaf starch which is a mixture of amylose and amylopectin (21). Overall, our results are consistent with and help to explain the observations made on barley starch degradation by different forms of amylase in the 1800s (24).

Arabidopsis AMY3 and its orthologs in other plant species are unique among plant ɑ-amylases due to their N-terminal extension containing a duplex CBM45 domain, association with and activation by BAM9, and apparent dimerization (19–21). To investigate the molecular mechanism of the preference for amylose and dimerization, we focused on the tandem CBMs in the N-terminus of AMY3 using purified CBM alone and the catalytic domain alone. Experiments using a truncated form of AMY3 lacking all but the catalytic domain indicated that AMY3_cat_ behaved like the full-length protein in preferring the amylose fraction of corn starch and in its kinetic preference for amylose (Figures 1A and 1B). This finding was unexpected, given that the duplex CBM of AMY3 can bind to starch (Figure 5). The lack of an apparent effect of the CBM on *in vitro* activity could be due to the lack of an additional component and that the *in vitro* conditions do not fully reflect the environment surrounding a starch granule. Alternatively, the CBM may not determine substrate specificity but could serve as a regulatory domain, aligning with our previous findings and proposal that full-length AMY3 forms dimers, which regulate its activity (20). We do want to note that these scenarios are not exclusive and support further biochemical and organismal work into this system.

Studying the CBM domain in isolation to understand its function has been challenging because this domain had not been successfully purified before (21). We were able to purify this domain and characterize its structure using SAXS (Figure 3). We found that it dimerized in solution and that this domain appears to be the main dimer interface within AMY3, as the catalytic domain alone did not form dimers in our hands (Figures 3 and 4). Binding assays indicated that the CBM appears to prefer carbohydrate polymers that contain amylose, but that the CBM could also bind to amylopectin. These data are in line with the enzyme assays of AMY3 and AMY3_cat_ showing their similar specificity toward amylose, but the amylose-specific behavior of AMY_cat_ suggests that the CBM is not required for the amylose-cleaving properties of AMY3. However, we cannot rule out additional involvement of the AMY3 CBM on a starch granule or under physiological conditions. The former point is supported by the preference of the duplex CBM for amylose-containing starch over amylopectin *in vitro* (Figure 5B). Our findings suggest that AMY3 is involved in amylose degradation in Arabidopsis and identify the CBM domain as a novel amylase dimerization domain. It is unclear from the existing work how BAM9 interacts with the hairpin to disrupt the CBM homodimer (20).

While amylases and other starch-acting enzymes are known to contain one or more CBMs, these modules/domains are not frequently found in duplex and close sequence proximity, as in AMY3 (50). Moreover, few, if any, CBMs are known to occur as dimers in solution (50, 60). Thus, the dimerization of the duplex CBMs of AMY3 is unique. CBM45 domains also occur in GWD1; however, the spacing of these domains is not conserved and is far larger than the spacing in AMY3 homologs. This larger spacing is likely not conducive to dimerization, and our predicted model of the AMY3 CBM45 dimer forms in part via interactions between the β-strands that compose the linker connecting the CBMs in each monomer. Functionally, GWD1 is known to phosphorylate the outer chains of amylopectin (6), while we show that AMY3 has a substrate preference for amylose. The different structures of the corresponding CBMs in these enzymes may play a role in their different substrate specificities. These differences between the CBMs in GWD1 and AMY3 suggest that the CBM45 family might need revision to differentiate between these distinct groups. Thus, we propose that CBMs connected by a flexible linker, as in GWD1, should be referred to as tandem CBMs, while those connected by a rigid linker should be called duplex CBMs.

The role of AMY3 in regulating the amylose content of starch granules is an intriguing finding with numerous potential applications. Starch with high amylose levels has been shown to offer benefits for human health and is thought to have properties similar to those of a high-fiber diet, with anti-obesity effects (61, 62). Arabidopsis plants lacking AMY3 accumulate starch with a higher percentage of amylose, suggesting that AMY3 prevents the generation of High-Amylose Starch. Given the conservation of AMY3 in various species of industrially relevant plants (20), it is tempting to speculate that genetic modification of AMY3 could lead to the generation of starch with high amylose in a broader set of plants, diversifying the potential human uses. Modification could occur in the catalytic domain, which we demonstrate appears to be the primary domain in dictating the enzyme’s specificity toward amylose, or in the duplex CBM, which we have shown is important for dimerization, regulating AMY3 function (20).

## Acknowledgements

National Science Foundation grants MCB RUI-2322867 and CHE REU-2150091 supported this work. This work was conducted in part at the Advanced Light Source (ALS), a national user facility operated by Lawrence Berkeley National Laboratory on behalf of the Department of Energy, Office of Basic Energy Sciences, through the Integrated Diffraction Analysis Technologies (IDAT) program, supported by DOE Office of Biological and Environmental Research. Additional support came from the National Institute of Health project ALS-ENABLE (P30 GM124169) and a High-End Instrumentation Grant S10OD018483.

## Notes

### Competing Interest Statement

The authors have declared no competing interest.

