## Supplemental Figure 1 for "Functional Characterization of Arabidopsis α-Amylase3 (AMY3): Amylose Specificity and Structural Insights into Its Duplex Carbohydrate-Binding Module"

```
                             1 ........10........20........30........40........50........60
Erigeron_canadensis_AMY3     1 ---MATVPLGTHLHHSCLFETTPNN--NKFS----S---KQSCFYYNNKKHSNFMLNNSN
Nymphaea_colorata_AMY3       1 ---MATLRVR----PPLYQ---FRRTYPRFVHKLG--GSRPSSLSFSRSV---S------
Erythranthe_guttata_AMY3     1 ---MSAVATELLFHHRVHRISP--KLIPKFH--LISRKSPTFNLICDTTT---T--D---
Beta_vulgaris_AMY3           1 ---MSTIAAE-----PLLHLALRQPTPK--IQPIKSRKNLPFSLNFSSRA---SKI----
Daucus_carota_AMY3           1 MSTM--VALG-----PLVHHQIPS--LPT----LFTKTPLTSSLNYPHRA---P------
Arabidopsis_thaliana_AMY3    1 ---MSTVPIESLLHHSYLRHNSKVNRG--------NRSFIPISLNLRSHF---T--SNKL
Ipomoea_triloba_AMY3         1 ---MSTVTID-----PLIHHSFRRTPKFCLN----HRQPSPSSVNCTR--------RALS
Ricinus_communis_AMY3        1 ---MSTL-----TVEPLLRFSGREKSLPIGS----RKILKPSSL--NF--------SKKL
Gossypium_hirsutum_AMY3      1 ---MAAVTIDSVLQNPTLLFRPKAKVM----------LKPPRSLDCSR--------NRKL
Phaseolus_vulgaris_AMY3      1 MSIYTTLSFD-----PLFSFNHCVNREREPS--IHSSRPKLFSLTSSSTL---TLFNSNN
Cucumis_sativus_AMY3         1 ---MSSIALD-----PLLY--HCAKGKHRFHHRPRFNMLRPCSFTYCPNK---L------
Fragaria_vesca_AMY3          1 ---MSTVSIE-----PLLH--HCRRGNSRHR--SASSS-KLIKLSYLSAF---P--KKV-
Nelumbo_nucifera_AMY3        1 ---MSTVTLE-----PLLH--QCCRQRVIFR--LESKKLRPSRVNYSPKL----------
Vitis_vinifera_AMY3          1 ---MSTVCIE-----PLFQ--RCRRENPRFR--LKSLATKPSSLNYSPK-----------
Camellia_sinensis_AMY3       1 ---MSTFALE-----PLGY--HCRREHPNFR--PNYKKSKAFSLNYTPRP---R--RRPR
Zea_mays_AMY3                1 ---MS-----------VGS--CCIRAIPGAAPPAR-VGLLGSA------FLQVSA-----
Amborella_trichopoda_AMY3    1 ---MATLRLK-----PSLH--HHTKWNPRSNQKLRNYSNWNPRLNHKLR-----------
Zingiber_officinale_AMY3     1 ---MALVHWK-----PVLH--RPPQETPGVIPRWKRRSFLGRQICRPRPV----------
Magnolia_sinica_AMY3         1 ---MSTIRLR-----PLLH--QCRRENPSFRHES-RRTKLSSGLNCGPKYYCYN------
Phoenix_dactylifera_AMY3     1 ---MSIVRWR-----PVLH--QPPRENPRFPPHELRRSKVLCPIHYRKPFFPAS------

                            61 ........70........80........90.......100.......110.......120
Erigeron_canadensis_AMY3    49 KCGGKASKTRPGFDSYWVMST-----NARLSANS-STSDTA---LLDTEVSDDVIFQEKF
Nymphaea_colorata_AMY3      40 ---D-----FRCLCNCKLRGR-----FRV-----SAGLAESFVEAAEAAASEEVVLRETY
Erythranthe_guttata_AMY3    46 RRFV-----GTGLCSRY-YKP----------LKAVSSAGAA---VLETSESSTITFSETF
Beta_vulgaris_AMY3          44 ---------RGSFSSNS---I-----QKSVSVAHCT-ATTT---LAQISEPTEVVYSEEF
Daucus_carota_AMY3          39 -LSV-----PTTSVSFKSSSY-----RPVVTARATETDATT---RTQTDQLADVVFTDTF
Arabidopsis_thaliana_AMY3   45 LHSI-----GKSVGVS--SMN-----KSPVAIRATSSDTAV----VETAQSDDVIFKEIF
Ipomoea_triloba_AMY3        41 VGAA-----RLSFCNCKHRPR-----RKALAVRAS----------STVIETSDVVFEETF
Ricinus_communis_AMY3       39 LLSN-----GSSFCNFKRSPP-----LSHT-VRASSTTDTALIET---FKSADVLFKETF
Gossypium_hirsutum_AMY3     40 PLSR-----GSCFCSFNPRRT-----IH-V-VRA-SSSETALIGNFDTSSSDDIFYKDIF
Phaseolus_vulgaris_AMY3     51 NCTY-----NYASCKPHRFHT-----P-K--FESFAT-NTD---TLESLQSSDVLFDRSF
Cucumis_sativus_AMY3        42 LCHG-----RKSFVHYNSYRP-----P-T--IKATT----T---NAPTFQSTDVLFNETF
Fragaria_vesca_AMY3         42 EELR-----GRSFCNFRRPTP-----L-T--LRASSADA-A---VAATFESTKPFFKQTF
Nelumbo_nucifera_AMY3       39 -CYH-----RRCFCNSKPY-R-----FRT--VRSSSTDAAL---V-EASEAADVSFRETF
Vitis_vinifera_AMY3         38 PLRN-----GGSFCNFKSLHG-----VRP--LGAAS----I---DTALFETTDVFFKETF
Camellia_sinensis_AMY3      44 PLSH-----GSIFCNFRPPLS-----L-P--LRATSTNTAL---VEETFESEEVLFKETF
Zea_mays_AMY3               33 --ARTRTRTRAGRCRAAQHG--LVRLGGRVVARAGAAETP----VAGTDEDAGAAFSETF
Amborella_trichopoda_AMY3   40 -----NSTFSGLNCIYKRFDIRSFSKIKPGVVVRASSTNT----SVEEAVASDVLFTETF
Zingiber_officinale_AMY3    41 --------FRS-ICCSKSSKD--DRKRTRPIVSSGATQTP----SLADVDEAKLLFTETF
Magnolia_sinica_AMY3        44 --NDSRKVFSD-FCKF--------KPWSLHRIRAGSTEMA----FVEADTTDRVLFKETF
Phoenix_dactylifera_AMY3    45 -----RGLSN----SY--------ARKALRIVRAGLAPTP----SLADAAETDVLFSETF

                           121 .......130.......140.......150.......160.......170.......180
Erigeron_canadensis_AMY3   100 PLQLVEKVEGKILIRLDSG-KD--EENWQLTIGCNLQGKWVLHWGVNYVD-DVGSEWDQP
Nymphaea_colorata_AMY3      82 PLKVTQKVEGKVYVRVDNG-KD--DLNWKLTIGCNLPGKWVLHWGVTYND-EIGSEWDQP
Erythranthe_guttata_AMY3    87 QLERVEKVEGKITIKLDKG-ET--EECWQLIVGCSIPGKWILHWGVNYVS-DVGSEWDQP
Beta_vulgaris_AMY3          83 ELPRLNKVEGKICVRLDQK--D--GDDWQLTVGCTLPGKWVLHWGVHYVG-DVGSEWDQP
Daucus_carota_AMY3          85 PVRRAQKVEGKLIIKLDKG-NS--EGKWELMVGCTLPGKWILHWGVNYVD-DTGSEWDQP
Arabidopsis_thaliana_AMY3   89 PVQRIEKAEGKIYVRLKEV--K--EKNWELSVGCSIPGKWILHWGVSYVG-DTGSEWDQP
Ipomoea_triloba_AMY3        81 DLKRPHRVEGRIAIRFDSG-RD--EENWKLTIGCNLPGKWVLHWGVHYAG-DSGSEWDQP
Ricinus_communis_AMY3       85 SLSRTETIEGKIFVRLDKEEKD--QQRWQLSVGCSLPGKWILHWGVSYVG-DVGSEWDQP
Gossypium_hirsutum_AMY3     87 PVKRIEKVEGKFFIRLDRS-KD--QHDWQLTVGCSLPGKWILHWGVSYLG-DSGSEWDQP
Phaseolus_vulgaris_AMY3     94 PINRTELVEGKIFVRLDHG-KD--LGNWELTVACNLTGKWILHWGVSRVD-DVGSEWDQP
Cucumis_sativus_AMY3        82 PLKRNEKLEGRISVRLAQG-KD--HNNWELTVGCNLAGKWILHWGVSLID-DSGSEWDQP
Fragaria_vesca_AMY3         85 PLERTELVEGKIYVRLDHG-KN--DRNWTLTVGCTLPGKWVLHWGVSHVDDDVVSEWEQP
Nelumbo_nucifera_AMY3       81 QLKRTERVEGKISVRLDPG-KD--EENWQLTVGCDLPGKWLLHWGVNYID-DVGSEWDQP
Vitis_vinifera_AMY3         79 ILKRTEVVEGKISIRLDPG-KN--GENWQLTVGCNIPGSWVLHWGVSYID-DVGSEWDQP
Camellia_sinensis_AMY3      88 PLKRTQKGEGKISIRLDNG-KD--QENWQLTVGCNLPGKWVLHWGVNYVN-DFGSEWDQP
Zea_mays_AMY3               85 PLRRCQAVEGKAWVRVDAEPDS--EGKCKVVVGCNVAGKWVLHWGVSYDD-EHGREWDQP
Amborella_trichopoda_AMY3   91 QLKRSEKVEGKISVRVDHQKDD---DKSQVAIGCNLPGKWVLHWGVTYYD-DVSSEWDQP
Zingiber_officinale_AMY3    86 PLKRPQTEEGKIIVRLDPVVEDGNASKWRLNVGCNLEGKWILHWGVSYCD-DLGSEWDQP
Magnolia_sinica_AMY3        89 LLKGTQMVEGKVSIRLDPQ--EG-GKNWQLTVGCNLPGKWVLHWGITYCG-DVGSEWDQP
Phoenix_dactylifera_AMY3    84 PLKRTQTVEGKISVRLDPA--EGEGSRWRLAVGCNLEGKWVLHWGVTYCD-DLGSEWDQP

                           181 .......190.......200.......210.......220.......230.......240
Erigeron_canadensis_AMY3   156 PVEMRPLDSITIKDYAVETPLTKSS---EADSLYEVKIDFNTKSSLAAINFVLKDAENGS
Nymphaea_colorata_AMY3     138 PAEMRPPGSVPIKDYAVETSLRKSTSSDGEQELHEIQIDVSNSYSIAAVHFVLKEEETGA
Erythranthe_guttata_AMY3   143 PLDMRPPGSIPVKDYAIETPLESKSELAEGEAFSEVAIVFDTKSSIASINFVLKDEETGR
Beta_vulgaris_AMY3         138 PNEMRPPGSVAIKDYAIETPLKQSSSSEGEDCLHEVNINFNCNSNIAAINFVLKDEEAGS
Daucus_carota_AMY3         141 PIEMRPPDSIPVKDYAIETPLKISSKEKEVENLQTVKIGFDSKSLIAAINFVLKDEETGA
Arabidopsis_thaliana_AMY3  144 PEDMRPPGSIAIKDYAIETPLKKLS---EGDSFFEVAINLNLESSVAALNFVLKDEETGA
Ipomoea_triloba_AMY3       137 PLEMRPQGSITIKDYAIETPLKRLSTALEGESFYEVEIDLNINSQIAAINFVLKDEETGA
Ricinus_communis_AMY3      142 PKNMRPRGSISIKDYAIETPLEKSS---EADMFYEVKIDLDPNSSIAAINFVLKDEETGA
Gossypium_hirsutum_AMY3    143 PKGMRPPGSIPIKDYAIETPLKKLS---EGDIFHEVKIDFNPSSEIAAIHFVLKDEETGA
Phaseolus_vulgaris_AMY3    150 PRDMIPPGSIPIKDYAIETPMQKSLSSAEGDALHEVKIDLKPNNDISAINFVLKDEETGA
Cucumis_sativus_AMY3       138 PKEMIPPGSITIKDYAIETPLKKSSSSSSGD-VHEVKIDLAPDKTIAAINFVLKDEETGI
Fragaria_vesca_AMY3        142 PEEMRPPGSIPIKDYAIDTPLTKLSSAVGGDNSQEVKIDFNLDGAIAAINFILKDEETGA
Nelumbo_nucifera_AMY3      137 PPEMIPPGSIPIKDYAIETPLKKSSSTSEGETFHEAKIKFNCNSSIVAINFVLKDEESGA
Vitis_vinifera_AMY3        135 PLEMRPPGSVAIKDYAIETPLKKLSSASERDTLHEVTIDFSPNSEIAAIRFVLKDEDYGA
Camellia_sinensis_AMY3     144 PVEMRPPDSILIKDYAIETPLKKSSAVVEGDLYYEVKINFSTNRDIAAINFVLKDEETGA
Zea_mays_AMY3              142 PSEMRPPGSVAIKDYAIETPLEILPN-SEGQPLYEMQIKFDKDIPIAAVNFVLKEEETGA
Amborella_trichopoda_AMY3  147 PPDMRPPDSIAIKDYAIETPLKKSPLAVEGNSLYEVQIDIKVNHSVGALHFVLKDEETGA
Zingiber_officinale_AMY3   145 PPEMRPPDSVVIKDYAIETPLKKGTSLSEGHIIYEAQIDFDSNIPISAIHFVLKEEETGA
Magnolia_sinica_AMY3       145 PPEMRPLDSVPIKDYAIETPLKKSPASLEGHLFYEVQIDFSSESSIAAIHFVLKEEGTGA
Phoenix_dactylifera_AMY3   141 PPEMRPPGSIPIKDYAIETPLKNSSSALKGQILHEVHIDFDSNSPIAAIHFVLKEEETGA

                           241 .......250.......260.......270.......280.......290.......300
Erigeron_canadensis_AMY3   213 WYQYKGRDFKVPLIDFSAK-DDNVVQTKVGFAVWPG-RF---PSMVIQSEDAVSEDENNN
Nymphaea_colorata_AMY3     198 WYQHRGGDFRVHFVNSDPA-DGHIDSNDKSFQIWP--VLDQMSNMLLKTKGSQSKGQGEK
Erythranthe_guttata_AMY3   203 WYQHRGRDFKVPLIEHPQI------DVKA----GEG-DVAQLTTVLVKPEAAESKGDDI-
Beta_vulgaris_AMY3         198 WYQHKGRDFVVPLVDNLRD-GDN-VIGKKGLSSWPG-NL---SHMLLEGEAPQSVAEGSG
Daucus_carota_AMY3         201 WYQHKGRDFKISVIDCLPG-NRETIGSKKGSKKWPG-ALGQLSSMILQSEGVDSDGVKSS
Arabidopsis_thaliana_AMY3  201 WYQHKGRDFKVPLVDDVPD-NGNLIGAKKGFG-----ALGQLSNIPLKQDKSS-------
Ipomoea_triloba_AMY3       197 WYQHRGMDFKVPLMDYAHD-DSNMVGGKKGFGIWTG-ALGQLSDMLLKSEVDHTKGENGS
Ricinus_communis_AMY3      199 WYQHKGRDFKVPLVDYLLE-GGNVVGAKRGFSIWPG-SLL--SNMLLKTETLPSKDEDNN
Gossypium_hirsutum_AMY3    200 WYQHRGMDFKVPLVDYLED-DGNTVGAKRGFGVWSG-ALQQFSNVLLKSEASHADSQNNS
Phaseolus_vulgaris_AMY3    210 WYQYKGRDFKVPLVNYLKE-DANIIGPKKGFSLWPG-ALGQISNILLKSDATHDKVQDGN
Cucumis_sativus_AMY3       197 WYQHKGRDFKVPLLDYCGE-DGNKVGTKKGLGLWPG-ALGQLSNLLVKAETNS-KDQGSS
Fragaria_vesca_AMY3        202 CYQHRGRDFKVPLVGYLQEEKGNVVGAKKGLGMLPG-VLGKLTNIFFKAEISNSQEKDSG
Nelumbo_nucifera_AMY3      197 WCQHRGRDYKVPLISYLHE-DANIIGAKKSFGIWPG-ALGQIPSILLKPEKPTHE--EDT
Vitis_vinifera_AMY3        195 WYQHRGRDFEVLLMDYLCE-GTNTVGAKEGFGIWPG-PLGQLSNMLLKAEGSHPKGQDSS
Camellia_sinensis_AMY3     204 WYQHKGRDFKVVLIDNLHE-DGNFVGAKKGLGIWPG-ALGQLSSVLLKSEGAHPKGEDSS
Zea_mays_AMY3              201 WFQHKGRDFRIPLNGSFND------GGKQDIDIWPGD----LGHVLKKSEGSSSQPQNTS
Amborella_trichopoda_AMY3  207 WYQHRGRDFRVCLLEDLQD-ENDKVGDKKSFSLWPG-DFVKMPEVLLTAIKREANGQEPN
Zingiber_officinale_AMY3   205 WFQHKGRDFRISFTDSSNG--GNTVDGSRGFSIWPGGALNQIPGMLLKAEGSTSKQKYSK
Magnolia_sinica_AMY3       205 WYQHRGRDFRVPLINQLDD--GNIVGDKKGFRLWPG-ALDQISTILLKPEGSRPKGEESV
Phoenix_dactylifera_AMY3   201 WFQHKGRDFKISLKDFIEE--ENTSSGKQGFDIWIG-AFDQISSLLVNAEGSSSKPQEAV

                           301 .......310.......320.......330.......340.......350.......360
Erigeron_canadensis_AMY3   268 NNGNLEVSVQQKKILESFCEEHSIVKETLVENSMNVYMKKGSDASNHLVHIETDILDDVI
Nymphaea_colorata_AMY3     255 ----------KMKQNDGIYEEYPILKETIVDGFLSISVQESKEMKKHLVFFDTDLSGDII
Erythranthe_guttata_AMY3   251 -----GGVRLQRGPLQSFYEEFSVVKETVNSNSISVSVKHTEEKDKCVLYIETDLPGDVV
Beta_vulgaris_AMY3         252 --EASPEPTLANRLLEGFYEEQPIVKEVPVCNSLFVSVRKCPETDKNLVSVETDLPGDVV
Daucus_carota_AMY3         259 --NDPKG-TSPRSSLKSFCEEHPIVKETLFDNAVKVSVSTCPETAKKLLYMETDLPGDIV
Arabidopsis_thaliana_AMY3  248 ---AETDSIEERKGLQEFYEEMPISKRVADDNSVSVTARKCPETSKNIVSIETDLPGDVT
Ipomoea_triloba_AMY3       255 --NGSSEPREKTRCLAGFYEEHVIVKETLVDNSVTVSVKEYPETAKNLLQIDTDLPGDVL
Ricinus_communis_AMY3      255 --SETKDVKQDSGQLKGFYEEQPITKQVTIQNSATVSVTKCPKTAKYLLYLETDLPGEVV
Gossypium_hirsutum_AMY3    258 --IESKDSKNKNRCLEGFYEEQSIVKEVSVGNLVSVAVRKSPETGKVIVCLETDIPGDVV
Phaseolus_vulgaris_AMY3    268 --TGSRNTKVENSQLEGFYVELPITKEISVNNSISVSIRKCSETAKNNLYLETDIPGDIL
Cucumis_sativus_AMY3       254 --SESGDTKEEKKSLEGFYKELPIVKEIAVDNSISVSVRKCSETTKYLLYLESDLPGDVI
Fragaria_vesca_AMY3        261 --GESRGTKEQTRSLEGFYEELPIAKEIAVVNSVTVSVRKCPETAKNLLYLETDLLNHVV
Nelumbo_nucifera_AMY3      253 --G-ETDDKKQNKCLEGFYEEHPIFKEVPVQNYMTVSVRKCPDKDKNLIHLDTDLPGDVI
Vitis_vinifera_AMY3        253 --S-V-----SGDLITGFYEEHSIVKEVPVDNSVNVSVKKCPETARNLLYLETDLIGDVV
Camellia_sinensis_AMY3     262 --E-SRYPNQKNKSLEGFYEEHSIVREVLISNSVTVSVRKCPEMAKNLLYMETDLPGDVV
Zea_mays_AMY3              251 ----PEDTGLSGKHISGFYEEYPILKSEYVQNLVTVTVRRDIEAHKRLVEFDTDIPGEVI
Amborella_trichopoda_AMY3  265 --GDGKDARKKAKLIEEFYDEYIFMKEKMVGNYLTVSVQENEEKNKALVLFDTDLPGNVI
Zingiber_officinale_AMY3   263 ----DTKILKHDRIIARFNKEFPILKEEFVSNFMTVSLTRSEDTHQKFVQFDTDLPGDVV
Magnolia_sinica_AMY3       262 --VDSKDAKQQNRRVEGFYEECSILKEEPTQNFVTVSVRNCDETDKNVVRFDTDLPGDVV
Phoenix_dactylifera_AMY3   258 --GEAKVTKRQNKRIQGFYEEYSLLKEEFVQNFMTVTVRKSDESDKNIVQFDTDMPGDVV

                           361 .......370.......380.......390.......400.......410.......420
Erigeron_canadensis_AMY3   328 VHWGICKDESKKWEIPSGPYPDN----TTVFKNKALRTQLQKKEGEDGRSGVFPLDKETA
Nymphaea_colorata_AMY3     305 VHWGVCKDDKKRWEIPSGPHPPK----THIFRKKALRTFLQPKEDGSGSWGLFSLDRKLS
Erythranthe_guttata_AMY3   306 VHWGVCRDESKKWEIPVEPYPPG----TIVFKNKALRTLLQRKNDGDGSGGSFTLGEGLL
Beta_vulgaris_AMY3         310 LHWGVCRDNPKEWEVPAAPHPPN----TEVFKNKAMRTKLQTKEDGFGSKGLFTLDNEIT
Daucus_carota_AMY3         316 VHWGVCKDESKKWEIPAEPHPVK----TTNFKNKALRTILQR-KGEEGAGGLFTMEQGLV
Arabidopsis_thaliana_AMY3  305 VHWGVCKNGTKKWEIPSEPYPEE----TSLFKNKALRTRLQRKDDGNGSFGLFSLDGKLE
Ipomoea_triloba_AMY3       313 IHWGVCRDEGKNWELPAKPYPTE----TTIFKNKALRTSLQQKDDGSGSQRSFTLDEGPV
Ricinus_communis_AMY3      313 LHWGVCRDDAKNWEIPSSPHPPE----TTVFKNKALQTMLQPNDGGNGCSGLFSLDEEFA
Gossypium_hirsutum_AMY3    316 VHWGVCRDDAKIWEIPSAPYPPE----TTVFKNKALRTLLQPKATGNRSGALFTLDEEHF
Phaseolus_vulgaris_AMY3    326 LHWGVCRDDLRWWEIPPTPHPPE----TIAFKDRALRTKLQSRDNGVGSSVQLSLGEELS
Cucumis_sativus_AMY3       312 VHWGACRDDTKKWEIPAAPHPPE----TTVFKNKALRTLLQPKEGGKGCSGVFTIEEDFG
Fragaria_vesca_AMY3        319 VHWGVCKDDSKRWEVPAAPHPPE----TVVFKDKALRTRLQQKEGGNGCWGLFTLEEGPA
Nelumbo_nucifera_AMY3      310 VHWGVCRDDDKKWEIPAAPHPPQ----TQVFKKKALRTLLQPKEDGHGCWGLFSLDREFK
Vitis_vinifera_AMY3        305 VHWGVCRDDSKTWEIPAAPHPPE----TKLFKKKALRTLLQSKEDGHGSWGLFTLDEELE
Camellia_sinensis_AMY3     319 VHWGVCKDDGKKWEIPAEPYPALYYIRTLVFKNKALRTLLK-KKGGHGGSSLFTLDEGYL
Zea_mays_AMY3              307 IHWGVCRDNTMTWEIPPEPHPPK----TKIFRHKALQTLLQQKADGAGNSISFSLDAEYS
Amborella_trichopoda_AMY3  323 IHWGVCRDNGKKWEIPQASHPPS----TNLFRKKALQTSLQFKENGGGSWGLFTLDKELA
Zingiber_officinale_AMY3   319 VHWGVCKADGRKWEIPTTPHPPR----TKIFRQKALQTVLQPKEDGQGNWGLFPIDEETS
Magnolia_sinica_AMY3       320 VHWGVCRDDKKRWEIPAAPHPPQ----TNIFRKKALQTLLQRNENGSGSWGSFPLDKELV
Phoenix_dactylifera_AMY3   316 VHWGVCRDNGKNWEIPPAPHPPA----TKIFRRKALQTLLQPKRNGLGNWGLFLLDKGIS

                           421 .......430.......440.......450.......460.......470.......480
Erigeron_canadensis_AMY3   384 GFLFVLKLKDNTWINN--MGKDFYIPVPKESKLDVEV-----------------------
Nymphaea_colorata_AMY3     361 GLLFVLKVNEYTWVNN--MGNDFYIPLQSQSSLSTLMEEILEEAENEKFVT---T-GESD
Erythranthe_guttata_AMY3   362 GFVFVLKLNENAWLNC--KGSDFYIPLPRTVVPDKLSVSP--------------------
Beta_vulgaris_AMY3         366 GFPFVLKLNDDKWLNN--RGNDYFIPLSGTHNKGGSSAEIEKNVS--SE-----------
Daucus_carota_AMY3         371 GFLFVLKLNDDTWLNC--MGNDFYVPLSTSSDLEKLSVKNQSEDSIQKVL----------
Arabidopsis_thaliana_AMY3  361 GLCFVLKLNENTWLNY--RGEDFYVPFLTSSSSPVETEAAQVSKPK--------------
Ipomoea_triloba_AMY3       369 GFVFVLKLDDGTWLNC--KGNDFYVPLPRSTKGQLSSIESEVETQNKEL-----------
Ricinus_communis_AMY3      369 GFLFVLKLNEGTWLKC--KGNDFYVPLSTSSSLPTQPG----QGQSEGV-----------
Gossypium_hirsutum_AMY3    372 GFLFVLKLDDNTWLKF--KENDFYVPLLGTSSVPGQYG----QSD---------------
Phaseolus_vulgaris_AMY3    382 GFLFVLKLNDGAWIND--MGDDFYIPLPRSSSLIIDNRENQFEGVQREV-----------
Cucumis_sativus_AMY3       368 GFLFVLKQKENSWLNY--KGDDFYIPFPSSGNLSNQQRKSKLKDTRA-------------
Fragaria_vesca_AMY3        375 GFLFVFKLNESTWLKC--KGNDFYIPLSSANKLPAVAKDDHSEGDKVDERSE--------
Nelumbo_nucifera_AMY3      366 ALLFVLKLNENTWLNY--MGCDFYVPLSKANSSPVQSSQSQTEGQGKQDILYLPKSEVSE
Vitis_vinifera_AMY3        361 GFLFVLKLNENTWLRC--MGNDFYIPLLGSSSLPAQSRQGQSEGWGKSERVV--------
Camellia_sinensis_AMY3     378 GFLFVLKLTDNTWLNY--MGNDFYIPLSSSSGLSAISRHGQSEGQVE-------------
Zea_mays_AMY3              363 CLFFVLKLDEYTWLRNLENGSDFYVPLTRVGQYGSTQDPDKA------------------
Amborella_trichopoda_AMY3  379 GLLFVLKLDGYTWLNN--NGSDFYIPLSAEIGTSSVRPTEKINAPEGHK--------EED
Zingiber_officinale_AMY3   375 AVVFVLKLNEYTWLNN--MGADFFVPTGNVINSLAEFGSA-LGTE---------------
Magnolia_sinica_AMY3       376 GLLFVLKLKEYIWMNN--MGSDFYIPLMSEGSFSTQTSRIQEKDV---------------
Phoenix_dactylifera_AMY3   372 GVLFVLKLNEYIWLNN--MGTDFYIPLTSASSSSIQTCQDSIANE---------------

                           481 .......490.......500.......510.......520.......530.......540
Erigeron_canadensis_AMY3   419 -----------------------------NEEGSVYTDEIIDEIRHLLSGISSEGNQKTK
Nymphaea_colorata_AMY3     415 LAITDGALSKDESSFTDGTVLGTSENEDEGATDVQYTNEIINEIRNLVSDISAERSINTK
Erythranthe_guttata_AMY3   400 ------------------LISEN--ASESNQTSSTYTDGIISEIRSLVTDISSEKTRQTK
Beta_vulgaris_AMY3         411 V-LSG-P-----------IRDENPPVENQEAHTTPFTDDIIKEIRNLVTGISSETKWKTK
Daucus_carota_AMY3         419 ---TE-T-----VESDSSPISKK--DGLSDPTISEYTDEIINEIRSLVTGISSEKKRKTK
Arabidopsis_thaliana_AMY3  405 ------------------------RKTDKEVSASGFTKEIITEIRNLAIDISSHKNQKTN
Ipomoea_triloba_AMY3       416 ------------------DSLGASGSTSEAIEASLYTDEIINEIRSLVSDISSEKNRKTK
Ricinus_communis_AMY3      412 ------------------LASGKDAEGNEEVSRTAYTDEIIDEIRNLVNGISSEKVRQTK
Gossypium_hirsutum_AMY3    411 -------------------------ITTEEISSKSYTDGIINEIRNLVSGLSSEKSQKTK
Phaseolus_vulgaris_AMY3    429 --------------------TEVTEEAGEEESISAFTDEIISEIRHLVTDISSEKNRKTK
Cucumis_sativus_AMY3       413 -----------------SKISG---EESEGVSVTAYTDGIIKEIRNLVTDISSQKTKKKK
Fragaria_vesca_AMY3        425 -------------------------EEIEESSFTEFTNGIINEIRTLVSGISSEKSRKTT
Nelumbo_nucifera_AMY3      424 VVINE-R-----DESSSSGISGKMADADKVVAQGGYTDGIINEIRNLVSDISSEKSHKTK
Vitis_vinifera_AMY3        411 --------------SVPTEISGKTAGENEIVSDAAYTDGIINDIRNLVSDISSEKRQKTK
Camellia_sinensis_AMY3     423 --------------------------TNQVASPATYTDEIIDDIRNLVTDISSEKGQIRR
Zea_mays_AMY3              405 -------------------------EAQKIEDKSSQADGLISDIRNLVVGLSSRRGQKAK
Amborella_trichopoda_AMY3  429 ISNDVKNDTWTIEESGSSQLEKSQSGANSPVSRVSYTDEIINEIRSLVSDISSERSANMK
Zingiber_officinale_AMY3   417 -TS---ELVND--------STGQAQVTAEKDNSVTYSYEIIKEIKNLVSDISSERSKGAK
Magnolia_sinica_AMY3       419 -SSDKTNLEGAG--SSFQDIQERTVEANQAIPPGAFTEGIINEIRNLVSDISSGKSTKTK
Phoenix_dactylifera_AMY3   415 ------QLTWTE--PQDMRHSSNNDETNQAVDHAADTDEIIYEIRNLVTDISSGKGRSTK

                           541 .......550.......560.......570.......580.......590.......600
Erigeron_canadensis_AMY3   450 SKEAQEMILQEIEKLAAEAYGIFRSSVQTSPAIIEEDV-DL---------ESPVKINSAT
Nymphaea_colorata_AMY3     475 SKEAHESILQEIEKLAAEAYSIFRSSRPTFSEETIAEVEASK----------EVAVSSGT
Erythranthe_guttata_AMY3   440 NKKVRESILQEIEKLAAEAYSIFRSSIPTVPKIDLAEDEVLE---------PPVKVCSGT
Beta_vulgaris_AMY3         458 SKEAQEDILQEIEKLAAEAYGIFRSSSMTLSEEAVAGFEEIE---------LPVEICSGT
Daucus_carota_AMY3         468 SKETQETILQEIEKLAAEAYSIFRSPIPAFTESDVIEKIE---------PEAPVKICSGT
Arabidopsis_thaliana_AMY3  441 VKEVQENILQEIEKLAAEAYSIFRSTTPAFSEEGVLEAEADK---------PDIKISSGT
Ipomoea_triloba_AMY3       458 TKEAQESILQEIEKLAAEAYSIFRSSVPTFSESALLEAED-L--------KPPVKISSGT
Ricinus_communis_AMY3      454 TKEAQESILQEIEKLAAEAYSIFRSSIPTFTEESVLESEVEK--------APPAKICSGT
Gossypium_hirsutum_AMY3    446 TKEVQESILQEIEQLAAEAYSIFRSSITTVPEEVVSETET-T--------KPAVKISSGT
Phaseolus_vulgaris_AMY3    469 SKEAQETILQEIEKLAAEAYSIFRNSVPTFSEETITESETAVESKTVIFPELPPQVSSGT
Cucumis_sativus_AMY3       453 TKEAQESILQEIEKLAAEAYSIFRSSAPTFTEEIIETPKPVE---------PPVRISSGT
Fragaria_vesca_AMY3        460 SKEAQESILQEIEKLAAEAYSIFRSNVPTFTEETTLESEELT---------PSVKISSGT
Nelumbo_nucifera_AMY3      478 NKEVQEIILEEIEKLAAEAYSIFRSSTPTFLEEAISDAETLK---------PPLKICSGT
Vitis_vinifera_AMY3        457 TKQAQESILQEIEKLAAEAYSIFRSSIPTFSEDAVLE--TLK---------PPEKLTSGT
Camellia_sinensis_AMY3     457 MKEAQESILQEIEKLAAEAYSIFRSSIPTFAEKVVLEAEEIV---------PAAKICSAT
Zea_mays_AMY3              440 NKVLQEDILQEIERLAAEAYSIFRSPTIDSVDESVQLDDTLS----------AKPACSGT
Amborella_trichopoda_AMY3  489 SKDARESILQEIEKLAAEAYSIFRSSIPTFLKELVSEPEIEK---------PQPKICSGT
Zingiber_officinale_AMY3   465 SKEAQESILQEIEKLAAEAYSIFRSSIPSY-AEPVTDSDLLK---------PAVELTSGT
Magnolia_sinica_AMY3       476 NKEDQEKILQEIEKLAAEAYSIFRSSSTIPFVEPVTEDETLE---------PPVKLCSGT
Phoenix_dactylifera_AMY3   467 SKEAQENILQEIEKLAAEAYSIFRSSIPNFVEEYVSDAQYLK---------PAVKLCPGT

                           601 .......610.......620.......630.......640.......650.......660
Erigeron_canadensis_AMY3   500 GSGSEILFQGFNWESNKSGRWYLELEEKAAELAALGFTIIWLPPPTESISAEGYMPRDLY
Nymphaea_colorata_AMY3     525 GTGYEILCQGFNWESNKSGKWYLEMHDKAAELASLGFTVVWLPPPTESVSPEGYMPKDLY
Erythranthe_guttata_AMY3   491 GSGNEIVCQGFNWESHKSGNWYIELQEKASKLSELGFTVIWLPPPTDSVSPEGYMPSDLY
Beta_vulgaris_AMY3         509 GTGFEILCQGFNWESHKSGRWYMELQDKVSELSSLGFTVIWLPPPTESVSPEGYMPKDLY
Daucus_carota_AMY3         519 GSGFEILCQGFNWESNKSGRWYMELQEKAAELASLGFTVVWLPPPTESISPEGYMPKDLY
Arabidopsis_thaliana_AMY3  492 GSGFEILCQGFNWESNKSGRWYLELQEKADELASLGFTVLWLPPPTESVSPEGYMPKDLY
Ipomoea_triloba_AMY3       509 GSGHEILCQGFNWESHKSGRWYLELQEKAELLSSLGFTVVWLPPPTESVSPEGYMPKDLY
Ricinus_communis_AMY3      506 GTGHEILLQGFNWESNKSGRWHMELKEKAAEISSLGFTVIWLPPPTESVSPEGYMPKDLY
Gossypium_hirsutum_AMY3    497 GTGFEILCQGFNWESHKSGRWYMELKEKASEISSLGFTVIWLPPPTESVSPEGYMPKDLY
Phaseolus_vulgaris_AMY3    529 GTGYEILCQGFNWESHKSGRWYMELKEKAAELASFGVTVIWLPPPTESVSPEGYMPKDLY
Cucumis_sativus_AMY3       504 GSGFEILCQGFNWESHKSGRWYMELKEKAAELSSLGFTVLWLPPPTESVSPEGYMPKDLY
Fragaria_vesca_AMY3        511 GTGFEVLCQGFNWESHKSGRWYMELKSKAAELSSLGFTVIWLPPPTDSVSPEGYMPTDLY
Nelumbo_nucifera_AMY3      529 GSGYEILCQGFNWESHKSGRWYMELTERASELSSLGFTILWLPPPTESVSPEGYMPKDLY
Vitis_vinifera_AMY3        506 GSGFEILCQGFNWESNKSGRWYMELSKKVAELSSLGFTVVWLPPPTASVSPEGYMPTDLY
Camellia_sinensis_AMY3     508 GSGFEILCQGFNWESHKSRRWYMELHEKVAELSSLGFTVVWLPPPTDSVSPEGYMPKDLY
Zea_mays_AMY3              490 GSGFEILCQGFNWESHKSGKWYVELGTKAKELSSLGFTIVWSPPPTDSVSPEGYMPRDLY
Amborella_trichopoda_AMY3  540 GTGYEVLCQGFNWESHKSGRWYSELYEKAADIVSLGFTVIWLPPPTESVSPEGYMPKDLY
Zingiber_officinale_AMY3   515 GSGYEILCQGFNWESHRSGKWYAELSAKTVELSSLGFTVVWLPPPTESVSPEGYMPKDLY
Magnolia_sinica_AMY3       527 GSGSEVLCQGFNWESHKSGSWYSILNEKATELSSIGFTVIWLPPPTDSVSPEGYMPRDLY
Phoenix_dactylifera_AMY3   518 GSGFEILCQGFNWESHKSGRWYSELSAKAKELSSLGFTIIWLPPPTESVSPEGYMPKDLY

                           661 .......670.......680.......690.......700.......710.......720
Erigeron_canadensis_AMY3   560 NLNSKYGTVDQLKAVVKKFHQVGIRVLGDAVINHRCAHYQNQNGVWNIFGGLMNWDDRAV
Nymphaea_colorata_AMY3     585 NLNSRYGTMEELKSLVARFHDVGIKVLGDVVLNHRCAHSQNQNGIWNVFGGKLAWDARAI
Erythranthe_guttata_AMY3   551 NLNSRYGSIDQLKVLVKKLHEVGIMVLGDAVLNHRCAQFQNQNGVWNIFGGRLNWDDRAI
Beta_vulgaris_AMY3         569 NLNSRYGTVDELKTVIKSFHQVGIKVLGDAVINHRCAHSKNQNGVWNIFGGRLNWDDRAV
Daucus_carota_AMY3         579 NLNSRYGTVDELKILIEKFHEFDIKVLGDAVLNHRCAHHQNQNGVWNIFGGRLNWDDRAV
Arabidopsis_thaliana_AMY3  552 NLNSRYGTIDELKDTVKKFHKVGIKVLGDAVLNHRCAHFKNQNGVWNLFGGRLNWDDRAV
Ipomoea_triloba_AMY3       569 NLNSRYGTIDELKMIVKRFHEVGIQVLGDVVLNHRCASFRNQNGVWNIFGGRLNWDDRAV
Ricinus_communis_AMY3      566 NLNSRYGSIDELKDLVKSLHRVGLKVLGDAVLNHRCAHFQNQNGVWNIFGGRLNWDDRAI
Gossypium_hirsutum_AMY3    557 NLNSRYGTIAELKELVKSLHEVGMKVLGDVVLNHRCAHFKNQNGVWNIFGGRLNWDDRAV
Phaseolus_vulgaris_AMY3    589 NLNSRYGTVDQLKDVVKSFHEVGIKVLGDVVLNHRCAHYKNQNGIWNLFGGRLDWDDRAI
Cucumis_sativus_AMY3       564 NLNSRYGNIDELKDVVKTFHDVGIKVLGDAVLNHRCAHFKNQNGIWNIFGGRLNWDDRAV
Fragaria_vesca_AMY3        571 NLNSRYGTMDELKETVREFHKVGIKVLGDAVLNHRCAQYQNKNGVWNIFGGRLNWDDRAV
Nelumbo_nucifera_AMY3      589 NLNSRYGSTEELKLVVKCFHQVGIKVLGDVVLNHRCAHYQNKSGVWNIFGGKLNWDDRAV
Vitis_vinifera_AMY3        566 NLNSRYGSSDELKVLVKSFHEVGVKVLGDVVLNHRCAQYQNQNGIWNIFGGRLNWDDRAI
Camellia_sinensis_AMY3     568 NLNSRYGSTDELKGLVKRFHQVNVRVLGDAVLNHRCAEYQNQNGVWNIFGGRLNWDDRAV
Zea_mays_AMY3              550 NLNSRYGSMDELKELVKIFHEAGIKVLGDAVLNHRCAQFQNNNGVWNIFGGRMNWDDRAV
Amborella_trichopoda_AMY3  600 NLNSRYGTIEELKTLVRRFHEVGIKVLGDAVLNHRCAHYKNQNGVWNIFGGRLNWDDRAI
Zingiber_officinale_AMY3   575 NLDSRYGNIDELKQLVKRFHEVGVKVLGDVVLNHRCAHYQNKNGVWNIFGGRLNWDDRAV
Magnolia_sinica_AMY3       587 NLNSRYGNMEELKILVKKFHEVGIKVLGDVVLNHRCAQYQNNNGVWNMFGGKLNWDDRAV
Phoenix_dactylifera_AMY3   578 NLNSRYGSKEQLKDLVKRFHEVNIKVLGDAVLNHRCAHYQNQNGIWNIFGGHLNWDDRAI

                           721 .......730.......740.......750.......760.......770.......780
Erigeron_canadensis_AMY3   620 VADDPHFQGRGNKSSGDNFHAAPNIDHSQEFVRKDLEEWLCWLRKEIGYDGWRLDYVRGF
Nymphaea_colorata_AMY3     645 VADDPHFQGRGNKSSGDNFHAAPNIDHSQEFVRNDIKEWLCWLRKEIGYDGWRLDFVRGF
Erythranthe_guttata_AMY3   611 VADDPHFQGRGNKSSGDNFHAAPNIDHSQEFVRKDIREWMNWLREEIGYDGWRLDFVRGF
Beta_vulgaris_AMY3         629 VADDPHFQGRGNKSSGDNFHAAPNIDHSQEFVRNDLKEWLGWLRKDIGYDGWRLDFVRGF
Daucus_carota_AMY3         639 VADDPHFQGRGNKSSGDNFHAAPNIDHSQDFVRKDLKEWLCWLRKDIGYDGWRLDYVRGF
Arabidopsis_thaliana_AMY3  612 VADDPHFQGRGNKSSGDNFHAAPNIDHSQDFVRKDIKEWLCWMMEEVGYDGWRLDFVRGF
Ipomoea_triloba_AMY3       629 VADDPHFQGRGNKSSGDNFHAAPNIDHSQEFVRKDLKEWLCWLREEIGYDGWRLDFVRGF
Ricinus_communis_AMY3      626 VADDPHFQGRGSKSSGDNFHAAPNIDHSQDFVRQDLKEWLCWLRDEIGYNGWRLDFVRGF
Gossypium_hirsutum_AMY3    617 VADDPHFQGRGNKSSGDNFHAAPNIDHSQEFVRKDLKEWLGWLREEIGYDGWRLDFVRGF
Phaseolus_vulgaris_AMY3    649 VADDPHFQGRGNKSSGDNFHAAPNIDHSQEFVRKDLKEWLLWLREEIGYDGWRLDFVRGF
Cucumis_sativus_AMY3       624 VSDDPHFQGRGNKSSGDNFHAAPNIDHSQDFVRNDIKEWLLWLRKEIGYDGWRLDFVRGF
Fragaria_vesca_AMY3        631 VADDPHFQGRGNKSSGDSFHAAPNIDHSQDFVRKDIKEWLCWLRHEIGYDGWRLDFVRGF
Nelumbo_nucifera_AMY3      649 VGDDPHFQGRGNKSSGDNFHAAPNIDHSQEFVRNDLKEWLCWLREEIGYDGWRLDFVRGF
Vitis_vinifera_AMY3        626 VADDPHFQGRGNKSSGDNFHAAPNIDHSQDFVREDIKEWLCWLRKEIGYDGWRLDFVRGF
Camellia_sinensis_AMY3     628 VADDPHFQGKGNKSSGDCFHAAPNIDHSQEFVRKDLKEWLCWLREEIGYDGWRLDFVRGF
Zea_mays_AMY3              610 VADDPHFQGRGNKSSGDNFHAAPNIDHSQEFVRNDLKEWLCWMRKEVGYDGWRLDFVRGF
Amborella_trichopoda_AMY3  660 VADDPHFQGRGNKSSGDNFHAAPNIDHSQDFVRNDLKEWLNWLRNEIGYDGWRLDFVRGF
Zingiber_officinale_AMY3   635 VADDPHFQGRGNKSSGDSFHAAPNIDHSQDFVRNDLKEWLCWLRKEVGYDGWRLDFVRGF
Magnolia_sinica_AMY3       647 VADDPHFQGRGNKSSGDNFHAAPNIDHSQDFVRKDLKEWLCWLRKGIGYDGWRLDFVRGF
Phoenix_dactylifera_AMY3   638 VADDPHFQGRGNKSSGDNFHAAPNIDHSQEFVRNDLKEWLCWLRKEVGYDGWRLDFVRGF

                           781 .......790.......800.......810.......820.......830.......840
Erigeron_canadensis_AMY3   680 WGGYVKEYMEASKPYFSVGEYWDSLSYTYGEMDHNQDAHRQRIVNWINDTNGTAGAFDVT
Nymphaea_colorata_AMY3     705 WGGYVKDYMNASEPYFAVGEYWDSLCYTYGNMDHNQDAHRQRIVDWINATNGTAGAFDVT
Erythranthe_guttata_AMY3   671 WGGYVKDYMESTEPYFAVGEYWDSLSYTYGEMDHNQDAHRQRIVDWINATNGTAGAFDVT
Beta_vulgaris_AMY3         689 WGGYVKDYIDSSKPYFAVGEFWDSLSYTYGEMDYNQDAHRQRIVDWINATGGSAGAFDVT
Daucus_carota_AMY3         699 WGGYVKDYMEVTEPYFAVGEYWDSLSYSYGEMDHNQDAHRQRIIDWINATSGTAGAFDVT
Arabidopsis_thaliana_AMY3  672 WGGYVKDYMDASKPYFAVGEYWDSLSYTYGEMDYNQDAHRQRIVDWINATSGAAGAFDVT
Ipomoea_triloba_AMY3       689 WGGYVKDYLETSEPYFAVGEYWDSLSYTYGEMDHNQDAHRQRIVDWINATNGTAGAFDVT
Ricinus_communis_AMY3      686 WGGYVKDYMEATEPYFAVGEYWDSLSYTYGEMDHNQDAHRQRIIDWINATNGTAGAFDVT
Gossypium_hirsutum_AMY3    677 WGGYVKDYLEASEPYFAVGEYWDSLSYTYGEMDHNQDSHRQRIVDWINATNGTAGAFDVT
Phaseolus_vulgaris_AMY3    709 WGGYVKDYLEATEPYFAVGEYWDSLSYTYGEMDHNQDAHRQRIVDWINATGGTAGAFDVT
Cucumis_sativus_AMY3       684 WGGYVKDYLDASEPYFAVGEYWDSLSYTYGEMDHNQDAHRQRIVDWINATNGTAGAFDVT
Fragaria_vesca_AMY3        691 WGGYVKDYMDASEPYFAVGEYWDSLSYTYGEMDHNQDAHRQRIIDWINATSGAAGAFDVT
Nelumbo_nucifera_AMY3      709 WGGYVKDYLEATQPYFAVGEYWDSLSYTYGQMDHNQDAHRQRIIDWINATNGTAGAFDVT
Vitis_vinifera_AMY3        686 WGGYVKDYMDASEPYFAVGEYWDSLSYTYGEMDHNQDAHRQRIIDWINATNGAAGAFDVT
Camellia_sinensis_AMY3     688 WGGYVKDYIEASEPYFAVGEYWDSLNYTYGEMDHNQDAHRQRIVDWINDTNGTAAAFDVT
Zea_mays_AMY3              670 WGGYVKDYLEASEPYFAVGEYWDSLSYTYGEMDYNQDAHRQRIVDWINATNGTAGAFDVT
Amborella_trichopoda_AMY3  720 WGGYVKDYLDATEPYFAVGEYWDSLSYTYGEMDHNQDAHRQRIIDWINATNGTAGAFDVT
Zingiber_officinale_AMY3   695 WGGYVKDYIEASEPYFAVGEYWDSLSYTYGEMDHNQDAHRQRIIDWINATNGTAGAFDVT
Magnolia_sinica_AMY3       707 WGGYVKDYLDASEPYFAVGEYWDSLSYTYGEMEHNQDAHRQRIIDWINATNGTAGAFDVT
Phoenix_dactylifera_AMY3   698 WGGYVKDYLEASEPYFAVGEYWDSLSYTYGEMDHNQDAHRQRIIDWINATNGTAGAFDVT

                           841 .......850.......860.......870.......880.......890.......900
Erigeron_canadensis_AMY3   740 TKGILHSAIERCEYWRLCDSNGKPPGVVGWWPSRAVTFIENHDTGSTQGHWRFPGGKEMQ
Nymphaea_colorata_AMY3     765 TKGILHSVLDKCEYWRLCDQKGKPAGVVGWWPSRAVTFIENHDTGSTQGHWRFPAGKEMQ
Erythranthe_guttata_AMY3   731 TKGILHSTLERCEYWRLSDGEGKPPGVMGWWPSRAVTFIENHDTGSTQGHWRFPSGKEMQ
Beta_vulgaris_AMY3         749 TKGILHSVLERCEYWRLSDPKGKPPGVVGWWPSRAVTFIENHDTGSTQGHWRFPGGKEMQ
Daucus_carota_AMY3         759 TKGILHAALERCEYWRLSDEKGKPPGVVGWWPSRAVTFIENHDTGSTQGHWRFPGGREMQ
Arabidopsis_thaliana_AMY3  732 TKGILHTALQKCEYWRLSDPKGKPPGVVGWWPSRAVTFIENHDTGSTQGHWRFPEGKEMQ
Ipomoea_triloba_AMY3       749 TKGILHAAIERCEYWRLSDTKGKPPGVVGWWPSRAVTFIENHDTGSTQGHWRFPGGKEMQ
Ricinus_communis_AMY3      746 TKGILHSALDRCEYWRLSDQKGKPPGVVGWWPSRAVTFIENHDTGSTQGHWRFPNGKEMQ
Gossypium_hirsutum_AMY3    737 TKGILHSALERCEYWRLSDKKGKPPGVVGWWPSRAVTFIENHDTGSTQGHWRFPGGKEMQ
Phaseolus_vulgaris_AMY3    769 TKGILHSALERCEYWRLSDQKGKPPGVLGWWPSRAVTFIENHDTGSTQGHWRFPSGKEMQ
Cucumis_sativus_AMY3       744 TKGILHSALDRCEYWRLSDEKGKPPGVVGWWPSRAVTFIENHDTGSTQGHWRFPGGKEMQ
Fragaria_vesca_AMY3        751 TKGILHAALERCEYWRLSDQKGKPPGVVGWWPSRAVTFIENHDTGSTQGHWRFPRDKEIQ
Nelumbo_nucifera_AMY3      769 TKGILHSALERCEYWRLSDQKGKPPGVIGWWPSRAVTFIENHDTGSTQGHWRFPSGKEMQ
Vitis_vinifera_AMY3        746 TKGILHSALGRCEYWRLSDQKRKPPGVVGWWPSRAVTFIENHDTGSTQGHWRFPGGKEMQ
Camellia_sinensis_AMY3     748 TKGILHAALERCEYWRLSDQKGRPPGVVGWWPSRAVTFIENHDTGSTQGHWRFPGGKEMQ
Zea_mays_AMY3              730 TKGILHAALERSEYWRLSDEKGKPPGVLGWWPSRAVTFIENHDTGSTQGHWRFPYGMELQ
Amborella_trichopoda_AMY3  780 TKGILHSALGKCEYWRLSDQKGKPPGVVGWWPSRAVTFIENHDTGSTQGHWRFPSGKEMQ
Zingiber_officinale_AMY3   755 TKGILHTALERCEYWRLCDEHGKPPGVVGWWPSRAVTFIENHDTGSTQGHWRFPSGKEMQ
Magnolia_sinica_AMY3       767 TKGILHAALERCEYWRLSDQKGKPPGVLGWWPSRAVTFIENHDTGSTQGHWRFPSGKEMQ
Phoenix_dactylifera_AMY3   758 TKGILHTALGKCEYWRLSDQKGKPPGVVGWWPSRAVTFIENHDTGSTQGHWRFPSGKEMQ

                           901 .......910.......920.......930.......940.......950.......960
Erigeron_canadensis_AMY3   800 GYAYILTHPGTPSVFYDHLFSRYKNEISALISLRNRNKINCRSIINITKSERDVYAAIID
Nymphaea_colorata_AMY3     825 GYAYILTHPGTPAVFYDHVFSHLRGEIAALISLRNRAQIHCRSTVQINRAERDVYAATID
Erythranthe_guttata_AMY3   791 GYAYLLTHPGTPSVFYDHIFSDYQNEISALISVRKRNTINCRSRVTIIKAERDVYAAIVD
Beta_vulgaris_AMY3         809 GYAYILTHPGTPAVFYDHIFSNYKHEISTLISVRNRKKINCRSIVNIHKAERDVYAAIID
Daucus_carota_AMY3         819 GYAYILTHPGTPSVFYDHIFSHLNSEISALISIRTRNKINCRSTVNITKAERDVYAAIID
Arabidopsis_thaliana_AMY3  792 GYAYILTHPGTPAVFFDHIFSDYHSEIAALLSLRNRQKLHCRSEVNIDKSERDVYAAIID
Ipomoea_triloba_AMY3       809 GYAYILTHPGTPSVFYDHVFSGYQSEIGSLLSLRKRNKIHCRSTVKITKAERDVYAAMID
Ricinus_communis_AMY3      806 GYAYILTHPGTPTVFYDHIFSHYRSEIASLISLRKRNEIHCRSSVKITKAERDVYAAIIE
Gossypium_hirsutum_AMY3    797 GYAYILTHPGTPAVFYDHIFSHHRSEIANLISVRNRNGIHCRSLVKIVKAERDVYAAIID
Phaseolus_vulgaris_AMY3    829 GYAYTLTHPGTPSVFFDHLFSHYKTEISTLLSIRKRNKIQCRSTVKICKAERDVYAAVID
Cucumis_sativus_AMY3       804 GYAYLLTHPGTPSVFYDHIFSHYKSEIAALISLRKRNKVNCRSVVKIVKAERDVYAAIID
Fragaria_vesca_AMY3        811 GYAYTLTHPGTPAVFYDHIFSHYRSEIAGLISLRNRNKINCRSIVKITKAERDVYAAIID
Nelumbo_nucifera_AMY3      829 GYAYILTHPGTPAVFYDHIFSHYQSEISALISLRHRTKITCRSAVQITKAEREVYAAVID
Vitis_vinifera_AMY3        806 GYAYILTHPGTPAVFFDHLFSHYRSEIASLISLRNRNEIHCRSTIQITMAERDVYAAIID
Camellia_sinensis_AMY3     808 GYAYILTHPGTPAVFYDHIFSHMKSEISELISLRNRNKIHCRSTIKITKAERDVYAAVIE
Zea_mays_AMY3              790 GYAYILTHPGTPAVFYDHIFSHLQPEIAKFISIRHRQKIHCRSKIKILKAERSLYAAEID
Amborella_trichopoda_AMY3  840 GYAYILTHPGTPAVFYDHIFSHYRDEISALIGLRHRKKINCRSTVEIRKAERDVYAATID
Zingiber_officinale_AMY3   815 GYAYILTHPGTPAVFYDHIFSHCQHDISKLISIRNRNKINCRSMLKITKAERDVYAAVID
Magnolia_sinica_AMY3       827 GYAYILTHPGTPAVFYDHIFSHNGHEISELIALRNRKKIHCRSTVNIMKAERDVYAANID
Phoenix_dactylifera_AMY3   818 GYAYILTHPGTPAVFYDHIFSHHQQEISRLISLRRQKKIHCRSTVKITKAERDLYAAEID

                           961 .......970.......980.......990......1000
Erigeron_canadensis_AMY3   860 DKVAMKIGPGHYEPH--SGSQKWSLALQGNDYKVWEAS--
Nymphaea_colorata_AMY3     885 DKVVMKIGPGYHEPP--RGSKNWILSLQGKDYQIWEAL--
Erythranthe_guttata_AMY3   851 EKLIMKIGPGHYEPT--TGAENWSVAVEGRDYKVWELTRK
Beta_vulgaris_AMY3         869 DKVAMKIGPGHYEPP--SESQQWSLAIEGNDYKVWEAL--
Daucus_carota_AMY3         879 EKVAVKIGPGHYEPQHQHGNQNWSLAREGRDYKVWEAS--
Arabidopsis_thaliana_AMY3  852 EKVAMKIGPGHYEPP--NGSQNWSVAVEGRDYKVWETS--
Ipomoea_triloba_AMY3       869 EKLVVKIGPGYYEPP--SGPQNWSVALEGSDYKVWEAS--
Ricinus_communis_AMY3      866 EKVAMKIGPGHYEPP--SG-KNWSMAIEGKDYKVWEAS--
Gossypium_hirsutum_AMY3    857 EKVAMKIGPGYYEPP--SGSQRWSLALEGRDYKVWETS--
Phaseolus_vulgaris_AMY3    889 EKVAMKIGPGQFEPP--SGSQKWSSVLEGRDYKIWEAS--
Cucumis_sativus_AMY3       864 ETVAVKIGPGNFEPP--SGSNGWSLVIEGKDYKVWEVSK-
Fragaria_vesca_AMY3        871 KKVAMKIGPGHYEPP--NGDQKWSKSLEGRDYKVWELAA-
Nelumbo_nucifera_AMY3      889 EKVAMKIGPGYYEPP--GASGRWVLAVEGRDYKVWEAS--
Vitis_vinifera_AMY3        866 EKVAMKIGPGYYEPP--KGQQRWTLALEGKDYKIWETS--
Camellia_sinensis_AMY3     868 QKVAMKIGPGHYEPP--SGPERWSLAIEGRDYKVWEAS--
Zea_mays_AMY3              850 EKVTMKIGSEHFEP---SGPQNWIVAAEGQDYKIWEASS-
Amborella_trichopoda_AMY3  900 DRVTVKIGPGHYEPP--SGSQNWSLIAQGQDYKVWEVL--
Zingiber_officinale_AMY3   875 EKVAVKIGPGHYEPP--NGSKRWNVAAEGVNYRVWETS--
Magnolia_sinica_AMY3       887 GKVTVKIGPGYYEPP--SDRQQWLLVLEGRDYKVWEAS--
Phoenix_dactylifera_AMY3   878 EKVAVKIGPGHYEPS--STPKKWVVAAEGRDYKVWEAS--


```
