## Supplemental Figure 2 for "Functional Characterization of Arabidopsis α-Amylase3 (AMY3): Amylose Specificity and Structural Insights into Its Duplex Carbohydrate-Binding Module"

```
                             1 ........10........20........30........40........50........60
Nymphaea_colorata_GWD1       1 MSSGLGKNLFQKSLVHGLHT-HEEQ-HIKSYP--RTPGSSLCQSSAKPTTTRVTYQLCK-
Beta_vulgaris_GWD            1 MSNLIGQNVSLLL-------LEQHQSKVVNSLGNGISGIPLF-------RTQATSKLRNS
Arabidopsis_thaliana_GWD     1 MSNSVVHNLLNRG-LIRPLN-FEHQNKLNSSV--YQTSTA----------NPA----LG-
Erigeron_canadensis_GWD      1 MSNSVGHSLLKSTVIT-------HQTQ---------IGNTLFQAQAN-----SNLRNSS-
Amborella_trichopoda_GWD1    1 MSNTLGQGLANQVHGQLLLTVSEQQNRA-GYG--V-SCNLPGQN-QSLNIKKKTKSHQK-
Fragaria_vesca_GWD           1 MSSSVSHNLLQSK--------------INPSG--I-PANTLLR---PKCVHQVPAQPRK-
Phaseolus_vulgaris_GWD       1 ----MSQSIFHQTVLCQTQTVAEHQSKVSSFA--V----------------SVNKGKKN-
Erythranthe_guttata_GWD      1 MSNTIGNNLLHQS-LLYPTV-LEHQGRINSST--CIGGNTFFQA-------QATSLTQK-
Cucumis_sativus_GWD          1 MSNSISQNILHQT-LLRRSV-FDNQSKFNASG--T-HKSTLFQ---AALTNQVPGQHWK-
Magnolia_sinica_GWD1         1 MSNTLGHGLIHRT-LCQPTF-LEHQSKI-HSG--I-PCNSSFQVPAASMSNQSACQMRI-
Nelumbo_nucifera_GWD         1 MSNTIGQGLLNQY-LCHPIT-LEHQSKSVCYG--I-SANSLFQALSVS---QKALQIR--
Ipomoea_triloba_GWD          1 MSNSIGNNLLHQR-FLHPVV-LEYKSRIGSTS--V-GGG-------SLFQSHAISHIRK-
Daucus_carota_GWD            1 MSNSIGSNFVHQR-LLEHQ------TRIDTCV--S-GGN------YTLFRVQDSFRIRK-
Gossypium_hirsutum_GWD       1 MSNSVGQNLIQQH-FLRPTV-LEHQSKLKGSS--GIASNSLC---ATASLNQSLAQPRK-
Ricinus_communis_GWD         1 MSNSISHNLLQQS-LVRHSVVLEHRNKLNSSS--S-SSAA-----ASGIASLSAPQIRR-
Vitis_vinifera_GWD           1 MSNTIGHNLLHKS-LLRHTL-LEHQSKISCSG--V-SGNA-------LFQAQSPTQIKK-
Camellia_sinensis_GWD        1 MSNSIGHNLVHQS-LLGPTV-LEHQSKINSSC--L-SGNT-------LFHAQATSQVRK-
Zea_mays_GWD                 1 MS---GFSAAANAAAAERCALAFRARPA---------------ASSPAKRQQ---QPQPA
Phoenix_dactylifera_GWD      1 MSDALGHSLPK-EALRRPCAIENRGRAHPG-----LSGSFLCGVPSGSHRNQ-------K
Zingiber_officinale_GWD      1 MNNCVGHTLPQ-QALFRPSVVERHNTACQR-----SSGNILCIVPSASKAED---VPSHK

                            61 ........70........80........90.......100.......110.......120
Nymphaea_colorata_GWD1      56 PLVSCKFLENTNLTKGLSPSQG--RGSLVSHVPCAVLATNPAS--EQRVKFNLDGRSELQ
Beta_vulgaris_GWD           47 TIFSKQFYGKSFKSYTNSKKLMA--SRRFSYVARAVLAADHAS--QQADRYNLDGDIELQ
Arabidopsis_thaliana_GWD    42 KIGRSKLYGKGLKQAGRSLVTETGGRPL-SFVPRAVLAMDPQ----AAEKFSLDGNIDLL
Erigeron_canadensis_GWD     39 ALLSTDFRGHRLTIRKSALHTFRL---HSLPTVQAVLATDPTS-EQLGEKYILEGDIEMQ
Amborella_trichopoda_GWD1   55 PLVSDIFPVNHLYVKRKYLSHE--IRRLISFVPRAVLATDPVS--QSAGKFDLDGRSELQ
Fragaria_vesca_GWD          40 SSLSKKFCGNSLQ--KPKLAMGSR-RPV-TFVPCAVLTTDQRSDQQLASKYNLDGNIELQ
Phaseolus_vulgaris_GWD      38 LGLRTSFRGNRLCVRKCKLAMGKH-RHV-DAIPRAVLTTNPAS--ELSGRFILGGNIELQ
Erythranthe_guttata_GWD     49 SSISTEFLGNRLKVRRHKLKMGKCCSSSRSRSTRAVLAADPSS--GLTEKFNLYQNVELQ
Cucumis_sativus_GWD         52 SPISTKFLGNGLNVKKPRMATGTGCRSF-PVNTRAVLATDPAS--ELAAKFKLDENIELQ
Magnolia_sinica_GWD1        54 SLVSTKFHGNTLNKAKAKVSRGRH-SGV-SIVPRAVLAADPAS-E-VVGKFNLDGDSELQ
Nelumbo_nucifera_GWD        51 KRLSTKFRENSLSRTKSIVIKE----KR-SMVTRAVLTTDPAS-EQIKGKFNLDGSSELK
Ipomoea_triloba_GWD         48 SPLSTEFHGKRLTVKKKKLLMQQQ-RAF-SVS-RDVLATVL-S-SEIAEKFNLDGDIELQ
Daucus_carota_GWD           44 LPLTATFRVN---VARKRLSMSSN-RGG-GVMTRAVLATDSPS--ELAEKFNLEGNTELQ
Gossypium_hirsutum_GWD      53 YQISTKFYGNSLSKRKHKLAMGSQ-RPL-AFIPQAVLATDPAS-E-NLGKFNIDGNIELQ
Ricinus_communis_GWD        51 SSISSSFYGNRLKISKSKLAIGTP-RPA-TITPRAVLAMDPAS--ELVGKFKLDGNSELQ
Vitis_vinifera_GWD          48 SPISTKFRGNRLNLRKTKLPMGTH-HLV-SVIPRAVLTTDTTS-EQLAGKFCLDKNIELQ
Camellia_sinensis_GWD       48 LPLSTEFRGRSLDVRKQKLLMGTP-RPN-SLFPRAVLTTGTAS-TQLAGKFKLDGNAELQ
Zea_mays_GWD                40 SL---RRSGGQR---RPTTLSASS--RGPVVPRAVATSADRAS-PDLIGKFTLDSNSELQ
Phoenix_dactylifera_GWD     48 PLLATRFLGNNLRTARTKLSEQTR--RAVSALTRAVLAADPAS-EELSGKFNLDSDSELQ
Zingiber_officinale_GWD     52 SFLSSRFLGKTPYVGKGNPLKQNL--RTVAMSPQALLAADPAS--ELARKFKLNANSELE

                           121 .......130.......140.......150.......160.......170.......180
Nymphaea_colorata_GWD1     112 IEVIDSSPGLSSRIDIQVSNSN-LSLLLHWGVIRKGHKNWTLPPRRPDGTKVYKHRALRT
Beta_vulgaris_GWD          103 VTVSPLSSGCISKVEILVTNSHDDNLVLHWGGIRDGKEKWILPSRRPEGTRVHKDRALRT
Arabidopsis_thaliana_GWD    97 VEVTST---TVREVNIQIAYTS-DTLFLHWGAILDNKENWVLPSRSPDRTQNFKNSALRT
Erigeron_canadensis_GWD     95 VDVKIS---SIPSVEIRLTNST-DHLYLHWGGLRNINEKWLLPGRRPEGTKVYKDRALRT
Amborella_trichopoda_GWD1  111 ISVDESNPGSLFQINIQVTNSS-PSLTLHWGTIHDGQQNWKLPSRHPEGTQNYKNRALRT
Fragaria_vesca_GWD          96 VYVDASSPGS-TKVDIHVSNSG-DSLVLHWGGIQDRKENWVLPSRHPDGTTVYKKKALRT
Phaseolus_vulgaris_GWD      94 VSVSSAQPGAATQVDIKVSYSS-GSLLLHWGVVCDQPGKWVLPSRRPEGTKVYKNKALRT
Erythranthe_guttata_GWD    107 VDVELPPSGSTSVVNIQVTSGI-DSLLLHWGAIKSHTKKWILPHHRPIGTTVYMDQALRS
Cucumis_sativus_GWD        109 VDVSAPTSGSIRRVNILVTNIG-GSLLLHWGAIRDRKDTWALPSHCPDGTQVYKNRALRT
Magnolia_sinica_GWD1       110 IGVSTPTSGSLSQVDIQVTNSN-NSLILHWGAIRDKNKNWARPSRRPDGTKVYKNKALRT
Nelumbo_nucifera_GWD       105 IDVSSPTQGSRFRIDFQVTNSS-NSLILHWGGISDGQKNWVLPSRWPDGTKMYKNKALRT
Ipomoea_triloba_GWD        103 VDVKPPTSGHITVVDFKVTGDI-DGLFLHWGAVKSGKDKWILPHRRPDGTKDYKNKALRT
Daucus_carota_GWD           97 VNVRSPGPGCLSQIELQVTNSS-DNLVFHWGGIQDKNGKWVLPSRHPDGTKVYKKRALRT
Gossypium_hirsutum_GWD     109 VDASAPTSGSITNVNFRVMYTS-DSLLLHWGAIRGSNDKWVLPSRQPEGTRNHKNRALRT
Ricinus_communis_GWD       107 VSVS--NAGSITQVNFQISYGS-DSLLLHWGGIRDRKEKWILPSRCPDGTKNYKNRALRS
Vitis_vinifera_GWD         105 VDVSVPTPGSMVQVNIQVTNCS-NSLLLHWGAIRDTKGKWVLPSHSPDGTKVYKNKALRT
Camellia_sinensis_GWD      105 VDVSAPVQGSTLQIDMLVTNSS-GTLLLHWGGIQNRKEKWVLPTRHPAGTRVYKNKALRT
Zea_mays_GWD                91 VAVNPAPQGLVSEISLEVTNTS-GSLILHWGALRPDKRDWILPSRKPDGTTVYKNRALRT
Phoenix_dactylifera_GWD    105 IAVRSPSRGSLVQIEIRVTNSS-GSLVLHWGVIRQGRKDWFLPSSRPDGTKVYKNRALRT
Zingiber_officinale_GWD    108 VSICRPTSESPIQIDFQVAYDI-GSLVLHWGVIRQTRREWSLPSHYPEGTKLYKNRALRT

                           181 .......190.......200.......210.......220.......230.......240
Nymphaea_colorata_GWD1     171 PLKKSGDDSFLSLEVDDPEIEFVEFLLVDEAQNKWFKKDGGNFRVQLARSGS--------
Beta_vulgaris_GWD          163 PFQKSGSGSSLNIEIDDPAMQAIEFLIFNEARNIWFKNDGQNFHVRLPEKAE--------
Arabidopsis_thaliana_GWD   153 PFVKSGGNSHLKLEIDDPAIHAIEFLIFDESRNKWYKNNGQNFHINLPTERN--------
Erigeron_canadensis_GWD    151 PFIKSGSNSFLKVEIDDPSIQSLEFLIVDEAKNKWYKNNGQNFHVKLPSREK--------
Amborella_trichopoda_GWD1  170 PFVKSGENSFLKIEVDDPQIKAIEFLLFDESQNKWFKNNGQNFQVRLVSDVR--------
Fragaria_vesca_GWD         154 PFAKSGPNSSLKIEIDDPAIQAIEFLIVDESQNRWFKNNGGNFHVKLPLKEM--------
Phaseolus_vulgaris_GWD     153 PFMKADSESFLRIEIHDPAAQSIEFLILDEAKNKWFKNNGENFHIKLPVKNK--------
Erythranthe_guttata_GWD    166 PFEKSGSNAVLRIEIDDPSIQALEFLVFDEAQNKWYKYSGGNFHVKLPKRES--------
Cucumis_sativus_GWD        168 PFLNSGSNSTLTIEVDDPAIEAIEFLLLDEARNKWYKNNDKNFHVKLPVKEK--------
Magnolia_sinica_GWD1       169 PFRKSGPNSSLTIEIDDPEIQAIEFLILDESRNKWFKNNGENFLVQLTKDKVIPN-----
Nelumbo_nucifera_GWD       164 PFVKSGPDSFLKMEIDDPKIQGIEFLILDESRNKWFKDNGENFRLLLSRKKN--------
Ipomoea_triloba_GWD        162 PFVKSGSNAMVRLEIDDPAIQAIEFLIFNEVQNKWIKSNGDNFHVKLSPRAI--------
Daucus_carota_GWD          156 PFVKSGSNSFLRLEIDDPAIQAIEFLIFDEAQNKWFKHKGDNFHIKLPSDEK--------
Gossypium_hirsutum_GWD     168 PFVKSGSSSYLKLEIDDPQIQAIEFLIFDEARNKWIKNNGQNFHVKLPQRKT--------
Ricinus_communis_GWD       164 PFVKSGSSSYLKIEIDDPAIQALEFLVLDEGQNKWFKYKGQNFHVKLPEREK--------
Vitis_vinifera_GWD         164 PFVKSGSKSILKIEVDDPAIQAIEFLIVDETQNKWFKNNGENFSVKLPVKGK--------
Camellia_sinensis_GWD      164 PFVRSGSNSILKIEIDDPIIQAIEFLIVDEEQNKWFKNNGENFHLKLPVKDK--------
Zea_mays_GWD               150 PFVKSGDNSTLRIEIDDPGVHAIEFLIFDETQNKWFKNNGQNFQVQFQSSRHQGTGASGA
Phoenix_dactylifera_GWD    164 PFVKSGSDSSLTIEIDDPNIQSLEFLVFDGEQNRWFKNNGQNFQVQLFGKGHGKQNAS--
Zingiber_officinale_GWD    167 PFTKVGSISSLRLKIDDPEIEIVEFLILDEAEKKWYKHNGRNFQVHLLKQGNQNPHVS--

                           241 .......250.......260.......270.......280.......290.......300
Nymphaea_colorata_GWD1     223 --SKNLSSIPEELVQIQAYLRWERKGKQAYAPEQEKEEYEAARRELMDEVAKGVTIDDLR
Beta_vulgaris_GWD          215 --HAQQVSVPEELVQVQAYLRWERKGKQMYTPYQEKEEYEAARAELLEEVAKGASIQDLQ
Arabidopsis_thaliana_GWD   205 --VKQNVSVPEDLVQIQAYLRWERKGKQMYNPEKEKEEYEAARTELREEMMRGASVEDLR
Erigeron_canadensis_GWD    203 --QVSSVVIPEDLVQIQAFLRWEKKGKQMYNPEQEKVEYEAARTELYDEIAKGSSLEDIR
Amborella_trichopoda_GWD1  222 --TAQNISVPEDLVQVQAYLRWERKGKQMYTPEQEKEEYEAARTELLEEVARGTAIDELR
Fragaria_vesca_GWD         206 --SISNVSVPEDLVQIQAYLRWERKGKQMYTPEQEKVEYEAARTELLEEVARGASIQELQ
Phaseolus_vulgaris_GWD     205 --LSQEVSVPEDLVQIQAYLRWERKGKQMYTPEQEKVEYEAARQELLEEVSRGTSVQDLR
Erythranthe_guttata_GWD    218 --RSPNVSIPEELVQVQAYLRWERNGKQNYSPEKEKEEFEAARRELLEEISRGASIQDLR
Cucumis_sativus_GWD        220 --FISDVSVPEELVQIQAYLRWERKGKQTYTPQQEQEEYEAARAELLQELTRGATLQDLR
Magnolia_sinica_GWD1       224 VLVPPDVSVPEDLVQIQAYLRWERKGKQAYTPEQEKEEYEAARSELFQEVARGTSIEQLR
Nelumbo_nucifera_GWD       216 --MSPHVSVPEDLVQIQAYLRWERKGRQMYTPDQEKKEYEAARIELMEEIAKGVSVEELR
Ipomoea_triloba_GWD        214 --QITNASVPEDLVQIQAYLRWERKGKQMYSPEQEKVEYEAARAELLEEITKGASVEEIR
Daucus_carota_GWD          208 --LSPNVSVPEDLVQIQAYLRWERNGKQMYTPEQEKEEYEAARTELLVEVSRGTPIQDIR
Gossypium_hirsutum_GWD     220 --LVSNISVPEDLVQVQAYLRWERKGKQMYTPEQEKEEYEAARAEILEEISKGASVDDIR
Ricinus_communis_GWD       216 -VMIQNVSVPEELVQVQAYLRWERKGKQIYTPEQEKEEYDAARVELLEELARGTSVEDLR
Vitis_vinifera_GWD         216 --MIPNASVPEELVQIQAYLRWERKGKQMYTPEQEKEEYEAARTELVEEIARGTSIEDMR
Camellia_sinensis_GWD      216 --LIPKISVPEDLVQIQAYLRWERKGKQTYTPEQEKEEFEAARFELLEEIARGTSVQDLR
Zea_mays_GWD               210 SSSATSTLVPEDLVQIQAYLRWERRGKQSYTPEQEKEEYEAARAELIEEVNRGVSLEKLR
Phoenix_dactylifera_GWD    222 VSGNPNVDLPEELVQIQAFLRWERKGKQTYTPDQEKEEYEAARAELLEEISRGTSIKELR
Zingiber_officinale_GWD    225 VSGNPNIIVPEDLVQIQAYLRWERKGRQAYTPDKEKEEYEAARMELLQEISRGMSIEELR

                           301 .......310.......320.......330.......340.......350.......360
Nymphaea_colorata_GWD1     281 NKLIRNDSTSKDT-----------LVPKSAIPDDLIQIQAYIRWEKAGKPHYSKDKQLME
Beta_vulgaris_GWD          273 ARLRKGSNASEVTKETP-------SNSKTQIPDDLVQIQAYIRWENNGKPNYSPDQQSKE
Arabidopsis_thaliana_GWD   263 AKLLKKDNSNESPKSNGT------------------------------------------
Erigeron_canadensis_GWD    261 TRLTKKDGKSSEQGT---------------------------------------------
Amborella_trichopoda_GWD1  280 AKLTSNSDTLKDPLDPLG------KVLVEKIPDDLIQIQAYIRWEKAGKPNYSQDQQIKE
Fragaria_vesca_GWD         264 ARLTKKSDSGNSHEQSQAGN----------------------------------------
Phaseolus_vulgaris_GWD     263 ARLTKNTKAAEVKEPSVS--------ETKTIPDELVQIQSYIRWEKAGKPNYSQEQQLME
Erythranthe_guttata_GWD    276 TKLTGKKDTSESKEQLV-------SGSKSSIPEDLVQIQSFIRWERAGKPNYSPEQQFKE
Cucumis_sativus_GWD        278 ARLTKENDGTETMELS--------TPKDMTIPDELAQIQAYLRWEKAGKPNFSPEQQLRE
Magnolia_sinica_GWD1       284 AKVMRKDDNS-KK--PI-------VPESNKIPDDLVQVQSYIRWEKAGKPNYSPDQQLME
Nelumbo_nucifera_GWD       274 AKLTKKDESK-AKEPTV-------LESKRKIPDDLVQIQAYIRWEKAGKPNYPPDKQIKE
Ipomoea_triloba_GWD        272 ARLTKKNDTTEHKEQHH-------PETKSDIPDDLVQIQAYIRWEKAGKPSYSPEKQLKE
Daucus_carota_GWD          266 ARLTNKSNVNESKGPLL-------SEKISQVPDDLVQTQAYIRWEKAGKPNFSPDQQLKE
Gossypium_hirsutum_GWD     278 SKITKKSGQE-YKE-------TAINEENNKIPDDLVQIQAYIRWEKAGKPNYSPEQQLRE
Ricinus_communis_GWD       275 TRLTNRNDRHEIKEPPV-------AETKTKIPDDLVQIQSYIRWEKAGKPSYSPEQQLRE
Vitis_vinifera_GWD         274 TRLTNESAKSEIKEQPH-------SETKSKIPDELVQVQAYIRWEKAGKPNYTPDQQLRE
Camellia_sinensis_GWD      274 ARLTSKNDTSESKEQQPSETKKQLSETKGQVPDNLVQLQAFIRWEKAGKPNFSADEQLKE
Zea_mays_GWD               270 AKLTKAPEA-PESDES---KSSASRMPIGKLPEDLVQVQAYIRWEQAGKPNYPPEKQLVE
Phoenix_dactylifera_GWD    282 AKLTKT----PDAEED---RTKRSPSTEGEIPKDLVQVQAYIRWEKAGKPNYPPEMQLRE
Zingiber_officinale_GWD    285 SKLTKKTEVKPASREE---KTHRVESHEGGISDDLVQVQAFIRWEKAGKPNYPPEKQLME

                           361 .......370.......380.......390.......400.......410.......420
Nymphaea_colorata_GWD1     330 FEEAKKELQRELDKGVSLDELRRKIMKGDIDTKVSKQLKTKGRFYIERITRKKRDILNLV
Beta_vulgaris_GWD          326 FEEARKDLQRELDKGTSLDDIRRKIAKGEVKTKVAKQLEKKSYFSADRIQRKKRDLMQIL
Arabidopsis_thaliana_GWD   281 -----------------------SSSGREEKKKVSKQPERKKNYNTDKIQRKGRDLTKLI
Erigeron_canadensis_GWD    276 -----------------------GILEKEIQPKVQKQVQEKSY-SIQEIQRKKRDIMQLL
Amborella_trichopoda_GWD1  334 FEEARKELQNELDKGMSLDEIRKKIVKGNIQTKVTKQLKNKKYFTVERIQRKKRDIMQLL
Fragaria_vesca_GWD         284 ----------------------------KEVSKAAKKHESKQVYRTGRINRKQRDLMKIL
Phaseolus_vulgaris_GWD     315 FEEARKELSAELEKGASLDEIRKKIIKGEVQTKVAKQLKTKTYFRAERIQRKNRDLRQII
Erythranthe_guttata_GWD    329 FEEARKELQVELDKGASLDEIRKRITKGGTQAKVSKQPEKKNYSTGERIQRKKRDVMSLL
Cucumis_sativus_GWD        330 FEEAKKELLSELNKGASIDEIRKKITKGEIKTKVAKQLQDKKYFRVDKIQRKTRDLVQLV
Magnolia_sinica_GWD1       334 FEEARKELQLELNQGTSLDEIRKKIVKGSIQTKVSKQLEKKKYFTAERIQRKKRDIMQFL
Nelumbo_nucifera_GWD       326 LEEARKELQMELDKGTSLEEIRNKIVKGEIQTKVSKQLKNKNYFTIERIHRKKRDFMQYL
Ipomoea_triloba_GWD        325 FEEARQELQLELEKGVTLEELRKRIVKGEIKTKVAKQLAKKSYFTIEKIQRKQRDLAQII
Daucus_carota_GWD          319 FEEARKELQTELEKGISIEAIRKKITKGEIKTNVSKIPDTKRYNAVGRIQRKKRDLMQLL
Gossypium_hirsutum_GWD     330 FEEARKELQSELEKGASLDEIRKKITKGEIKTKVAKQLQNKKYFSPERIQRKQRDLMQLL
Ricinus_communis_GWD       328 FEEARQDLQREVKRGVSLDEIRKKIAKGEIQSKVSKQLQKQKYVSSEKIQRKRRDLAQLI
Vitis_vinifera_GWD         327 FEEARKDLQTELEKGLSLDEIRKKMIKGEIQVKVSKQQKSRRYFGVERIQRKKRDLMQLL
Camellia_sinensis_GWD      334 FEEARKELQMELDKGTSLDEIGNKITKGEIQTKVSKKLETKKYFTVQRIQRKKRDLMHLL
Zea_mays_GWD               326 FEEARKELQAEVDKGISIDQLRQKILKGNIESKVSKQLKNKKYFSVERIQRKKRDITQLL
Phoenix_dactylifera_GWD    335 FEEARKELQLELDKGTPLAELRKKIMKGDIQTKVSKQLKTKKYFTVERIQRKKRDIMQLL
Zingiber_officinale_GWD    342 FEEARKELQLELDKGTSLAELREKIMKGSISTKVLKQLKAKKYFSIERIQRKERDIMEIL

                           421 .......430.......440.......450.......460.......470.......480
Nymphaea_colorata_GWD1     390 NKFTSEAVSRK------HVVEPKAPSLLELCSKSKEDQDAGHTLSKKIYEIENGELLVLV
Beta_vulgaris_GWD          386 SKKAAEPEVKV----STKVTLPKALSTVEHFVEAKEAEDGGLVVVKKIFELADKQIMVLV
Arabidopsis_thaliana_GWD   318 YKHVADFVEPE----SKSSSEPRSLTTLEIYAKAKEEQETTPVFSKKTFKLEGSAILVFV
Erigeron_canadensis_GWD    312 SKHIPVSVKSA---DENLSVKPKTLSAIELYSKAIEEQSDIDIVNKKTYKLADEELLVLV
Amborella_trichopoda_GWD1  394 NKHAAESLKTE------VSVMPRAPTTLELCSKVKEEQDGGCVLNKKVFKFGDKELLALV
Fragaria_vesca_GWD         316 NKHTTKPVDEA-KLANEESAKPKALKAVELFAKEKEEQDGASTLNKQIYKLGDKDLLVLV
Phaseolus_vulgaris_GWD     375 NRIVDENIVEQ------FIDVPKSLTVIEHYAKEREENESGPVLNKTIYKLDDNDLLVLV
Erythranthe_guttata_GWD    389 SKFDSVPVEEK------ISLEPPVLSAIKQFAREKEDHIDGPIVNKKIYKLADKELLVLV
Cucumis_sativus_GWD        390 NQYKSQPIEET------YTAKPKALTEFEKFAKIKEEQDGDDVINKIIYKLGDKDLLVLV
Magnolia_sinica_GWD1       394 NKYATEAVVKE-----KTPPTPQVPTVLELWSQAKEGQDGGRIMNKKIFKLGDKELLVLV
Nelumbo_nucifera_GWD       386 NKHAAESVKD-------LPVQLRALTTIEIFSKAKEEQDGGVILNKKIFKLGDKELLVLA
Ipomoea_triloba_GWD        385 NRNVPWSGSESGSWVEQILSEPQSLSTIELFAKAKEEQVDGPILNKKIYKVANSELMVLV
Daucus_carota_GWD          379 NKYTPGTIKET------IPAKPPTLSSMDLFAKAKEEQSGGPALRKNVYKLANKELLVLV
Gossypium_hirsutum_GWD     390 NKHAVKVVEES----ISVEVEPKPLTAVEPFAKEKE-LDGSPVMNKKIYKLGEKELLVLV
Ricinus_communis_GWD       388 TKYAATPVEE------PVSSEPKALKAIELFAKAKEEQVGGAVLNKKMFKLADGELLVLV
Vitis_vinifera_GWD         387 HRHVTEWTEEK----TPIPIKKTELTAVEQFAKLKEEQDSGSVLNKKIYKISDKELLVLV
Camellia_sinensis_GWD      394 NKYSAGSVEEK------ISVKPRALSTIERFAMKMEEKGGVPIMNKKIFRLADKELLVLV
Zea_mays_GWD               386 SKHKHTLVEDK------VEVVPKQPTVLDLFTKSLHEKDGCEVLSRKLFKFGDKEILAIS
Phoenix_dactylifera_GWD    395 NKHAPEIIEQK------ISDLPKASKVLEPCLKPIEEQDGGLILNKKHFKLDDKGLLVLV
Zingiber_officinale_GWD    402 NKNVAETLDEK------TSQIAPPLTVLELLAKSIHEQDGESVLNQKIYKLDNKDLLVLV

                           481 .......490.......500.......510.......520.......530.......540
Nymphaea_colorata_GWD1     444 TKLNDKMKVYVATDVKEPLVLHWGLAK-SAGQWMVPPASNLPPQSLLQDGSCETHFSEGF
Beta_vulgaris_GWD          442 TKFADKTKISLATDFSEPLTLHWALSR-KGNEWLTPPQNVLPPGSVLLEKAAETQFARVS
Arabidopsis_thaliana_GWD   374 TKLSGKTKIHVATDFKEPVTLHWALSQ-KGGEWLDPPSDILPPNSLPVRGAVDTKLTITS
Erigeron_canadensis_GWD    369 TKASSKIKVHLATSVKEPLTLHWALSK-KAGEWLAPPESLLPQGSISLAAAADTHFSTLS
Amborella_trichopoda_GWD1  448 TNPNGKIKIYLATDLKGPVTLHWGLSK-RAGEWMAPPPGIIPPGSTLEQKASETQFVEGF
Fragaria_vesca_GWD         375 TKVADKIRVHLATDIKEPVNLHWALSVNRAGEWLEPPPNVLPQGSVSLNGAVETQFVSSS
Phaseolus_vulgaris_GWD     429 TKDAGKIKVHLATNSKKPLTLHWALSR-TSEEWLLPPGNSLPPGSVTMNEAAETPFKAGS
Erythranthe_guttata_GWD    443 AKSSGKTKVYLATDLPEPVVLHWALSK-IPGEWTAPPATVLPLDSVSLDKAAETKLAIIS
Cucumis_sativus_GWD        444 TKTSSKTKVYLATDLQQPITLHWGLSRTNAGEWLTPPPDVLPPGSVSLSQAAETQFIFND
Magnolia_sinica_GWD1       449 TRTSGKTKVYVATDLQEPLVLHWALSQ-KTGEWKAPPPSILPPGSVLLDKASETQFVEES
Nelumbo_nucifera_GWD       439 TKPSDKTKVYLATDLKESLTLHWALSR-NGGDWETPPQSALPQGSVPLGNAVETQFAETY
Ipomoea_triloba_GWD        445 TKPSGKVKVYLATDLNEPAILHWALSK-NAGEWLAPPENDLPPGSTSSDKYAETLFSTS-
Daucus_carota_GWD          433 TKPSGKTRIDIATDFKDPLTLHWALSE-NPGEWLAPPPSTLPAGSVLLDKAVETQFSSGS
Gossypium_hirsutum_GWD     445 TKPAGKIKIHLATDLEEPLTLHWALSE-KDGEWLAPPPAVLPPGSVSLEKAAESKFSTST
Ricinus_communis_GWD       442 TKPPGKTKIYVATDFREPVTLHWALSR-NSREWSAPPSGVLPPGSVTLSEAAETQLTNVS
Vitis_vinifera_GWD         443 TKPAGKTKVYFATDSKEPLTLHWAVSK-KAGEWLAPPPSVLPLDSISLNGAVQTQFVNSS
Camellia_sinensis_GWD      448 MKPAGKTKVHLATDLTEPVTLHWALSK-KPGEWLAPPPSLLPPGSTSLDKAIQTKFISS-
Zea_mays_GWD               440 TKVQNKTEVHLATNHTDPLILHWSLAK-NAGEWKAPSPNILPSGSTLLDKACETEFTKSE
Phoenix_dactylifera_GWD    449 TNAVGKTKVYLATDWKGPLILHWGLSK-KAGEWMAPPSSMLPPGSVLLDKSCQTPFGEAF
Zingiber_officinale_GWD    456 TKPFGRTKVYFTTDQSEPLILHWGLSR-KSGEWMVPPTSSIPPGSLLLEKSCETPFTKGL

                           541 .......550.......560.......570.......580.......590.......600
Nymphaea_colorata_GWD1     503 SGEPS----LKSIEIEMEDGDFVGMPFVLRSGEKWLKTNDSDFYVPFAAESRKAKKD---
Beta_vulgaris_GWD          501 PDDPLHQLQVQSVEIEIEGEDFIGMPFVLVSAGNWIKDNGSDFYINIGVQPK--QKD---
Arabidopsis_thaliana_GWD   433 -TDLP--SPVQTFELEIEGDSYKGMPFVLNAGERWIKNNDSDFYVDFAKEEKHVQKD---
Erigeron_canadensis_GWD    428 IDGAT--NKIQSLEIEVGEGSFVGMPFVLFSGKRWIKNNNSDFYVEFVA-PKKTIMD---
Amborella_trichopoda_GWD1  507 SGDLS----LQSVEIEIGDDQYVGMPFVLQSGGQWIKSNDSDFYIELGVGKEK-KKD---
Fragaria_vesca_GWD         435 -VDST--YEIQSLEIETQEKSFKGMPFVLCSSGNWIKNQGSDFYADFGVAVKKVQKD---
Phaseolus_vulgaris_GWD     488 LSHPS--FEVQSLDIEVDDDTFKGIPFVILSEGKWIKNNGSNFYIEFAG-KKQIRKD---
Erythranthe_guttata_GWD    502 TDNQP--DKVQSLEITIEDESFVGMPFVLLSGEKWVKNGGSDFYVELNTGSVK-KKD---
Cucumis_sativus_GWD        504 -DGST--LKVQYLEILIEEDGFLGMSFVLQSSGNWIKNKGSDFYVAFAIQPKKVRKV---
Magnolia_sinica_GWD1       508 YGDTS--YQVKFVEIELDEGDFKGMPFVLLSSGNWIKNNGSDFYVEFSSRIEKAKKD---
Nelumbo_nucifera_GWD       498 CGDPP--QQVQALEIEIE-DNFVGMPFVLVSNGNWIKNNGSDFYVDFNTESKKVKKD---
Ipomoea_triloba_GWD        503 SDNLS--YKVQSLEIAIEDEDFVGMPFVLLCGEKWIKNNGSDFYVDFRTQP---RRD---
Daucus_carota_GWD          492 ADDLK--YQVQSIQIEVE-ENFAGMPFVLLSGANWIKNNGSDFYVNFSSGPKLIHKD---
Gossypium_hirsutum_GWD     504 SGDLP--KQVQCIEMEIADGNFKGMPFVLLSGGKWIKNNGSDFYVEFSQRFKQVQKD---
Ricinus_communis_GWD       501 SAELP--YQVQSFELEIEEDNFVGMPFVLLSNGNWIKNKGSDFYIEFSGGPKQVQKD---
Vitis_vinifera_GWD         502 SADPA--YEVQTLKIEIEEDSFVGMPFVLLSQGNWIKNGGSDFYIEFRVGPKQVKKD---
Camellia_sinensis_GWD      506 SDNPI--DKVQSLEIEIKDDSYVGMPFVLVSGGKWIKNDGSDFYVKFDTGYKQVQKD---
Zea_mays_GWD               499 LDGLH----YQVVEIELDDGGYKGMPFVLRSGETWIKNNGSDFFLDFSTHDVRNIKLKGN
Phoenix_dactylifera_GWD    508 SADLF----YQAIEIEIDGGDYNGMPFVLLSDGKWMKDNDSDFYIDFGSETIESWKD---
Zingiber_officinale_GWD    515 MIDQD----YQAIQIEIDGGDYAGIPFVLRSDDKWIKNSGLDFYIELGNGSIKSRKA---

                           601 .......610.......620.......630.......640.......650.......660
Nymphaea_colorata_GWD1     556 -IDGKGTAQSLLNRIAELESEAQRSLMHRFNIATDLMEWAKDEGELGLAGILVWMRFMAT
Beta_vulgaris_GWD          556 AGDGSGTAKLLLEKIASLESEAQKSFMHRFNIAADLLDEAIDAGELGLAGMLVWMRFMAT
Arabidopsis_thaliana_GWD   487 YGDGKGTAKHLLDKIADLESEAQKSFMHRFNIAADLVDEAKSAGQLGFAGILVWMRFMAT
Erigeron_canadensis_GWD    482 VGDGKGTAKALLDKIAGLESEAQKSFMHRFNIAADLMEEAKNTGELGLAAILVWMRFMAT
Amborella_trichopoda_GWD1  559 AGNGEGTAKALLDRISELESDAERSFMHRFNIATDLTEWAKDQGELGLAGLLVWMRFMAT
Fragaria_vesca_GWD         489 AGDGKGTAKALLDNIADLESEAQKSFMHRFNIAADLMSQAKDAGALGLAGILVWMRFMAT
Phaseolus_vulgaris_GWD     542 FGDSKGTAKFLLDKIAEQESEAQKSFMHRFNIASNLIDEAKSAGRLGLAGILVWMRFMAT
Erythranthe_guttata_GWD    556 AGDGKGTSKSLLDKIADLESEAQKSFMHRFNIAADLMEQATNAGELGLAAIVVWMRYMAT
Cucumis_sativus_GWD        558 TEGGKGTAKSLLDNIAELESEAEKSFMHRFNIAADLVDQAKDAGELGLAGILVWMRFMAT
Magnolia_sinica_GWD1       563 AGDGKGTAKALLDKIAQMESEAQKSFMHRFNIAADLMEWAKDAGELGLAGMLVWMRFMAT
Nelumbo_nucifera_GWD       552 VGDGKGTAKALLDKIAEMEGEAQKSFMHRFNIASDLTEWAKDAGELGLAGILVWMRFMAT
Ipomoea_triloba_GWD        555 VGDGMGTAKALLDKIADMESEAQKSFMHRFNIAADLTEEATNAGELGFAGILVWMRFMAT
Daucus_carota_GWD          546 VGDGKGTAKSLLDKIAGLESEAQKSFMHRFNIAADLVQEAKDAGELGLAGILVWMRFMAT
Gossypium_hirsutum_GWD     559 AGDGKGTSKVLLDRIAALESEAQKSFMHRFNIASDLMDQAKNIGELGLAGILVWMRFMAT
Ricinus_communis_GWD       556 AGNGRGTAKALLDKIAEMESEAQKSFMHRFNIAADLMEQAKDSGELGLAGILVWMRFMAT
Vitis_vinifera_GWD         557 AGDGKGTAKALLDKIAEKESEAQKSFMHRFNIAADLMDQAISAGKLGLAGIVVWMRFMAT
Camellia_sinensis_GWD      561 AGDGQGTAKTLLDKIAKMESEAQKSFMHRFNIAADLVDEAKYAGELGFAGILVWLRFMAT
Zea_mays_GWD               555 GDAGKGTAKALLERIADLEEDAQRSLMHRFNIAADLADQARDAGLLGIVGLFVWIRFMAT
Phoenix_dactylifera_GWD    561 ARDGRGTAKSLLDKIAELESDAQRSLMHRFNIAADLMEQARDAGQLGFAGILVWMRFMAT
Zingiber_officinale_GWD    568 PGDGSGVAKSLLDKIAELETEAQRSLMHRFNIAADLAEQARGSGQLGLAGILVWLRFMAM

                           661 .......670.......680.......690.......700.......710.......720
Nymphaea_colorata_GWD1     615 RQLTWNKNYNVKPREISRAQDSLTDSLQKIFQTYPQYREIVRMVMFTVGRGGQGDVGQRI
Beta_vulgaris_GWD          616 RQLIWNKNYNVKPREISKAQDRLTESLQNIFRSHPQYGELLRMIMSTIGRGGEGDVGQRI
Arabidopsis_thaliana_GWD   547 RQLVWNKNYNVKPREISKAQDRLTDLLQDVYASYPEYRELLRMIMSTVGRGGEGDVGQRI
Erigeron_canadensis_GWD    542 RQLIWNKNYNVKPREISRAQDRLTDLLQSVYVNYPQYSELLRMIMSTVGRGGEGDVGQRI
Amborella_trichopoda_GWD1  619 RQLTWNRNYNVKPREISKAQDNLTDSLQRIYESYPQYREIVRMIMSTVGRGGEGDVGQRI
Fragaria_vesca_GWD         549 RQLIWNKNYNVKPREISKAQDRLTDLLQSIYISYPQHRELLRMILSTVGRGGEGDVGQRI
Phaseolus_vulgaris_GWD     602 RQLIWNKNYNVKPREISKAQDRLTDLLQDVYASYPQYREIVRMILSTVGRGGEGDVGQRI
Erythranthe_guttata_GWD    616 RQLIWNKNYNVKPREISQAQDRLTDLLQNVYKSSPQYREILRMIMSTVGRGGEGDVGQRI
Cucumis_sativus_GWD        618 RQLIWNKNYNVKPREISKAQDRLTDLLENIYANHPQYREILRMIMSTVGRGGEGDVGQRI
Magnolia_sinica_GWD1       623 RQLIWNKNYNVKPREISKAQDRLTDLLQDVYKTCPQYREMVRMIMSTVGRGGEGDVGQRI
Nelumbo_nucifera_GWD       612 RQLIWNKNYNVKPREISKAQDRLTDLLQNIYKNKPQYREILRMILSTVGRGGEGDVGQRI
Ipomoea_triloba_GWD        615 RQLIWNKNYNVKPREISKAQDRLTDLLQNVYLSRPQYRELLRMILSTVGRGGEGDVGQRI
Daucus_carota_GWD          606 RQLIWNKNYNVKPREISKAQDRLTDLLQNVYKSHPEYRELLRMIMSTVGRGGEGDVGQRI
Gossypium_hirsutum_GWD     619 RQLIWNKNYNVKPREISKAQDRLTDLLQSIYTTHPQHRELLRMIMSTVGRGGEGDVGQRI
Ricinus_communis_GWD       616 RQLIWNKNYNVKPREISKAQDRLTDLLQNIYTSQPQYREILRMIMSTVGRGGEGDVGQRI
Vitis_vinifera_GWD         617 RQLVWNKNYNIKPREISKAQDRLTDLLQNSYKTHPQYRELLRMIMSTVGRGGEGDVGQRI
Camellia_sinensis_GWD      621 RQLIWNKNYNVKPREISRAQDRLTDLLQDVYKTHAQYRELLRMILSTVGRGGEGDVGQRI
Zea_mays_GWD               615 RQLTWNKNYNVKPREISKAQDRFTDDLENMYKAYPQYREILRMIMAAVGRGGEGDVGQRI
Phoenix_dactylifera_GWD    621 RQLIWNKNYNVKPREISKAQDRLTDLLQNMYKICPQYREILRMIMSTVGRGGEGDVGQRI
Zingiber_officinale_GWD    628 RHLIWNKNYNVKPREISKAQDRLTDLLQDIYKDSPQYREILRMIMATVGRGGEGDVGQRI

                           721 .......730.......740.......750.......760.......770.......780
Nymphaea_colorata_GWD1     675 RDEILVIQRNNDCKGGMMEEWHQKLHNNTSPDDVVICQALIDYIRNDFDVNVYWNTLISN
Beta_vulgaris_GWD          676 RDEILVIQRNNDCKGGFMEEWHQKLHNNTSPDDVVICQALLDYIKSDFDISVYWKTLNEN
Arabidopsis_thaliana_GWD   607 RDEILVIQRKNDCKGGIMEEWHQKLHNNTSPDDVVICQALMDYIKSDFDLSVYWKTLNDN
Erigeron_canadensis_GWD    602 RDEILVIQRNNDCAGGMMEEWHQKLHNNTSPDDVVICQALIDYIKNDFDISVYWNTLNTN
Amborella_trichopoda_GWD1  679 RDEILVIQRNNDCKGGMMEEWHQKLHNNTSPDDVVICQALIDYISSDFDISVYWNTLNSN
Fragaria_vesca_GWD         609 RDEILVIQRNCDCKGGMMEEWHQKLHNNTSPDDVVICQALIDYIKSDFDIGVYWKTLNDN
Phaseolus_vulgaris_GWD     662 RDEILVIQRNNDCKGGMMEEWHQKLHNNTSPDDVVICQALIDYIKNDFDTGVYWKTLNDN
Erythranthe_guttata_GWD    676 RDEILVIQRNNECKGGMMEEWHQKLHNNTSPDDVVICQALIDYIKNDFDIGVYWKTLNDN
Cucumis_sativus_GWD        678 RDEILVIQRNNDCKGGMMEEWHQKLHNNTSPDDVVICQALIDYINSDFDIGVYWKTLNEN
Magnolia_sinica_GWD1       683 RDEILVIQRNNDCKGGMMEEWHQKLHNNTSPDDVVICQALIDYIKSDFNIDVYWNTLNSN
Nelumbo_nucifera_GWD       672 RDEILVIQRNNDCKGGMMEEWHQKLHNNTSPDDVIICQALIDYIKSDFDISVYWKTLNSN
Ipomoea_triloba_GWD        675 RDEILVIQRKNDCKGGMMEEWHQKLHNNTSPDDVVICQALIDYIKSDFDISVYWQTLNSN
Daucus_carota_GWD          666 RDEILVIQRNNDCKGGMMEEWHQKLHNNTSPDDVVICQALLDYIKCDFDIDVYWRTLNDN
Gossypium_hirsutum_GWD     679 RDEILVIQRNNDCKGGMMEEWHQKLHNNTSPDDVIICQALIDYIKSDFDISVYWKTLNEN
Ricinus_communis_GWD       676 RDEILVIQRNNDCKGGMMEEWHQKLHNNTSPDDVVICQALIDYISSGFDISMYWKSLNEN
Vitis_vinifera_GWD         677 RDEILVLQRNNDCKGAMMEEWHQKLHNNTSPDDVIICQALIDYIKCDFDISAYWKTLNEN
Camellia_sinensis_GWD      681 RDEILVIQRKNECKGGMMEEWHQKLHNNTSPDDVVICQALIDYIKSDFDISVYWKTLNEN
Zea_mays_GWD               675 RDEILVIQRNNDCKGGMMEEWHQKLHNNTSPDDVVICQALIDYIKSDFDISVYWDTLNKN
Phoenix_dactylifera_GWD    681 RDEILVIQRNNNCKGGMMEEWHQKLHNNTSPDDVVICQALIDYIKSDFDIKVYWDTLSKS
Zingiber_officinale_GWD    688 RDEILVIQRNNDCKGGMMEEWHQKLHNNTSPDDVVICQALIDYIKSDFDISVYLDTLNKN

                           781 .......790.......800.......810.......820.......830.......840
Nymphaea_colorata_GWD1     735 GITKERLLSYDRAIHSEPQFRRDQKEGLLRDLGNYMKTLKAVHSGADLESAIATCMGYST
Beta_vulgaris_GWD          736 GITKERLLSYDRAIHSEPKLRSDQKEGLLRDLGHYLKTLKAVHSGADLEAAISNCMGYKT
Arabidopsis_thaliana_GWD   667 GITKERLLSYDRAIHSEPNFRGEQKDGLLRDLGHYMRTLKAVHSGADLESAIQNCMGYQD
Erigeron_canadensis_GWD    662 GITKERLLSYDRAIRNEPKFRGDQKESLLRDLGNYMRTLKAVHSGADLESAISNCMGYKS
Amborella_trichopoda_GWD1  739 GITKERLLSYDRGIHSEPHFRRDQKEGLLRDLGNYLRTLKAVHSGADLQSAIATCMGYSA
Fragaria_vesca_GWD         669 GITKERLLSYDRAIHNEPRFRADQKDRLLHDLGNYLRTLKAVHSGADLESAITNCLGYSA
Phaseolus_vulgaris_GWD     722 GITKERLLSYDRAIHSEPNFRRDQKEGLLRDLGNYMRTLKAVHSGADLESAISNCMGYKS
Erythranthe_guttata_GWD    736 GITKERLLSYDRAIHSEPNFRREQRDGLLRDLGHYMRTLKAVHSGADLESAIANCMGYKA
Cucumis_sativus_GWD        738 GITKERLLSYDRAIHSEPNFRGDQKDGLLRDLGNYMRTLKAVHSGADLESAIQNCFGYRS
Magnolia_sinica_GWD1       743 GITKERLLSYDRAIHNEPHFRRDQKEGLLRDLGHYMRTLKAVHSGADLESAIANCMGYRA
Nelumbo_nucifera_GWD       732 GITKERLLSYDRAIHSEPNLRRDQKDGLLRDLGNYMRTLKAVHSGADLESAIANCMGYRS
Ipomoea_triloba_GWD        735 GITKERLLSYDRAIHSEPNFRSDQREGLLRDLGNYMRTLKAVHSGADLESAIANCMGYKT
Daucus_carota_GWD          726 GITKERLLSYDRAIHSEPNFRRDQKDGLLRDLGSYMRTLKAVHSGADLESAISNCMGYKS
Gossypium_hirsutum_GWD     739 GITKERLLSYDRAIHSEPSFKRDQKDGLLRDLGHYMRTLKAVHSGADLESAISNCMGYRA
Ricinus_communis_GWD       736 GITKERLLSYDRAIHSEPNFRRDQKDGLLRDLGNYMRTLKAVHSGADLESAIANCMGYRA
Vitis_vinifera_GWD         737 GITKERLLSYDRGIHSEPNFRKDQKDGLLRDLGKYMRTLKAVHSGADLESAISNCMGYRS
Camellia_sinensis_GWD      741 GITKERLLSYDRAIHSEPNFRRDQKDGLLRDLGSYMRTLKAVHSGADLESAISNCMGYRS
Zea_mays_GWD               735 GITKERLLSYDRAIHSEPNFRSEQKAGLLRDLGNYMRSLKAVHSGADLESAIASCMGYKS
Phoenix_dactylifera_GWD    741 GITRERLLSYDRAIHSEPSFRSDQKEGLLRDLGNYMRTLKAVHSGADLESAIATCIGYKS
Zingiber_officinale_GWD    748 GITKERLLSYDRAIHSEPSFRRDQKEGLLRDLGNYMRTLKAVHSGADLESAIATCMGYKS

                           841 .......850.......860.......870.......880.......890.......900
Nymphaea_colorata_GWD1     795 EGQGFMVGVNIHPVQGLPTGFPELLQFVLDHVEDKNVEPLLEGLLEARAELRSLLLGSCD
Beta_vulgaris_GWD          796 EGQGFMVGVNIDPIPGLPAGFPNLLQFVLEHVEDKNVEPLVEGLLEARQELRPSLFKSQD
Arabidopsis_thaliana_GWD   727 DGEGFMVGVQINPVSGLPSGYPDLLRFVLEHVEEKNVEPLLEGLLEARQELRPLLLKSHD
Erigeron_canadensis_GWD    722 EGQGFMVGVNINPVSGLPSGFPELLQFILEHVEDRNVETLLEGLLEAREELRPLLSKPND
Amborella_trichopoda_GWD1  799 QGQGFMVGVEVHPISGLPSGFPELLQFILHHVEDKQVEPLLEGLLEARVELRPLLLRSHD
Fragaria_vesca_GWD         729 KGQGFMVGVQINPVSGLPSEVPGLLQFVMEHVEDRNVEVLVEGLLEARQEIWPLLSKPND
Phaseolus_vulgaris_GWD     782 EGQGFMVGVQINPVPGLPAGFQGLLEFVMEHVEDKNVEPLLEGLLEAREELHPSLGKSQS
Erythranthe_guttata_GWD    796 EGQGFMVGVNINPVSGLPSGFPELLQFVLTHIEDKQVESLLEGLLEAREELRPSLSRPSD
Cucumis_sativus_GWD        798 EGQGFMVGVQINPISGLPSELPGLLQFVLEHIEIKNVEPLLEGLLEARQELRPLLLKPRD
Magnolia_sinica_GWD1       803 EGQGFMVGVQINPISGIPSGYPELLQFVLDHVEDKRVEYLLEGLLEARAELRPLLLKSHD
Nelumbo_nucifera_GWD       792 EGQGFMVGVQINPVPGLPSGFPELLEFVLDHVEDTNVEPLLEGLLEARQELQPLLLKSYE
Ipomoea_triloba_GWD        795 EGEGFMVGVQINPVSGLPSGFPELLQFVLEHVEDKNVEALLEGLLEAREELRPLLFQSNN
Daucus_carota_GWD          786 EGQGFMVGVEINPVSGLPSGFPDLLQFILAHVEDKNVEPLLEGLLEVREELRPLLSNPND
Gossypium_hirsutum_GWD     799 EGQGFMVGVQINPIPGLPSGFPDLLRFVLEHIEDRNVEALLEGLLEARQELRPLLLKSTG
Ricinus_communis_GWD       796 EGQGFMVGVQINPISGLPSGFPELLQFVLEHVEDKNVEALLEGLLEARQELRPLLFKSHD
Vitis_vinifera_GWD         797 EGQGFMVGVKINPIPGLPSGFPELLQFVLEHVEDKNVEPLLEGLLEARQELQSLLIKSHD
Camellia_sinensis_GWD      801 EGQGFMVGVQINPISGLPSGFPELLQFVREHVEDKNVEALLEGLLEAREELRPLLLKSND
Zea_mays_GWD               795 EGEGFMVGVQINPVKGLPSGFPELLEFVLEHVEDKSAEPLLEGLLEARVELRPLLLDSRE
Phoenix_dactylifera_GWD    801 EGQGFMVGVQIDPIRSLPSGFCDLLKFILDHLEDKMAEPLLEGLLEARVELRQLLLNSHE
Zingiber_officinale_GWD    808 EGQGFMVGVQINPIRGLPSGISGLMEFILKHVEDKSVEPLIEGLLEARVELRPLLLSSHE

                           901 .......910.......920.......930.......940.......950.......960
Nymphaea_colorata_GWD1     855 RLKDLIFLDLALDSTVRTAVERGYEELDHAEPQKVMYFITLVLENLALSSNENEDIVYCL
Beta_vulgaris_GWD          856 RLKDLMFLDIALDSSVRTVIERGYEELNQAGPEKIMSFISMVVENLALSSDDNEDLLYCI
Arabidopsis_thaliana_GWD   787 RLKDLLFLDLALDSTVRTAIERGYEQLNDAGPEKIMYFISLVLENLALSSDDNEDLIYCL
Erigeron_canadensis_GWD    782 RLKDLLFLDIALDSTVRTAIERSYEDLSNAKPEKVMYLITLLLENLILSSDNNEDLIYCL
Amborella_trichopoda_GWD1  859 RLKDLIFLDLALDSTVRTAIERGYEELNNAEPQKIMHFIALVLENLVLSSDSNEDLIYCL
Fragaria_vesca_GWD         789 RLKDLLFLDIALDSTVRTAIERGYEELNNAGPEKIMYFISMVLENLALSSDDNEDLLYCL
Phaseolus_vulgaris_GWD     842 RLKDLLFLDVALDSTVRTAVERGYEELNNAAPEKIMYFICLVLENLSLSSDDNEDLIYCL
Erythranthe_guttata_GWD    856 RLKDLIFLDIALDSAVRTAVERGYEELNNASPEKIIYFISLVVENLALSVDNNEDLIYCL
Cucumis_sativus_GWD        858 RLRDLLFLDIALDSAVRTAVERGYEELNTAGPEKIMYFITLVLENLALSSDDNEDLIYCL
Magnolia_sinica_GWD1       863 RLRDLLFLDIALDSTVRTALERGYEELNNAEPEKIMYFISLVLENLALSSDNNEDLIYCL
Nelumbo_nucifera_GWD       852 RLRDLLFLDIALDSMVRTAIERGYEELNKAGPEKIMYFISMVLENLALSSDNNEDLINCL
Ipomoea_triloba_GWD        855 RLKDLLFLDIALDSTVRTAIERGYEELNNASPEKIMYFISLVLENLALSVDDNEDLVYCL
Daucus_carota_GWD          846 RLKDLLFLDIALDSTVRTAIERGYEELNGAKPEKVMYFITLLLENLALSSDNNEDLIFCL
Gossypium_hirsutum_GWD     859 RLKDLLFLDIALDSTVRTAIERGYEELNNARPEKIMHFITLVLENLALSSDDNEDLVYCL
Ricinus_communis_GWD       856 RLKDLLFLDIALDSTVRTVIERGYEELNNAGQEKIMYFITLVLENLALSSDDNEDLIYCM
Vitis_vinifera_GWD         857 RLKDLLFLDIALDSTVRTAIERGYEELNNAGAEKIMYFITLVLENLVLSSDDNEDLIYCL
Camellia_sinensis_GWD      861 RLKDLLFLDIALDSTVRTAVERGYEELNNAKPEKVMFFITLLLENLALSSDDNEDLIYCL
Zea_mays_GWD               855 RMKDLIFLDIALDSTFRTAIERSYEELNDAAPEKIMYFISLVLENLALSIDDNEDILYCL
Phoenix_dactylifera_GWD    861 RTKDLLFLDIALDSTVRTAIERAYEELNNAEPEKIMYLIILVLENLALSTDDNEDLIYCL
Zingiber_officinale_GWD    868 RLKDLIFLDIALDSTVRTVVERGYEELSNAEPEKLIYLITLLLENLALSTDDNEDLIYCL

                           961 .......970.......980.......990......1000......1010......1020
Nymphaea_colorata_GWD1     915 KGWDHAIQMLKSKSGCWALFAKSVLDRTRLALATKAEHFQNVLQPSAEYLGSMLGVEQWA
Beta_vulgaris_GWD          916 KGWNHALDMLKSKDGDWALYAKSVLDRTRLALASKGEFYQQVLQPSADYLGSLLGVDQWA
Arabidopsis_thaliana_GWD   847 KGWQFALDMCKSKKDHWALYAKSVLDRSRLALASKAERYLEILQPSAEYLGSCLGVDQSA
Erigeron_canadensis_GWD    842 KGWNQALSMLETRDGNWALFAKSVLDRTRLALASKGEFYHHILQPSAEYLGARLNLDQRA
Amborella_trichopoda_GWD1  919 KEWNYTLQMSKSQDDHWALYAKSVLDRSRLALTSKAEHYQRILQPSAEYLGSLLGVDKWA
Fragaria_vesca_GWD         849 KGWNQALNMLKNNADHWALFAKSILDRTRLALANKAELYHSILQPSAEYLGSKLGVDKLA
Phaseolus_vulgaris_GWD     902 KGWDLALTKCKSNDTHWALYAKSVLDRTRLALTNKAQLYQEILQPSAEYLGSLLGVDQWA
Erythranthe_guttata_GWD    916 KGWNQALSMKKSGDGNWALFAKSVLDRTRLSLTSKSESYNQLLQPSAEYLGAQLGVDQSA
Cucumis_sativus_GWD        918 KGWDLALNLTRSKNDHWALYAKSVLDRTRLALANKGEEYHRILQPSAEYLGSLLGVDQWA
Magnolia_sinica_GWD1       923 KGWNNALDLSRKRDDQWALYAKAVLDRTRLALASKGEYYHQVLQPSAEYLGSLLGVDQWA
Nelumbo_nucifera_GWD       912 KGWSHALDMSKSRDDHWALYAKSVLDRTRLALASKAEHYQQVLQPSAEYLGSLLGVDQWA
Ipomoea_triloba_GWD        915 KGWNQALKMSKSGDNNWALFAKSVLDRTRLSLANKAESYHHLLQPSAEYLGSKLGVDEWA
Daucus_carota_GWD          906 KGWNQAISMLSSGDDHWALYAKSVLDRTRLALTSKAEWYHRQLQPSAEYLGAQLGVDEWA
Gossypium_hirsutum_GWD     919 KGWHHSISMCKSKSAHWALYAKSVLERTRLALASKAETYQRILQPSAEYLGSLLGVDQWA
Ricinus_communis_GWD       916 KGWNHALSMSKSKSDQWALYAKSVLDRTRLALSSKAEWYQQVLQPSAEYLGSLLGVDQWA
Vitis_vinifera_GWD         917 KGWNHALGMSKSRDGHWALYAKSVLDRTRLALTSKAEEYHQVLQPSAEYLGSLLGVDQWA
Camellia_sinensis_GWD      921 KGWNQALSMIKNGDSHWALYAKSVLDRTRLALTSKAEWYQHLLQPSAEYLGSRLGVDQWA
Zea_mays_GWD               915 KGWNQALEMAKQKDDQWALYAKAFLDRNRLALASKGEQYHNMMQPSAEYLGSLLSIDQWA
Phoenix_dactylifera_GWD    921 KGWNHALEMSKQKDDQWALYAKSFLDRTRLALSSKAELYHQILQPSAEYLGSLLGVDQWA
Zingiber_officinale_GWD    928 KGWKHSMEMSKQKDDQWALFAKSFLDRTRLALSSKAEYYHQILQPSAEYLGSLLDVDPWA

                          1021 ......1030......1040......1050......1060......1070......1080
Nymphaea_colorata_GWD1     975 LSIFTEEVIRAGSAASLSMLLNRIDPVLRRLANLGSWQVISPVEVTGSVAVVDELLSVQN
Beta_vulgaris_GWD          976 VSIFTEEIVRAGSAASLSALLNRLDPVLRNTANLGSWQIISPVEAIGYVVVVKELLSVQN
Arabidopsis_thaliana_GWD   907 VSIFTEEIIRAGSAAALSSLVNRLDPVLRKTANLGSWQVISPVEVVGYVIVVDELLTVQN
Erigeron_canadensis_GWD    902 VSIFTEEMIRAGSAAPLSSLVNRLDPILRSVANLGSWQVISPIEVVGYVVVVDELLSVQN
Amborella_trichopoda_GWD1  979 VSIFTEEIIRAGSAASLSLLLNRLDPILRETAHLGSWQVISPVEVIGYVVIVNELLAVQN
Fragaria_vesca_GWD         909 LSIFTEEVIRAGSAASLSTLVNRLDPVLRKTANLGSWQVISPVEVVGYVVVVDELLTVQN
Phaseolus_vulgaris_GWD     962 VEIFTEEIIRAGSAASLSTLLNRLDPVLRKTANLGSWQVISPVETVGYVEVVDELLSVQN
Erythranthe_guttata_GWD    976 VSIFTEEIIRAGSAASLSSLLNRLDPVLRQTANLGSWQVISPIEAIGYVVVVDQLLSVQN
Cucumis_sativus_GWD        978 VDIFTEEIIRSGSASSLSSLLNRLDPVLRTTANLGSWQIISPVEAVGYVVVVDELLAVQN
Magnolia_sinica_GWD1       983 VNLFTEEVIRAGSAASLSSLLNRLDPVLRKTAHLGSWQVISPVEAVGYVVVVDELLAVQN
Nelumbo_nucifera_GWD       972 INIFTEEIIRAGSAASLSSLLNRLDPILRKTAHLGSWQIISPVETVGCVVVVDELLAVQN
Ipomoea_triloba_GWD        975 VNIFTEEIIRAGSAASLSSLLNRLDPILRQTAHLGSWQIISPVEAVGYIVVVDELLSVQN
Daucus_carota_GWD          966 VDIFTEEIIRAGSAASLSVLLNRLDPTLRETANLGSWQVISPVEAVGYVVVVDELLSVQN
Gossypium_hirsutum_GWD     979 INIFTEEIIRAGSAATLSSLINRLDPVLRETAHLGSWQVISPVEVVGYVEVVDELLSVQN
Ricinus_communis_GWD       976 VNIFTEEIIRAGSAASLSSLLNRLDPILRKTANLGSWQVISPVEVAGYVVVVDELLTVQN
Vitis_vinifera_GWD         977 VNIFTEEIIRAGSAASLSSLLNRLDPVLRKTANLGSWQVISPVEAVGRVVVVGELLTVQN
Camellia_sinensis_GWD      981 VNIFTEEIIRAGSAASLSSLINRLDPILRETANLGSWQVISPVEAIGYVVVVDELLTVQN
Zea_mays_GWD               975 VNIFTEEIIRGGSAATLSALLNRFDPVLRNVAHLGSWQVISPVEVSGYVVVVDELLAVQN
Phoenix_dactylifera_GWD    981 VSIFTEEVIRGGSAASLSALLNRLDPVLRKVAHLGSWQVISPVEVAGYVAVVNELLAVQN
Zingiber_officinale_GWD    988 VSIFTEEIIRAGSAASLSALLQRLDPLLRKVAHLGSWQVISPVEVAGYVEVVEELLAVQN

                          1081 ......1090......1100......1110......1120......1130......1140
Nymphaea_colorata_GWD1    1035 KSYEKPTILIAKRVKGEEEIPDGTVAVLTPDMPDVLSHVSVRARNSKVCFATCFDNNILA
Beta_vulgaris_GWD         1036 KIYAQPTILVANHVKGEEEIPDGTVAVLTPDMPDVLSHVSVRARNCKVCFATCFDSSIFS
Arabidopsis_thaliana_GWD   967 KTYDRPTIIVANRVRGEEEIPDGAVAVLTPDMPDVLSHVSVRARNGKICFATCFDSGILS
Erigeron_canadensis_GWD    962 KTYDMPTILVAKSVRGEEEIPDGAVAVVTPDMPDVLSHVSVRARNSKVCFATCFDTGILD
Amborella_trichopoda_GWD1 1039 VSYERPTVLVSKRVKGEEEIPDGTVAVLTPDMPDILSHVSVRARNSKVCFATCFDPNILS
Fragaria_vesca_GWD         969 KVYDKPTILVARSVRGEEEIPDGTVAVLTPDMPDVLSHVSVRARNSKVCFATCFDSNILA
Phaseolus_vulgaris_GWD    1022 KSYERPTILIAKSVKGEEEIPDGTVAVLTPDMPDVLSHVSVRARNSKVCFATCFDPNILA
Erythranthe_guttata_GWD   1036 NSYSKPTILVAKSVRGEEEIPDGVVAVLTPDMPDVLSHVSVRARNSKVCFATCFDPNILA
Cucumis_sativus_GWD       1038 KSYEKPTILVANRVKGEEEIPDGTVAVLTPDMPDVLSHVSVRARNGKVCFATCFDSSILS
Magnolia_sinica_GWD1      1043 ESYERPTVLVAKSVTGEEEIPDGTVAVLTPDMPDVLSHVSVRARNSKVCFATCFDPNILA
Nelumbo_nucifera_GWD      1032 KSYGQPTILVAKRVKGEEEIPDGTVAVLTPDMPDVLSHVSVRARNSKVCFATCFDTNVLS
Ipomoea_triloba_GWD       1035 KTYDKPTILVANTVKGEEEIPDGTVAVLTPDMPDVLSHVAVRARNSKVCFATCYDPSVLA
Daucus_carota_GWD         1026 KTYEKPTILVAKSVRGEEEIPDGTVAVLTPDMPDVLSHVSVRARNSKVCFATCFDPNILS
Gossypium_hirsutum_GWD    1039 KSYDRPTILVAKSVKGEEEIPDGTIAVLTPDMPDVLSHVSVRARNCKVCFATCFDPNILA
Ricinus_communis_GWD      1036 KSYGRPTILVARRVKGEEEIPDGTVAVLTPDMPDVLSHVSVRARNGKVCFATCFDHNILE
Vitis_vinifera_GWD        1037 KSYGQPTILVVKTVKGEEEIPDGAVAVLTPDMPDVLSHVSVRARNGKVCFATCFDPKILA
Camellia_sinensis_GWD     1041 KSYGRPTILVAKSVKGEEDIPDGAVAVLTPDMPDVLSHVSVRARNSKVCFATCFDPNILA
Zea_mays_GWD              1035 KSYDKPTILVAKSVKGEEEIPDGVVGVITPDMPDVLSHVSVRARNSKVLFATCFDHTTLS
Phoenix_dactylifera_GWD   1041 KSYGRPTILVARSVKGEEELPDGAVAVLTPDMPDVLSHVSVRARNSKVCFATCFDSNILT
Zingiber_officinale_GWD   1048 KSYTQSTILVAKHVRGEEEIPDGTVAVLTPDMPDVLSHVSVRARNSKVCFATCFDDNILD

                          1141 ......1150......1160......1170......1180......1190......1200
Nymphaea_colorata_GWD1    1095 DLQSKEGKILSLKPTSSDLVYRMIDAGEVSSNNE-ITSKEGSAPSLTISRKQFAGKYAIC
Beta_vulgaris_GWD         1096 DLQAAEGKLLRLQPTSTDVLYSEVNESELKGASSSEISDN--IPSISLSKKIFGGKYAVA
Arabidopsis_thaliana_GWD  1027 DLQGKDGKLLSLQPTSADVVYKEVNDSELSSPSSDNLEDAP--PSISLVKKQFAGRYAIS
Erigeron_canadensis_GWD   1022 DLRAKEGKLLQLKPTSSDITYSEVQEGDLKR--SNKLEEVGPSPNIKLVKKEFAGKFAIS
Amborella_trichopoda_GWD1 1099 DLQSKEGKLIRVKPTSSDLIYSEVKETETLNGSPLTAKVEESSPAITIARKEFAGRYAIS
Fragaria_vesca_GWD        1029 DLQACEGKLLRVKPTSADVVYSEVNESELGDASSTNLNEDT--PALTLVKKQFTGRYAIS
Phaseolus_vulgaris_GWD    1082 NLQESRGKLLRLKPTSADVVYSQVEEGEFIDDKSSHLKDVGSVSPISLVRKKFSGRYAVS
Erythranthe_guttata_GWD   1096 SIQASEGKLLCLKPTSADVVYSEMTDDELLSS-TNSKDDVSSAPSLTLVKKKFSGRYAIS
Cucumis_sativus_GWD       1098 DLQVKEGKLIRLKPTSADIVYSEVKEDEVQDASSIHENDAAPSP-VTLVRKHFSGKYAIV
Magnolia_sinica_GWD1      1103 DLQSKQGKLLSLKATSAEILYSEIEESELSDASSTNLKEDGSSPSLTLVRKQFAGRYAIP
Nelumbo_nucifera_GWD      1092 DLQAKAGKLLRLRPTSTDIIYSEAKDNELLKT-SSNLKEDESLPSISLVRKKFCGRYAIS
Ipomoea_triloba_GWD       1095 ELQAKGGQFLRLKPTSADIIYSEEKEVEVQ--SSANLMEVGPSQSLTLVKKHFAGRYAIT
Daucus_carota_GWD         1086 DLQSKEGKILHLRPTSADIVYSEAKDVDITG--SNNSEEVGPSGSITLTKKQFGGRYAIS
Gossypium_hirsutum_GWD    1099 DLQAKKGKLLRLKPSSADVVYSEVKEGELVDSSSSNLKGDG--PSVTLVRKQFVGKYAIS
Ricinus_communis_GWD      1096 KLQAHEGKLLQLKPTSADIVYNEISEGELADSSSTNMKEVGSSP-IKLVKKQFSGRYAIS
Vitis_vinifera_GWD        1097 DLQANEGKLLHLKPTSADIVYSAVKEGELTDSISTKSKDNDSLPSVSLVRKQFGGRYAIS
Camellia_sinensis_GWD     1101 DLQANEGKLLQIKPTSVDIVYSKVEGG-LTSASSTNLNEEGPLPSIKLVRKQFGGRYAIS
Zea_mays_GWD              1095 ELEGYDQKLFSFKPTSADITYREITESELQQSSSPNAEVGHAVPSISLAKKKFLGKYAIS
Phoenix_dactylifera_GWD   1101 ELQGNEGKLFRLKPTSSDILYSEIEEIEIS--SSAGIGDDQSPPFVSLVRKQFSGRYAIS
Zingiber_officinale_GWD   1108 EFRRNSGKLFRLKPTSDDIVYSEIGKTEPEDVGPVQAGDEQAPPSVTLVGKHFSGKYAIS

                          1201 ......1210......1220......1230......1240......1250......1260
Nymphaea_colorata_GWD1    1154 AHEFSNDLVGAKSRNIGFLKDKVPSWVGIPTSVALPFGVFEKVISYDANKEIADKVSSLK
Beta_vulgaris_GWD         1154 AEEFTGDMVGAKSRNISYLRGRVPSWVGIPPSVALPFGAFEEVLANGPNKEVSDKLKILK
Arabidopsis_thaliana_GWD  1085 SEEFTSDLVGAKSRNIGYLKGKVPSWVGIPTSVALPFGVFEKVISEKANQAVNDKLLVLK
Erigeron_canadensis_GWD   1080 SEEFTSELVGAKSRNIGYLKGKVPSWVGIPTSVALPFGSFEKVLSDELNQGVFEKLQLLK
Amborella_trichopoda_GWD1 1159 SDEFSPEMVGAKSRNISYLKGKVPSWVGLPTSVALPFGVFEKVLSEDSNKNVAEKIEVLK
Fragaria_vesca_GWD        1087 SDEFTSEMVGAKSRNISYIKGKLPSWVGIPTSVALPFGVFEKVLSEDSNKAVADKIETLK
Phaseolus_vulgaris_GWD    1142 SEEFTGEMVGAKSRNITYLKGKVASWIGIPTSVAIPFGVFEHVLSDKSNQAVAERVNILK
Erythranthe_guttata_GWD   1155 SEEFTNDMVGAKSRNIANLKGKLPSWVNIPTSVALPFGVFETVLSDDLNKAVASKLDILK
Cucumis_sativus_GWD       1157 SEEFTSDLVGAKSRNISYLKGKVPSWVGIPTSVALPFGVFEEVLSDESNKAVAEKVHDLK
Magnolia_sinica_GWD1      1163 AHEFTSEMVGAKSRNISYLKGKVPSWVGIPTSVALPFGVFEKVLSNDSNQEVADKLDALK
Nelumbo_nucifera_GWD      1151 SEEFSSEMVGAKSRNIAYLKGKVPPWVGIPTSIALPFGVFEKVLTDDSNKVVADTLQTLK
Ipomoea_triloba_GWD       1153 SEEFTSELVGAKSRNIANLKGKVPSWIGIPTSVALPFGVFEKVLSDDINQGVDAKLQVLK
Daucus_carota_GWD         1144 SEEFTSELVGAKSRNIGYLKGKVPSWVGIPTSVALPFGVFEKVLSDDINKEVAEKVKVLQ
Gossypium_hirsutum_GWD    1157 AEEFTPEMVGAKSRNISYLKGKVPSWVGIPTSVALPFGVFEKVLADEANKEVDQKLQILK
Ricinus_communis_GWD      1155 SDEFTSEMVGAKSRNISHLKGKVPSWIGIPTSVALPFGVFEKVLSDGSNKEVAKKLELLK
Vitis_vinifera_GWD        1157 SEEFTSEMVGAKSRNISYLKGKVPLWVQIPTSVALPFGVFEKVLSDGLNKEVSEKLRSLK
Camellia_sinensis_GWD     1160 SEEFTSEVVGAKARNIGYLKGKVPSWVGIPTSVALPFGVFEKVLSDGLNQKVAEKLQLLK
Zea_mays_GWD              1155 AEEFSEEMVGAKSRNIAYLKGKVPSWVGVPTSVAIPFGTFEKVLSDGLNKEVAQSIEKLK
Phoenix_dactylifera_GWD   1159 AEEFTSEMVGAKSRNISFLKGKVPSWIGIPTSVALPFGVFEKVLSDNKNQAVADNLQMLK
Zingiber_officinale_GWD   1168 AEEFTNEMVGAKSRNISYLKGKVPSWVGIPTSVALPFGVFEEVLSNDINKEIVSQLQLLK

                          1261 ......1270......1280......1290......1300......1310......1320
Nymphaea_colorata_GWD1    1214 KKLEEGEFNVLHEIRDTVLQLVAPNELVEELKLKLRESGMPWPGDEGAERWEQAWMAIKR
Beta_vulgaris_GWD         1214 KELEGGDIGALGKIRTTVLGLVEPPQLVEELKSKMKSSGMPWPGDEGEKRWQQAWTSIKK
Arabidopsis_thaliana_GWD  1145 KTLDEGDQGALKEIRQTLLGLVAPPELVEELKSTMKSSDMPWPGDEGEQRWEQAWAAIKK
Erigeron_canadensis_GWD   1140 KKLVAGDFDVLEEIRKTVLDLVAPSQLVQELKTKMQSSDMPWPGDGGQQRWEQAWVAIKK
Amborella_trichopoda_GWD1 1219 KRLQGGEFSALHDIRETVLQLTASPQLVQELKDKMKSAGMPWPGDEGEQRWQQAWMAIKK
Fragaria_vesca_GWD        1147 KKLKEGDFGSLGEIRETVLQLTAPPPLVQELKSKMQSSGMPWPGDEGEQRWEQAWLSIKK
Phaseolus_vulgaris_GWD    1202 KKLIEGDFSVLKEIRETVLQLNAPPQLVEELKSKMKSSGMPWPGDEGEQRWEQAWKAIKK
Erythranthe_guttata_GWD   1215 RDLDEGNVGALGEIRNTVLELSAPPQLIKELKEKMQKSGMPWPGDEGAQRWEQAWIAIKK
Cucumis_sativus_GWD       1217 IKLGSGESSALKEIRKTVLQLAAPPQLVLELKSKMKSSGMPWPGDEGEKRWEQAWMAIKK
Magnolia_sinica_GWD1      1223 KRLGEGDFAVLGEIRQTVLRLSAPPQLVQELKDTMKSSGMPWPGDEGEQRWEQAWMAIKK
Nelumbo_nucifera_GWD      1211 KRLG-GDFSILGEIRKTVLQLSAPPQLVQELKNKMKSSGMPWPGDEGEQRWEQAWVAIKK
Ipomoea_triloba_GWD       1213 KKLSEGDFSVLGEIRNTVLELSAPPQLINELKDKMQSSGMPWPGDEGPERWEQAWMAIKK
Daucus_carota_GWD         1204 NKLEEEELSVLQEIRQTVLALQAPPQLVQELKSKMQSSGMPWPGDEGDQRWDQAWMAIKK
Gossypium_hirsutum_GWD    1217 KKLGEGDFGALEEIRQTVLQLRAPSQLVQELKTKMLTSGMPWPGDEGEQRWEQAWTAIKK
Ricinus_communis_GWD      1215 KKLGEGDFSVLGKIRETVLGLAAPQQLVQELKTSMQSSGMPWPGDEGEQRWQQAWMAIKK
Vitis_vinifera_GWD        1217 GGLGKGNFAVLTEIRKTVLQLSAPSQLVQELKDKMKSSGMPWPGDEGEQRWEQAWMAIKK
Camellia_sinensis_GWD     1220 KKLGEGDVSALKELRKTVLELSAPSQLVQELKDKMKSSGMPWPGDEGQQRWEQAWTAIKK
Zea_mays_GWD              1215 IRLAQEDFSALGEIRKVVLNLTAPMQLVNELKERMLGSGMPWPGDEGDKRWEQAWMAIKK
Phoenix_dactylifera_GWD   1219 ERLGQGEFGALDEIRRVVLQLAAPPQLVQELKEKMQGARMPWPGDEGVHRWEQAWMAVKK
Zingiber_officinale_GWD   1228 EKLAIGEFDALLNIRKMILQLASPIELVQELKGKMQASGMPWPGDEGEHRWELAWMAIKR

                          1321 ......1330......1340......1350......1360......1370......1380
Nymphaea_colorata_GWD1    1274 VWASKWNERAYFSTRKVKLDHDSLCMAVLVQEIISADYAFVIHTTNPSSSDPSEIYAEVV
Beta_vulgaris_GWD         1274 VWASKWNERAYFSTRKVGLDHEHLCMAVLVQEIISADYAFVIHTTNPSSGDSSEIYTEVV
Arabidopsis_thaliana_GWD  1205 VWASKWNERAYFSTRKVKLDHDYLCMAVLVQEVINADYAFVIHTTNPSSGDSSEIYAEVV
Erigeron_canadensis_GWD   1200 VWASKWNERAYFSTRKVGLDHDLLCMAVLVQEIINADYAFVIHTTNPSSGDSSEIYAEVV
Amborella_trichopoda_GWD1 1279 VWASKWNERAYFSTRKAKLDHNYLCMAVLVQEIISADYAFVIHTINPSSRDSSEIYAEVV
Fragaria_vesca_GWD        1207 VWASKWNERAYFSTRKVKLDHDYLCMAVLVQEIINADYAFVIHTTNPSSGDSSEIYAEVV
Phaseolus_vulgaris_GWD    1262 VWGSKWNERAYFSTRKVKLDHEYLSMAVLVQEVVNADYAFVIHTTNPSSGDSSEIYAEVV
Erythranthe_guttata_GWD   1275 VWASKWNERAYFSTRKVKLDHDYLCMAVLVQEIINADYAFVIHTTNPSSEDSSEIYAEVV
Cucumis_sativus_GWD       1277 VWASKWNERAYFSTRKVKLDHDYLCMAVLVQEIINADYAFVIHTTNPSSGDSSEIYAEVV
Magnolia_sinica_GWD1      1283 VWGSKWNERAYFSTRKVKLDHDYLCMAVLVQEIINADYAFVIHTTNPSSGDSSEIYAEVV
Nelumbo_nucifera_GWD      1270 VWASKWNERAYFSTRKVKLDHDYLCMAVLVQEIINADYAFVIHTTNPSSGDSSEIYAEVV
Ipomoea_triloba_GWD       1273 VWASKWNERAYFSTRKVKLDHDYLCMAVLVQEIINADYAFVIHTTNPSSGDLSEIYAEVV
Daucus_carota_GWD         1264 VWASKWNERAYFSTKKVKLDHNFLCMAVLVQEIINADYAFVIHTTNPSSGDPSEIYTEVV
Gossypium_hirsutum_GWD    1277 VWASKWNERAYFSTRKVKLDHDYLCMAVLVQEVINADYAFVIHTTNPSSGDTSEIYAEVV
Ricinus_communis_GWD      1275 VWASKWNERAYFSTRKVKLDHDYLCMAVLVQEIINADYAFVIHTTNPSSGDSSEIYAEVV
Vitis_vinifera_GWD        1277 VWASKWNERAYFSTRKVKLDHDYLCMAVLVQEIINADYAFVIHTTNPSSGDSSEIYAEVV
Camellia_sinensis_GWD     1280 VWASKWNERAYFSTRKVKLDHDYLCMAVLVQEIINADYAFVIHTTNPSSGDSSEIYAEVV
Zea_mays_GWD              1275 VWASKWNERAYFSTRKVKLDHEYLSMAVLVQEVVNADYAFVIHTTNPSSGDSSEIYAEVV
Phoenix_dactylifera_GWD   1279 VWASKWNERAYFSTRKVKLDHDFLCMAVLVQEIISADYAFVIHTTNPSSGDSSEIYAEVV
Zingiber_officinale_GWD   1288 VWASKWNERAYFSTRKVKLDHDYLCMAVLVQEIISADYAFVIHTTNPSSGDSSEIYAEVV

                          1381 ......1390......1400......1410......1420......1430......1440
Nymphaea_colorata_GWD1    1334 RGLGETLVGAYPGRALSFICKKNDLNSPKILGYPSKPFGLFIRQSIIFRSDSNGEDLEGY
Beta_vulgaris_GWD         1334 RGLGETLVGAYPGRALAFICKKSDLESPKVLGYPSKPIGLFIKRSIIFRSDSNGEDLEGY
Arabidopsis_thaliana_GWD  1265 KGLGETLVGAYPGRSLSFICKKNNLDSPLVLGYPSKPIGLFIRRSIIFRSDSNGEDLEGY
Erigeron_canadensis_GWD   1260 KGLGETLVGAYPGRALSFISKKDNLNSPKVLGYPSKPIGLFIRRSIIFRSDSNGEDLEGY
Amborella_trichopoda_GWD1 1339 KGLGETLVGAYPGRALSYVCKKTNLDSPKILGYPSKPIGLFIKRSIIFRSDSNGEDLEGY
Fragaria_vesca_GWD        1267 KGLGETLVGAYPGRALSFISKKNDLDSPQLLGYPSKPIGLFIRRSIIFRSDSNGEDLEGY
Phaseolus_vulgaris_GWD    1322 KGLGETLVGAYPGRALSFICKKSDLNSPQVLGYPSKPVGLFIRQSIIFRSDSNGEDLEGY
Erythranthe_guttata_GWD   1335 KGLGETLVGAYPGRALSFICKKSDLNSPQVLGYPSKPIGLFIRQSIIFRSDSNGEDLEGY
Cucumis_sativus_GWD       1337 KGLGETLVGAYPGRALSFICKKNDLDTPKVLGYPSKPIGLFIRRSIIFRSDSNGEDLEGY
Magnolia_sinica_GWD1      1343 KGLGETLVGAYPGRALSFVSKKDNLDSPKVLGYPSKPIGLFIRRSIIFRSDSNGEDLEGY
Nelumbo_nucifera_GWD      1330 KGLGETLVGAYPGRALSFVCKKNDLNSPKVLGYPSKPIGLFIRQSIIFRSDSNGEDLEGY
Ipomoea_triloba_GWD       1333 RGLGETLVGAYPGRALSFVCKKNDLNSPQVLGYPSKPIGLFIRRSIIFRSDSNGEDLEGY
Daucus_carota_GWD         1324 KGLGETLVGAYPGRALSFVCKKDDLNSPKVLGYPSKPIGLFIRRSIIFRSDSNGEDLEGY
Gossypium_hirsutum_GWD    1337 KGLGETLVGAYPGRALSFVCKKNNLNSPEVLGYPSKPIGLFIRRSIIFRSDSNGEDLEGY
Ricinus_communis_GWD      1335 RGLGETLVGAYPGRALSFVCKKQDLNSPQVLGYPSKPIGLFIRRSIIFRSDSNGEDLEGY
Vitis_vinifera_GWD        1337 RGLGETLVGAYPGRALSFICKKNDLNSPQVLGYPSKPIGLFITRSIIFRSDSNGEDLEGY
Camellia_sinensis_GWD     1340 KGLGETLVGAYPGRALSFVCKKNNLNSPQVLGYPSKPIGLFIRRSIIFRSDSNGEDLEGY
Zea_mays_GWD              1335 KGLGETLVGAYPGRAMSFVCKKDDLDSPKLLGYPSKPIGLFIRQSIIFRSDSNGEDLEGY
Phoenix_dactylifera_GWD   1339 KGLGETLVGAYPGRALSFVCKKNDLNSPKVLNFPSKPIGLFIRRSIIFRSDSNGEDLEGY
Zingiber_officinale_GWD   1348 KGLGETLVGAYPGRALSFVCNKNDLNSPKVLGFPSKPIGLFIKQSIIFRSDSNGEDLEGY

                          1441 ......1450......1460......1470......1480......1490......1500
Nymphaea_colorata_GWD1    1394 AGAGLYDSVPMDKEEKVFLDYTLDPLITDTNFQNSMLSSIARAGNAIEELYGTPQDIEGV
Beta_vulgaris_GWD         1394 AGAGLYDSVPMDEEEKVVLDYSADPLINDSNFRQSVLSSIAKAGSAIEELYGSPQDIEGV
Arabidopsis_thaliana_GWD  1325 AGAGLYDSVPMDEEDQVVLDYTTDPLITDLSFQKKVLSDIARAGDAIEKLYGTAQDIEGV
Erigeron_canadensis_GWD   1320 AGAGLYDSVPMDEEEKVVLDYSSDPLMVDVNFQKSILSSIAQAGDAIEKLYGSPQDIEGV
Amborella_trichopoda_GWD1 1399 AGAGLYDSVPMDEEEKVVLDYSTDRLLVDPGFRNSILSSIAKAGSAIEELYGSPQDIEGV
Fragaria_vesca_GWD        1327 AGAGLYDSVPMDKEEEVVLDYSSDPLVTDGNFQKKILSSIAHAGNAIEELYGLPQDIEGV
Phaseolus_vulgaris_GWD    1382 AGAGLYDSVPMDEEEKVVLDYSSDQLMLDGSFRRTILSSIARAGNEIEGLYGSPQDIEGV
Erythranthe_guttata_GWD   1395 AGAGLYDSVPMDEEDKVVLDYSSDALINDSKFRHEILSSIARAGSAIEELYGSAQDIEGV
Cucumis_sativus_GWD       1397 AGAGLYDSVPMDEEEKVVLDYTTDPLIVDDNFRKSILSSIARAGNAIEELYGSPQDIEGV
Magnolia_sinica_GWD1      1403 AGAGLYDSVPMDEEKKVVLDYLSDQLIIDGNFRKSILSSIARAGSAIEELYGSPQDIEGV
Nelumbo_nucifera_GWD      1390 AGAGLYDSVPMDEEEKVVLDYSSDRLITDGSFRHSILSSIARAGSAIEELYGSPQDIEGV
Ipomoea_triloba_GWD       1393 AGAGLYDSVPMDEEEKVVIDYSADPLITSNDFRHSMLSNIARAGSIIEELYGSPQDIEGV
Daucus_carota_GWD         1384 AGAGLYDSVPMDEEDKVVVDYSSDPLLIDSNFQKSILSSIARAGSAIEELYGSPQDIEGV
Gossypium_hirsutum_GWD    1397 AGAGLYDSVPMDKEEKVVVDYSSDPLINDGKFQQAILSSIAGAGNAIEELYGSPQDIEGV
Ricinus_communis_GWD      1395 AGAGLYDSVPMDEEEKVVIDYSSDPLIMDGNFRQSILSSIARAGSAIEELHGSAQDIEGV
Vitis_vinifera_GWD        1397 AGAGLYDSVPMDKEEKVVLDYSSDPLMIDGNFRQSILSSIARAGNAIEELYGSPQDIEGV
Camellia_sinensis_GWD     1400 AGAGLYDSVPMDEEEKIVLDYSSDPLIIDANFRQSILSSIARAGNAIEELYGSPQDIEGV
Zea_mays_GWD              1395 AGAGLYDSVPMDEEDEVVLDYTTDPLIVDRGFRSSILSSIARAGHAIEELYGSPQDVEGV
Phoenix_dactylifera_GWD   1399 AGAGLYDSVLMDEEEKVVLDYTSDPLVVDRNFRLSILSSIAKAGAAVEELYGTPQDIEGV
Zingiber_officinale_GWD   1408 AGAGLYDSVPMDEEEKVVLDYVADPLIMDKNFRNSLLSSIARAGYAIEELYGSPQDIEGV

                          1501 ......1510.....
Nymphaea_colorata_GWD1    1454 VKDGKIFVVQTRPQM
Beta_vulgaris_GWD         1454 VKDGKIFVVQTRPQM
Arabidopsis_thaliana_GWD  1385 IRDGKLYVVQTRPQV
Erigeron_canadensis_GWD   1380 VRDGKIYVVQTRPQM
Amborella_trichopoda_GWD1 1459 VKDGKIFVVQTRPQV
Fragaria_vesca_GWD        1387 IRDGKLYVVQTRPQM
Phaseolus_vulgaris_GWD    1442 IKDGKLYVVQTRPQM
Erythranthe_guttata_GWD   1455 VKDGKIYVVQTRPQM
Cucumis_sativus_GWD       1457 IRDGEVYVVQTRPQM
Magnolia_sinica_GWD1      1463 VKDEKIFVVQTRPQM
Nelumbo_nucifera_GWD      1450 VRDGKIFVVQTRPQM
Ipomoea_triloba_GWD       1453 VRDGKIFVVQTRPQM
Daucus_carota_GWD         1444 VKDGKIYVVQTRPQM
Gossypium_hirsutum_GWD    1457 IRDGKVYVVQTRPQM
Ricinus_communis_GWD      1455 IRDGKLYVVQTRPQM
Vitis_vinifera_GWD        1457 VRDGKIYVVQTRPQM
Camellia_sinensis_GWD     1460 VKDRKIYVVQTRPQM
Zea_mays_GWD              1455 VKDGKIYVVQTRPQM
Phoenix_dactylifera_GWD   1459 VKDGGIFVVQTRPQM
Zingiber_officinale_GWD   1468 VKDGKIFVVQTRPQM


```
