## Supplemental Table 1 and Figures 3-7 for "Functional Characterization of Arabidopsis α-Amylase3 (AMY3): Amylose Specificity and Structural Insights into Its Duplex Carbohydrate-Binding Module"

**Supplemental Table 1 NCBI accession numbers for AMY3 and GWD sequences from 20 orders of Angiosperms.**

**Order	Species	AMY3	GWD1**

Amborellales	*Amborella trichopoda*	XP_011622500.1	XP_006841018.2

Apiales	*Daucus carota*	XP_017222730.1	XP_017226385.1

Arecales	*Phoenix dactylifera*	XP_008808375.2	XP_038990345.1

Asterales	*Erigeron canadensis*	XP_043638843.1	XP_043613595.1

Brassicales	*Arabidopsis thaliana*	NP_564977.1	NP_001318975.1

Caryophyllales	*Beta vulgaris*	XP_010683251.1	XP_010695868.2

Cucurbitales	*Cucumis sativus*	XP_004135194.1	XP_031741257.1

Ericales	*Camellia sinensis*	XP_028099207.1	XP_028085458.1

Fabales	*Phaseolus vulgaris*	XP_068462925.1	XP_068462870.1

Lamiales	*Erythranthe guttata*	XP_012855272.1	XP_012848067.1

Magnoliales	*Magnolia sinica*	XP_058105304.1	XP_058088674.1

Malpighiales	*Ricinus communis*	XP_002520134.1	XP_002527902.1

Malvales	*Gossypium hirsutum*	XP_016705150.2	XP_016707329.2

Nymphaeales	*Nymphaea colorata*	XP_031504981.1	XP_031501423.1

Poales	*Zea mays*	XP_008675104.1	NP_001348353.1

Proteales	*Nelumbo nucifera*	XP_010256483.1	XP_010248572.1

Rosales	*Fragaria v*esca	XP_004297334.1	XP_004298987.1

Solanales	*Ipomoea triloba*	XP_031124848.1	XP_031124522.1

Vitales	*Vitis vinifera*	XP_002270049.1	XP_010651715.1

Zingiberales	*Zingiber officinale*	XP_042390883.1	XP_042408613.1

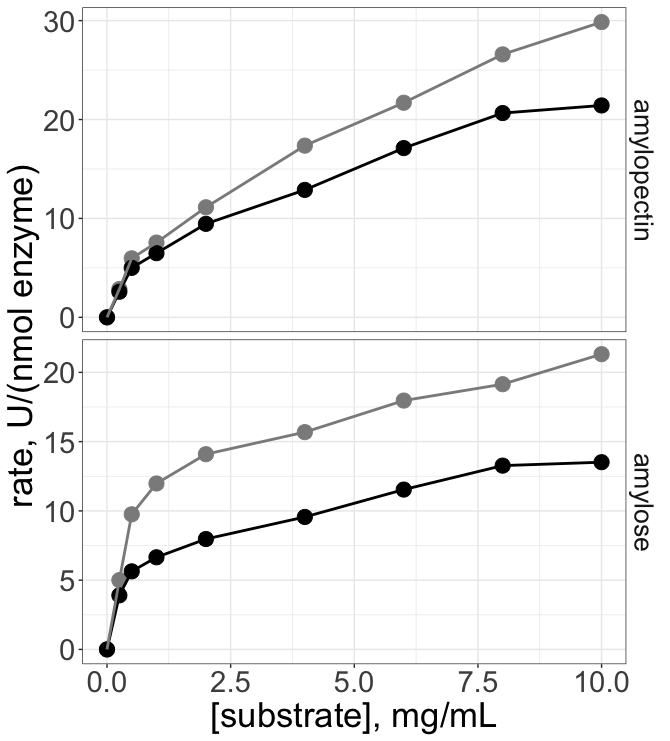

**Supplement Figure 3 Effect of BAM9 on AMY3 preference for amylose**

Kinetics of AMY3 (black) and AMY3+BAM9 (grey) assayed with various concentrations of solubilized amylose or amylopectin. Rates are expressed as μmoles of reducing equivalents produced per minute per nmol enzyme and are representative of 2 independent experiments for each enzyme.

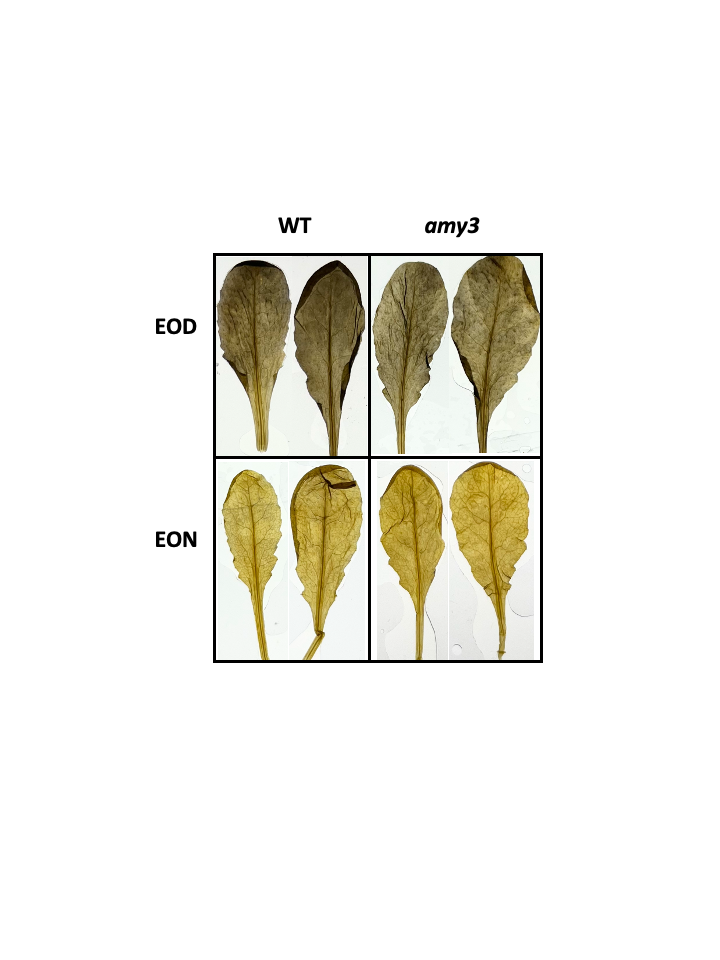

**Supplemental Figure 4** Arabidopsis leaves (WT and *amy3*) stained with Lugol’s iodine solution at the end of the day (EOD) or end of the night (EON).

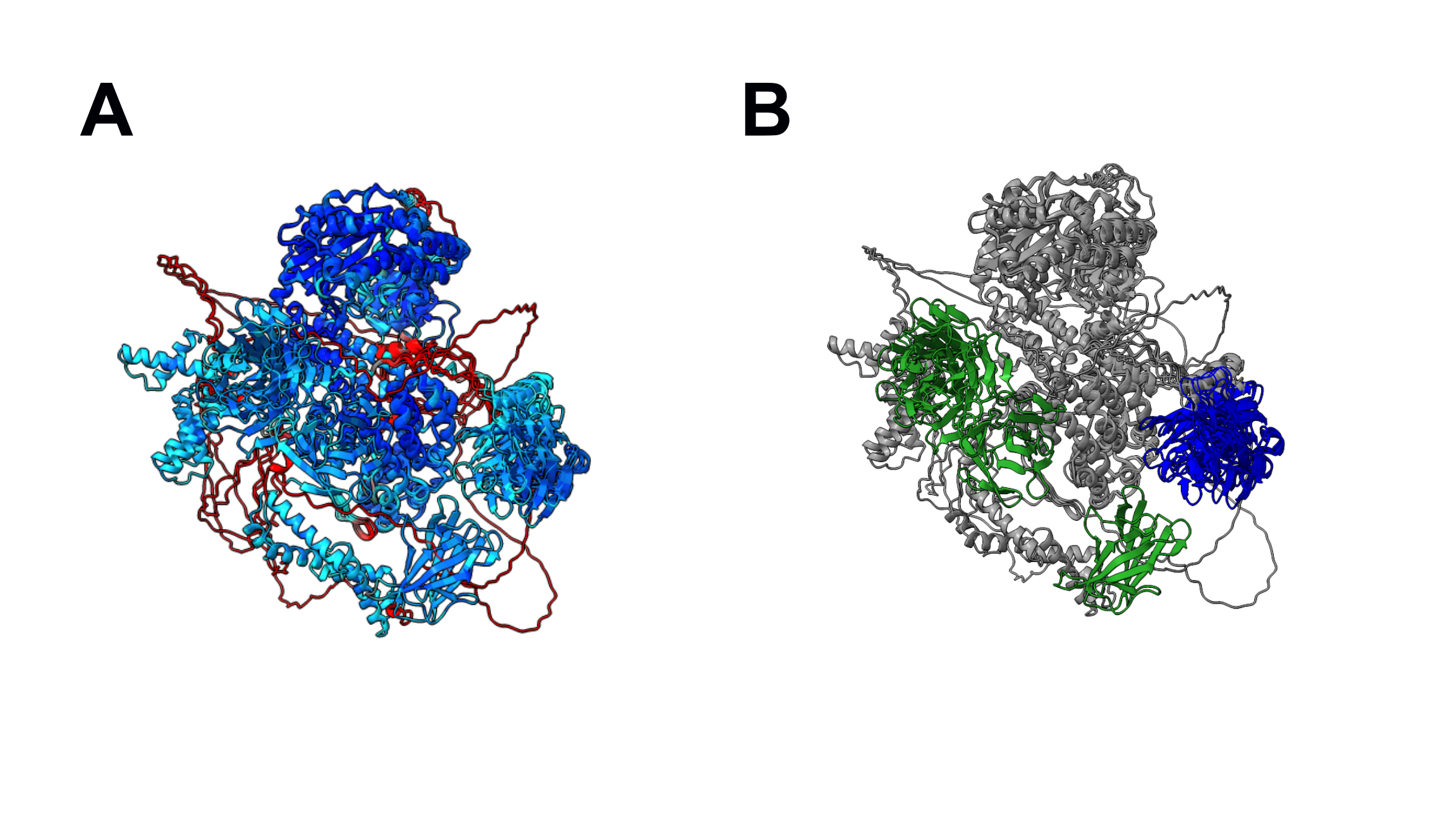
 **Supplemental Figure 5 Alignment of 5 AlphaFold3 models of GWD1**

Structural alignment of 5 AlphaFold3 models. **(A)** Models colored by pLDDT where blue colors indicate higher confidence, light blue moderate confidence, and red low confidence in the 3-D placement of the atoms. **(B)** Alignment recolored to show the relative positions of the CBMS. CBM1 is shown in green and CBM2 is shown in blue.

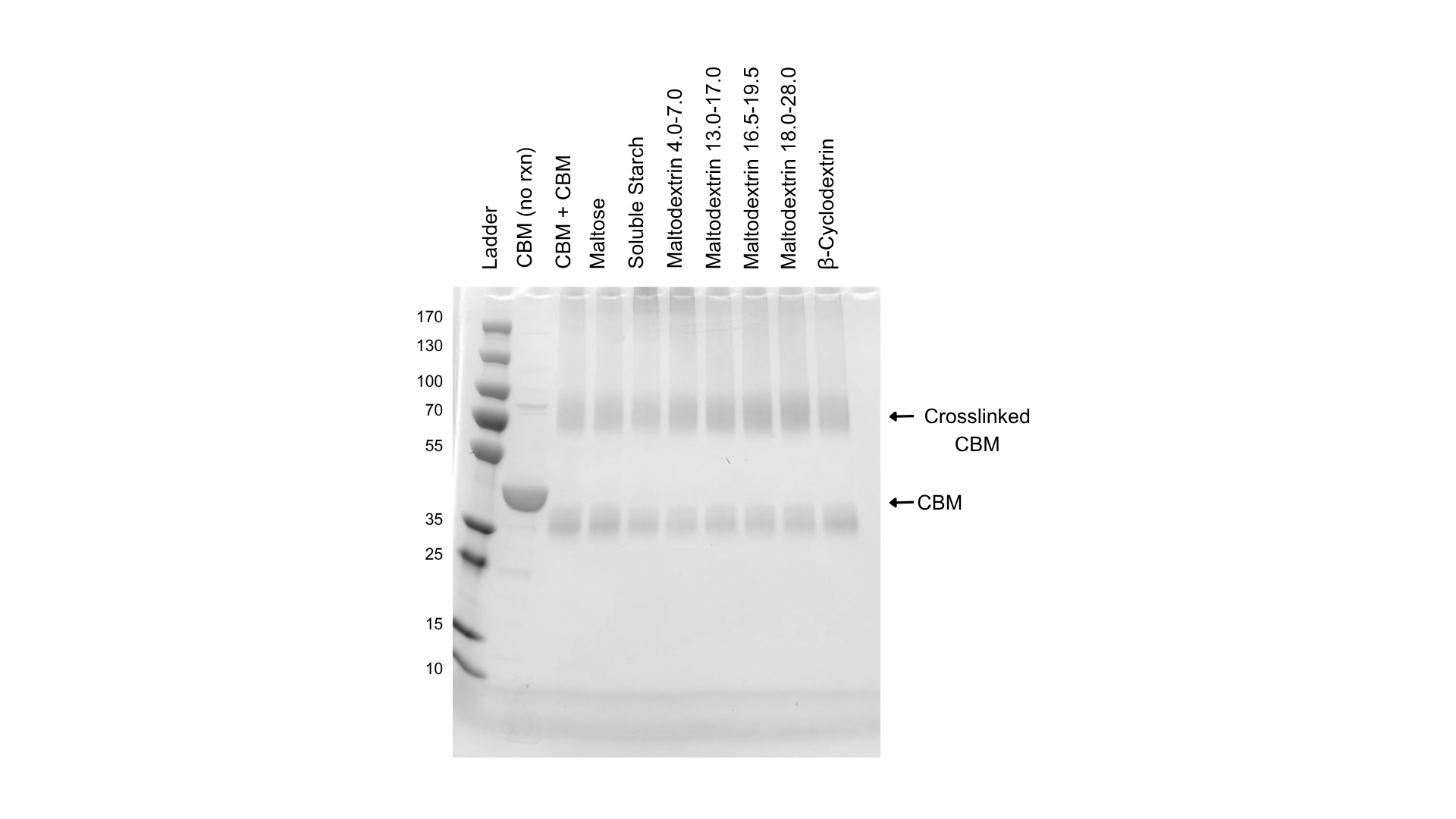

**Supplemental Figure 6: Presence of sugars does not affect CBM crosslinking.**

Glutaraldehyde crosslinking of the CBM in the absence or presence of 0.015 mg/mL of the indicated sugars.

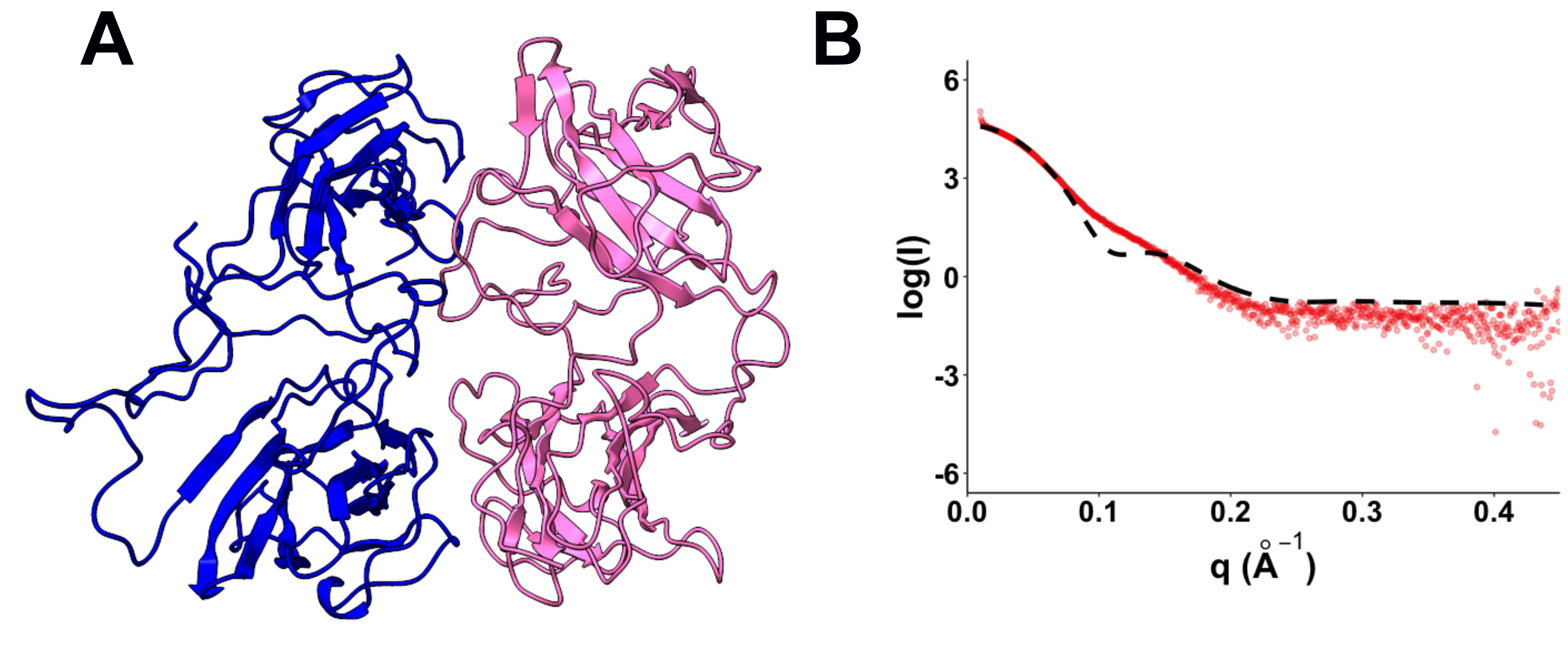

**Supplemental Figure 7: ESMFold model of CBM does not fit well to the SAXS data**

**(A)** BilboMD refined model of the ESMFold generated dimer of the CBM. **(B)** Fit of the BilboMD refined model to the SAXS data for CBM. The X^2^ value for this fit is 13.
